# EPILEPTOGENESIS INHIBITS THE CIRCADIAN CLOCK AND RESHAPES THE DIURNAL TRANSCRIPTOMIC RHYTHMICITY IN THE MOUSE HIPPOCAMPUS

**DOI:** 10.1101/2024.07.02.601732

**Authors:** Radharani Benvenutti, Danielle C. F. Bruno, Matheus Gallas-Lopes, Morten T. Venø, Estela Maria Bruxel, Tammy Strickland, Arielle Ramsook, Aditi Wadgaonkar, Yiyue Jiang, Amaya Sanz-Rodriguez, Lasse Sinkkonen, Marina K.M. Alvim, Clarissa L. Yasuda, Fabio Rogerio, Fernando Cendes, David C. Henshall, Annie M. Curtis, Katja Kobow, Iscia Lopes-Cendes, Cristina R. Reschke

**Affiliations:** School of Pharmacy and Biomolecular Sciences, RCSI University of Medicine and Health Sciences, D02 YN77, Dublin, Ireland; FutureNeuro SFI Research Centre, RCSI University of Medicine and Health Sciences, D02 YN77, Dublin, Ireland; Department of Life Sciences and Medicine, Faculty of Science, Technology and Medicine, University of Luxembourg, 4367, Esch-Belval Esch-sur-Alzette, Luxembourg; Department of Translational Medicine, School of Medical Sciences, University of Campinas (UNICAMP), 13083-888, Campinas, Brazil; Brazilian Institute of Neuroscience and Neurotechnology (BRAINN), Brazil; Pharmacology Department, Federal University of Rio Grande do Sul, 90035-003, Porto Alegre, Brazil; Omiics ApS, Aarhus, Denmark; Department of Physiology and Medical Physics, RCSI University of Medicine and Health Sciences, D02 YN77, Dublin, Ireland; Department of Neurology, School of Medical Sciences, University of Campinas (UNICAMP), 13083-888, Campinas, Brazil; Department of Pathology, School of Medical Sciences, University of Campinas (UNICAMP), 13083-888, Campinas, Brazil; Tissue Engineering Research Group (TERG), RCSI University of Medicine and Health Sciences, D02 YN77, Dublin, Ireland; Department of Neuropathology, Universitätsklinikum Erlangen, Friedrich-Alexander University Erlangen-Nuremberg (FAU), Erlangen, Germany

## Abstract

Epileptogenesis is the process that leads the brain into epileptic activity. Clinical evidence shows that ∼90% of people with epilepsy present rhythmicity in the timing of their seizures presentation. However, whether the circadian clock is a key player during epileptogenesis remains unknown. Here, we triggered epileptogenesis in mice by the intra-amygdala injection of kainic acid and profiled by RNA sequencing their hippocampal diurnal mRNA rhythmicity. We show that epileptogenesis largely reshapes the hippocampal transcriptomic rhythmicity and that the molecular clock machinery is inhibited due to the disruption of the core clock gene *Bmal1*. We identified relevant dysregulated pathways and their dynamics in epileptogenesis, predicting a key role for microglial-driven neuroinflammation. We predicted the genes that *Bmal1* is directly controlling over time. Finally, we sought for translational relevance evidence by performing RNA sequencing in hippocampal samples resected from patients with mesial temporal lobe epilepsy with hippocampal sclerosis (mTLE-HS) and cross-analyzing datasets.

## INTRODUCTION

There are many possible causes of epilepsy, ranging from pathogenic genetic variants to developmental, neoplastic, or acquired structural brain lesions, among others^1–3^. This complex disease presentation hinders understanding the primary biological mechanisms driving epileptogenesis and spontaneous seizures occurrence. It has been well established that both the etiology and the epileptogenic process broadly impact gene expression^4,5^. In turn, changes in gene expression modify brain cells’ and network function triggering further changes in the epileptogenic landscape^6^. More than 80% of the protein-coding genes are known to rhythmically oscillate approximately every 24 h, controlled by the circadian clock^7^.

These circadian-driven transcriptomic changes seem to be particularly prominent in the hippocampus^8^, a brain region that is the seizure onset zone in common types of drug-resistant epilepsies, such as mesial temporal lobe epilepsy (mTLE)^9^. Clinical evidence shows that more than 90% of people with epilepsy present circadian or multidien rhythmicity in their seizure occurrence^10,11^. However, this seems to be highly specific to each person, and in spite of pre-clinical evidence^8^, the underlying molecular features are not fully known. Furthermore, no studies have mapped the molecular oscillatory profile during epileptogenesis, which may lead to these circadian-driven patterns. Defining the impact of epileptogenesis on the diurnal transcriptomic profile is critical for enhancing our understanding of the early events that will lead to the epileptic phenotype.

Circadian rhythmicity is created by an autonomous and intrinsic timekeeping system that regulates the body’s physiological and behavioral changes with an approximately 24 h periodicity^12^. In mammals, a master clock located in the hypothalamic suprachiasmatic nucleus (SCN) acts as the pacemaker synchronizing all the peripheral clock networks across the brain and other tissues^13^. At the cellular level, a ∼24 h long transcriptional translational feedback loop (TTFL), orchestrated by core clock genes, drives the circadian rhythmicity^14^. The heterodimeric transcriptional activators, brain and muscle arnt-like (BMAL1) and circadian locomotor output cycles kaput (CLOCK), bind in the E-box region and increase the transcription of their own inhibitors period (*PER*) and cryptochrome (*CRY*)^12,15,16^. On the other hand, the nuclear receptor subfamily 1 group D members (NR1D1 and NR1D2) and RAR-related orphan receptors (RORA, RORB, and RORC) act as regulators of *BMAL1* expression, through the transcriptional repression or activation, respectively^17,18^. The core clock gene *BMAL1* plays a pivotal role in regulating the clock genes’ rhythmicity in mammals^16^. BMAL1 regulates the expression of many genes and network downstream output pathways, such as sleep-wake cycle, metabolism, cell growth, immunity, and neuronal, astrocyte, and microglia activity^14,19^. The deletion of *Bmal1* in mice, but not other clock genes, is associated with the circadian behavior pattern abolishment^20^. These central clock networks can be synchronized by environmental temporal cues, such as light, also known as *Zeitgeber* time (ZT; the German word for ‘time giver’)^21,22^. Pre-clinical and clinical evidence suggests that the abnormal expression of *BMAL1* may be involved in the pathogenesis of epilepsy^23–25^.

Here, we triggered epileptogenesis in mice by the intra-amygdala microinjection of kainic acid (i.a.KA), a well-established model that leads to treatment-resistant mTLE ^26–28^, and profiled the hippocampal diurnal gene expression during early epileptogenesis by RNA sequencing (RNAseq). We showed that epileptogenesis triggers significant changes in the physiological transcriptomic oscillatory profile and inhibition of the core molecular clock pathway in the mouse hippocampus. We also sought for indication of the translational potential of the identified targets by performing RNAseq in hippocampal samples resected from mTLE patients with hippocampal sclerosis (mTLE-HS). Our findings reveal new aspects to the epileptogenesis pathophysiology, potential novel therapeutic targets, and highlight the relevance of diurnal transcriptomic dynamics.

## RESULTS

### Epileptogenesis triggers significant changes in gene expression

We used RNAseq to examine diurnal molecular dynamics in the hippocampus during early epileptogenesis. Adult male C57BL/6J mice received i.a.KA injections (n=30) to induce status epilepticus (SE) or i.a.PBS (control group, n=30) at 8 am. Ipsilateral hippocampi were collected every 4 h over 24 h at six ZTs (ZT0, ZT4, ZT8, ZT12, ZT16, and ZT20; n=5 per treatment group and ZT; **Fig. 1a**) starting at 24 h after injections when lights were switched on. None of the mice exhibited seizures at the time of tissue collection.

**Figure 1.**
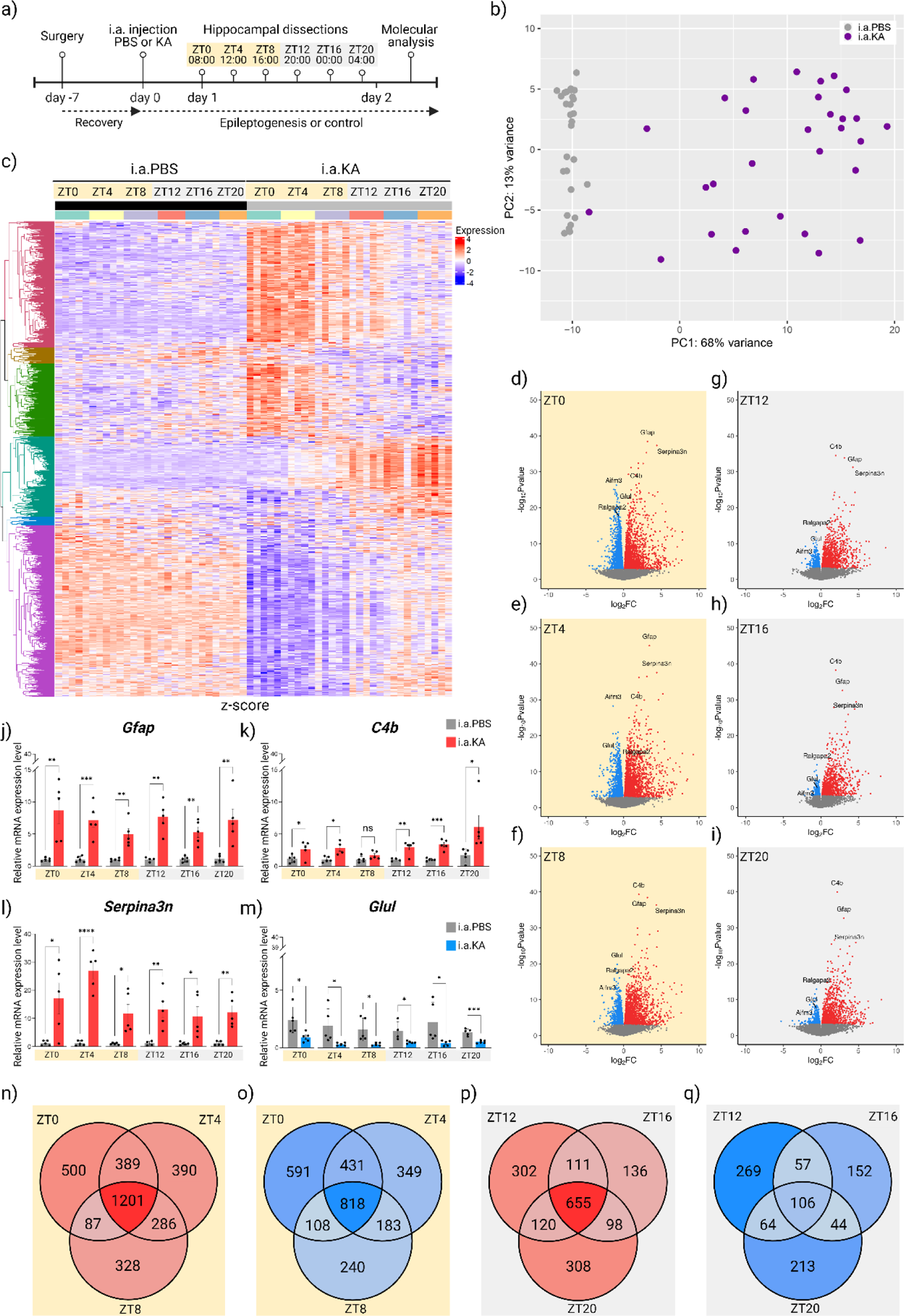
Epileptogenesis largely disrupts gene expression pattern in the mouse hippocampus. **(a)** Experimental design of the diurnal molecular analysis during epileptogenesis (created with BioRender) depicts the multiple hippocampal resections collected at six different ZTs starting at 24 h after intra-amygdala (i.a.) administrations of PBS (phosphate buffered saline) or KA (kainic acid). **(b)** Principal Component (PCA) Analysis showing i.a.KA samples spread pattern on the primary principal component (PC1, 68% variance), while i.a.PBS samples were tightly grouped on the second principal component (PC2, 13% variance). **(c)** RNA sequencing analysis during epileptogenesis in hippocampal samples collected at six different ZTs starting at 24 h after i.a. injections of PBS or KA. **(d-i)** Genes showing significantly differential temporal expression patterns between i.a.PBS and i.a.KA animals (adj p<0.05) are shown with relative high or low expression levels indicated in red and blue, respectively; Volcano plots of differential expression at specific ZTs with the i.a.KA vs i.a.PBS log_2_FC gene expression and statistical significance (−log_10_ p-value) during epileptogenesis in hippocampal samples collected at **(d)** ZT0, **(e)** ZT4, **(f)** ZT8, **(g)** ZT12, **(h)** ZT16 and **(i)** ZT20. Individual genes showing up- or down-regulation are indicated in red and blue, respectively (adj p<0.01). Yellow background represents the light phase and grey background represents the dark phase. **(j-m)** The top 3 upregulated genes and the top 1 downregulated gene in the RNAseq analysis were validated by qPCR in the 6 ZTs. KA increased the relative gene expression (2−ΔΔCT) of **(j)** *Gfap*, **(k)** *C4b*, and **(l)** *Serpina3n* and decreased the gene expression of **(m)** *Glul* at all the ZTs. Multiple Student’s t-tests ± SEM. *p <0.05; **p<0.01; ***p<0.001; ****p<0.0001. **(n-q)** Venn diagrams showing the **(n)** upregulated genes in each ZT during the light phase, **(o)** downregulated genes in each ZT during the light phase, **(p)** upregulated genes in each ZT during the dark phase, **(q)** downregulated genes in each ZT during the dark phase. In the Venn diagrams, color intensity strictly correlates with the number of altered genes within each individual diagram. A total of 59 hippocampal samples were used, n=5 mice per ZT/group (except for i.a.PBS at ZT12, where n=4). i.a.KA, intra-amygdala kainic acid; i.a.PBS, intra-amygdala PBS; ZT, *Zeitgeber* time; ZT0, 8 am; ZT4, 12 pm; ZT8, 4 pm; ZT12, 8 pm; ZT16, 12 am; ZT20, 4 am.

Principal Component Analysis (PCA) showed clear dimension separation between the i.a.KA and control samples (**Fig. 1b**), with the former displaying greater variability. RNAseq identified more than 50,000 transcripts mapping to 6,435 differentially expressed genes (DEGs) between the i.a.KA and control groups (**Fig. 1c**, DESeq2<0.05). Hierarchical clustering revealed six primary gene clusters (**Fig. 1c** and **Supplementary Data 1**). Significant DEGs, consistent across all time points during epileptogenesis, included upregulated genes such as glial fibrillary acidic protein (*Gfap*), serine peptidase inhibitor a3n (*Serpina3n*), and complement component 4B (*C4b*), and downregulated glutamate-ammonia ligase (*Glul*), as confirmed by qPCR (**Fig. 1j-m**). For detailed information on the top 20 most significantly up- or downregulated genes (adj p≤0.01) per time point and their biological functions, refer to **Supplementary Tables 1** and **2**.

### Differences in gene expression in epileptogenesis are exacerbated in the light phase

Gene expression during early epileptogenesis showed a clear time of day pattern, with greater differences during the light phase (ZT0, ZT4, ZT8) compared to the dark phase (ZT12, ZT16, ZT20). During the light phase, 5,901 genes were differentially expressed (3,181 upregulated, 2,720 downregulated) when comparing epileptogenic to control mice (**Fig. 1d-f** and **Fig. 1n-o**). In contrast, during the dark phase, the expression of 2,635 genes showed significant changes (1,730 upregulated, 905 downregulated; **Fig. 1g-i**, and **Fig. 1p-q**). Our findings highlight that the expression profile of specific genes can vary within the same phase, underlying the importance of considering the time of day in sampling and transcriptomic studies.

We identified the top 500 most significant DEGs (adj p<0.05) and conducted pathway enrichment analysis using DAVID^29^. For control (i.a.PBS) samples, Gene Ontology (GO) Biological Process terms related to circadian rhythm were the top results (**Supplementary Fig. S2a**), affirming the influence of circadian oscillations in hippocampal function. Conversely, in the i.a.KA samples, circadian rhythm-related terms were absent from the top 10 GO terms (**Supplementary Fig. S2b**). Instead, terms related to cell division and cell cycle dominated, suggesting a disruption in diurnal gene expression oscillation during early epileptogenesis. Overall, our findings indicate that epileptogenesis significantly alters gene expression patterns, particularly during the light phase.

### A unique diurnal molecular oscillatory profile is triggered by epileptogenesis in mice

To determine if the changes in gene expression during early epileptogenesis were rhythmic, we used the MetaCycle R package, incorporating ARSER, JTK_CYCLE, and Lomb-Scargle methods. Genes having an adj p-value (meta2d_BH.Q) <0.05 were considered significantly rhythmic. Under a conservative threshold (adj p<0.001), we found that 1,489 genes displayed ∼24 h oscillations in control mice (i.a.PBS group), while 161 genes were rhythmic in the i.a.KA epileptogenic group (**Fig. 2a** and **Supplementary Data 2a-b**). Out of these, only 21 genes maintained diurnal rhythms across both groups (**Fig. 2a** and **Supplementary Data 2c**). This indicates a loss of rhythmicity for 1,468 genes during epileptogenesis, while 140 genes newly acquired rhythms. The biological functions of the top 100 genes that lost or gained rhythmicity and the 21 genes that maintained diurnal rhythms are detailed in **Supplementary Tables 3-5**. Interestingly, 19 out of the 21 genes that kept cycling in both groups showed an inversion in their rhythmic phase. Only two genes - Kruppel-like transcription factor 15 (*Klf15*) and patatin-like phospholipase domain containing 7 (*Pnpla7*) – retained their original oscillatory profiles (Pearson correlation coefficient range was −1 to 1; **Supplementary Table 5**). *Klf15* is involved in metabolic signaling, and *Pnpla7* is linked to lysophosphatidylcholine (LPC) hydrolysis in the endoplasmic reticulum. The biological functions of the 19 genes (*Slc25a18, Myorg, Ptprz1, Prrg1, Gja1, Nkain4, Phgdh, Ucp3, S1pr1, Carmil1, Hmgcll1, Acsl3, Cdh12, Cdh20, Utp14b, Map7, Selenop, Gpt2* and *Cyp2j9*) inverting their oscillatory phases during early epileptogenesis are listed in **Supplementary Table 5**. Notably, genes that altered their rhythmicity in epileptogenesis might also be transiently or consistently differentially expressed over ∼24 h, indicating complex regulatory changes in response to epileptogenic conditions.

**Figure 2.**
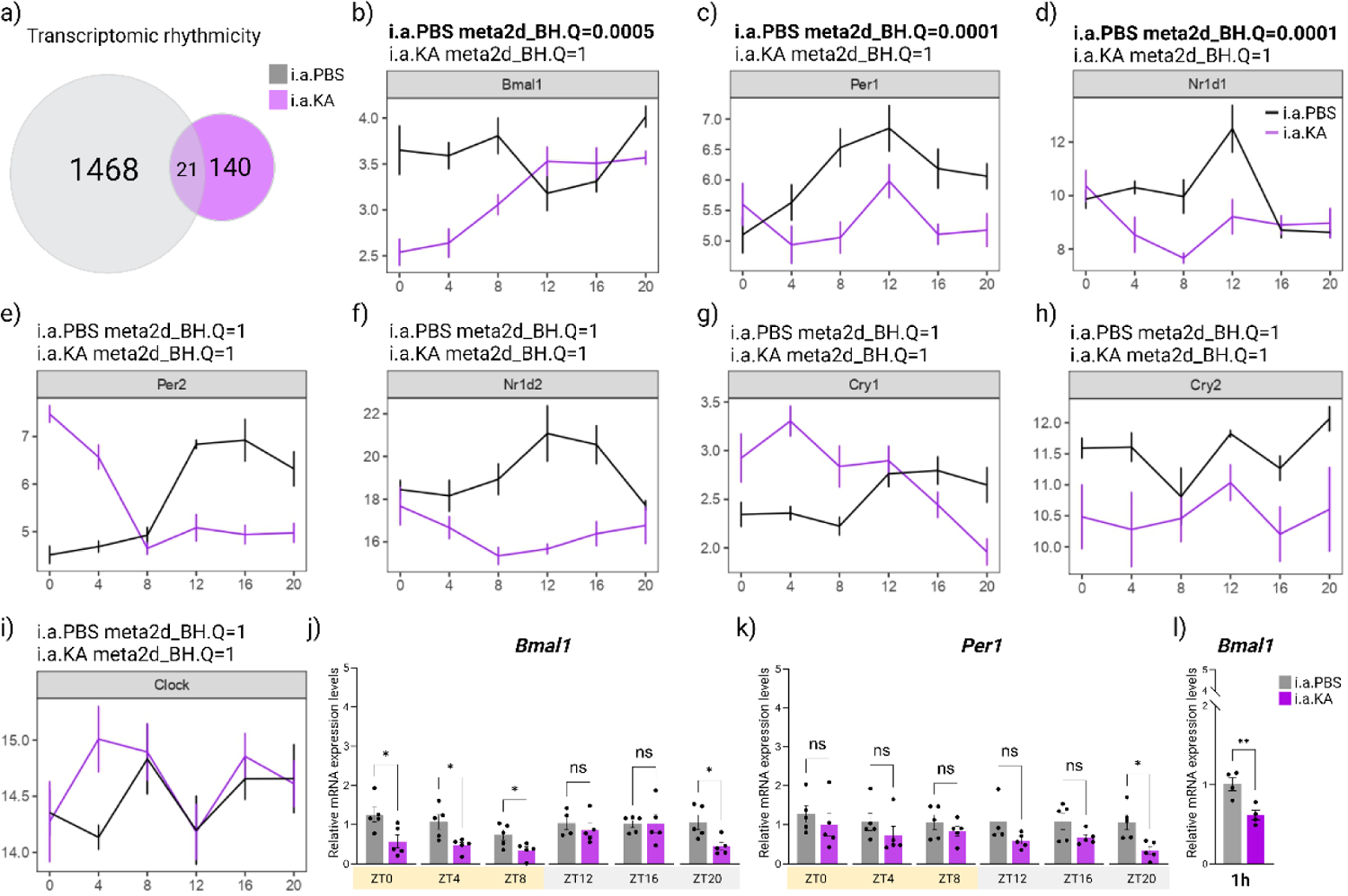
Diurnal mRNA rhythmicity disruption during epileptogenesis also impacts the key core clock components. **(a)** Venn diagrams show the number of genes presenting significant rhythmicity in hippocampus of i.a.PBS and/or i.a.KA groups. **(b-i)** Rhythmicity analysis of i.a.PBS (black) and i.a.KA (purple) of the core clock genes **(b)** *Bmal1*, **(c)** *Per1,* **(d)** *Nr1d1,* **(e)** Per2, **(f)** *Nr1d2*, **(g)** *Cry1,* **(h)** Cry2, and **(i)** *Clock*. The X-axis represents each ZT, while the Y-axis indicates the intensity of transcript counts. The rhythmicity analysis showed that the main core clock genes *Bmal1, Per1*, and *Nr1d1* lost their rhythmicity during epileptogenesis (i.a.PBS meta2d_BH.Q-value=0.0005, 0.0001 and 0.0001, respectively, while all presented i.a.KA meta2d_BH.Q-value=1). The other core clock genes did not present rhythmicity in both groups (i.a.PBS meta2d_BH.Q-value=1 and i.a.KA meta2d_BH.Q-value=1). **(j)** *Bmal1* and **(k)** *Per1* relative gene expression (2−ΔΔCT) was also validated by qPCR in the 6 ZTs. Multiple Student’s t-tests ± SEM, *p<0.05. A total of 59 hippocampal samples were used, n=5 per group (except for i.a.PBS at ZT12, where n=4). **(l)** *Bmal1* relative gene expression (2−ΔΔCT) assessed by qPCR analysis 1h after i.a. injection of PBS or KA. Student’s t-test ± SEM, **p<0.01. n=4 per group. i.a.KA, intra-amygdala kainic acid; i.a.PBS, intra-amygdala PBS; ZT, *Zeitgeber* time; ZT0, 8 am; ZT4, 12 pm; ZT8, 4 pm; ZT12, 8 pm; ZT16, 12 am; ZT20, 4 am.

### The core clock transcriptional machinery is inactive in the hippocampus during epileptogenesis

We further examined the diurnal rhythmicity of the primary core clock genes *Bmal1, Per1, Nr1d1, Per 2, Nr1d2, Cry1, Cry2,* and *Clock* (**Fig. 2b-i**, respectively). Among the 1,468 genes that lost their rhythmicity during epileptogenesis, we found *Bmal1*, *Per1,* and *Nr1d1*. These genes had significant rhythmicity in controls (i.a.PBS meta2d_BH.Q-value=0.0005, 0.0001, and 0.0001, respectively), but this was absent in epileptogenic mice (i.a.KA meta2d_BH.Q-value=1; **Fig. 2b-d**).

To validate these findings, we used qPCR to measure the expression of *Bmal1* and *Per1*, TTFL positive and negative regulators, respectively, at the six ZTs (**Fig. 2j-k**). The levels of *Bmal1* were significantly reduced at ZT0 (p=0.0271), ZT4 (p=0.0172), ZT8 (p=0.0367), and ZT20 (p=0.0175) in the i.a.KA group, with no differences at ZT12 and ZT16 (p>0.05, **Fig. 2j**), aligning with our initial observations (**Supplementary Fig. S1**). In fact, *Bmal1* was downregulated in the i.a.KA group as early as 1 h after status epilepticus induction when compared to time-matching controls (p=0.0085, **Fig. 2l**). Meanwhile, *Per1* levels showed a significant decrease in the epileptogenic samples only at ZT20 (p=0.0101; **Fig. 2k**). Additionally, *Bmal2*, a paralog of *Bmal1*, was downregulated in the i.a.KA group exclusively at ZT0 (p=0.0291; **Supplementary Fig. S3**).

To understand the broader implications of these disruptions, we conducted *in silico* analyses using the Ingenuity Pathway Analysis (IPA). The analyses predicted widespread inhibition of the core circadian regulator pathway in the hippocampus during epileptogenesis across all ZTs (**Fig. 3** and **Supplementary Fig. S4-8)**. This involved the inhibition of key components such as *Clock*, *Per1*, *Cry1*, *Cry2,* largely due to the transient downregulation of *Bmal1*. This inhibition extends to other crucial TTFL molecules, including *Nr1d1* and *Nr1d2* (**Fig. 3** and **Supplementary Fig. S4-8**). Moreover, the transcription factors *Rora* and *Rorc* were also predicted to be less active in the i.a.KA group (**Fig. 3** and **Supplementary Fig. S4-8**). These findings collectively underscore the significant impact of epileptogenesis on the core clock machinery, leading to its inactivation in the mouse hippocampus.

**Figure 3.**
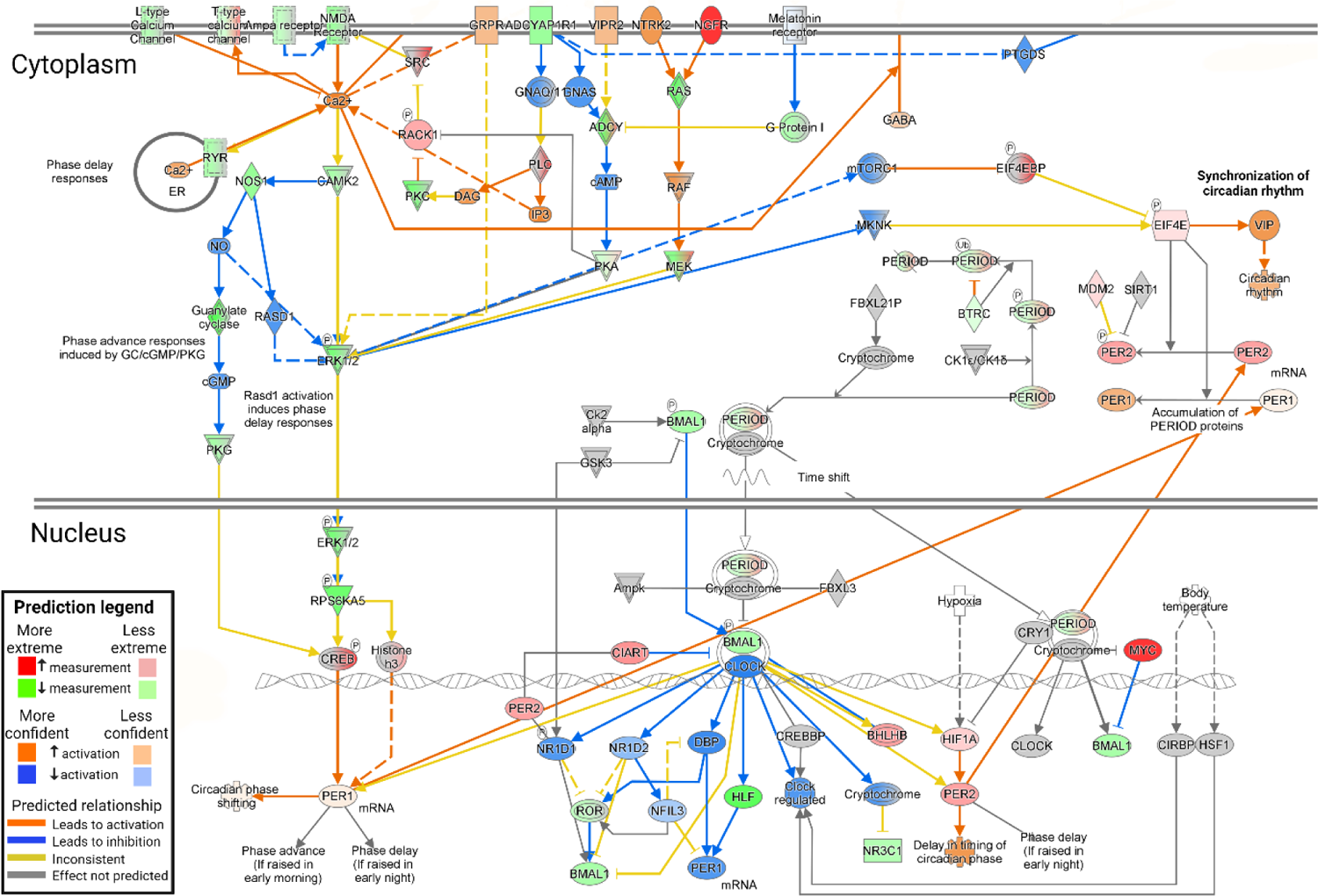
The circadian molecular clock is inhibited in the hippocampus during epileptogenesis. The schematic depicts the circadian molecular machinery in the signaling pathway in the hippocampus at ZT0 (8 am, when lights were turned on at 24 h after i.a. injections of PBS or KA). Predictions of activation/inhibition of the pathway were identified by IPA analysis of the RNAseq dataset. The intensity of colors (green and red) corresponds to the level of log_2_FC; weaker color indicates a lower expression level, while stronger color indicates a higher expression. Similarly, activation/inhibition of molecule activity, as defined by the activation z-score (orange lines) and inhibition z-score (blue lines), is depicted by color intensity, in which weaker color represents lower activation/inhibition level, whereas stronger color indicates higher activation/inhibition. Targets and/or molecular interactions marked in yellow lines represent inconsistency according to the literature; white represents molecules involved in the signaling pathway but absent in our dataset; gray represents that the genes and/or interactions are present in our dataset but the algorithm was unable to predict their contribution to the pathway activation/inhibition. Sharp arrows represent stimulation; blunt arrows represent inhibition; dashed line indicates indirect interaction. See **Supplementary Fig. S22** for extra legend information. This figure was simplified from the IPA to focus only on the intracellular processes.

### Dynamic microglial-mediated neuroinflammation is predicted as a key regulator of the early epileptogenic process

To explore how differential gene expression dynamics contribute to epilepsy development, we identified the most significant biological processes across the six ZTs using the Reactome database. Immune system-related pathways dominated the top 20 enriched pathways by False Discovery Rate (FDR) value (**Fig. 4a**). Comparative IPA core analysis at each ZT revealed that ‘central nervous system inflammation’ was the top disease/dysfunction active across all ZTs. Other inflammatory processes were similarly active throughout (**Fig. 4b**). Notably, terms related to seizure disorders were active only during the light phase (ZT0, ZT4, and ZT8; **Fig. 4b**). The terms ‘epilepsy’, ‘epileptic seizures’, and ‘generalized seizures’ were active only when lights were turned on (ZT0) (**Fig. 4b**).

**Figure 4.**
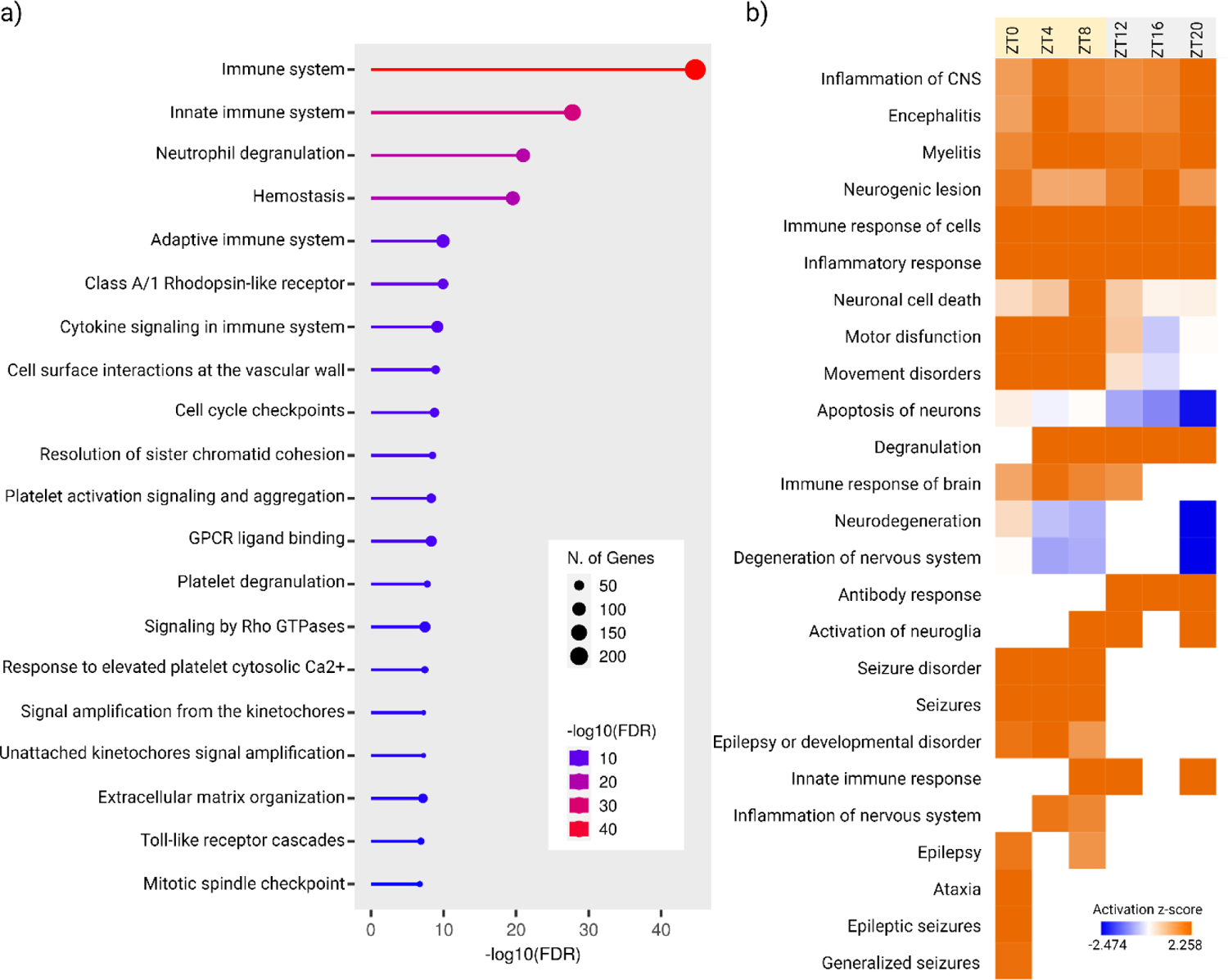
Neuroinflammation is the main process in the hippocampus during epileptogenesis. **(a)** Reactome pathway analysis was conducted using ShinyGO to identify the top 20 significantly enriched pathways across the 6 ZTs. Pathways are ranked on the y-axis based on FDR enrichment, with colors representing -log(FDR) values. Dot size reflects the number of genes associated with each pathway. **(b)** Top disease and biological processes activated/inhibited using comparative IPA core analysis from differentially expressed coding genes (adj p<0.01) for each 6 ZTs. Data filtering was applied specifically to neurological diseases, inflammatory responses, and inflammatory diseases, focusing on those demonstrating significant activation/inhibition processes with an absolute z-score above 2, and -log_10_ p-value and -log_10_BH-adjp≥1.3, respectively. The top disease/dysfunction identified was ‘inflammation of the central nervous system (CNS)’, followed by other four inflammatory-related hits, which were active across the six ZTs. Seizure disorder’, ‘seizures’, ‘epilepsy or developmental disorder’ were found as active processes only during the light phase (ZT0, ZT4, and ZT8), while ‘epilepsy’, ‘epileptic seizures’ and ‘generalized seizures’ were also identified in our list as processes active only when lights are turned on (ZT0). The heatmap shows the z-scores of each entity in each analysis. Orange indicates the activation and blue the inhibition predicted of the diseases and/or functions. CNS, central nervous system; i.a.KA, intra-amygdala kainic acid; i.a.PBS, intra-amygdala PBS; ZT, Zeitgeber time; ZT0, 8 am; ZT4, 12 pm; ZT8, 4 pm; ZT12, 8 pm; ZT16, 12 am; ZT20, 4 am.

Enriched canonical pathways from core IPA analysis revealed that neuroinflammation signaling pathways were predominantly active in the hippocampus during epileptogenesis (**Fig. 5** and **Supplementary Fig. S9-13**). Microglial activation was predicted as the main driver of these neuroinflammatory responses, with astrocytes and neurons contributing modestly (**Fig. 5** and **Supplementary Fig. S9-13**). Key regulators included CX3C chemokine receptor 1 *(Cx3cr1)*, interferons (*If-α*, *If-β*, and *If-γ*, respectively), mitogen-activated protein kinase (*Mapk),* nuclear factor kappa-light-chain-enhancer of activated B cells *(Nfkb),* and Toll-like receptors (*Tlr2* and *Tlr4*). These regulators exhibited dynamic profiles across different ZTs (**Fig. 5**, **Supplementary Fig. S9-13**, and **Supplementary Table 6**).

**Figure 5.**
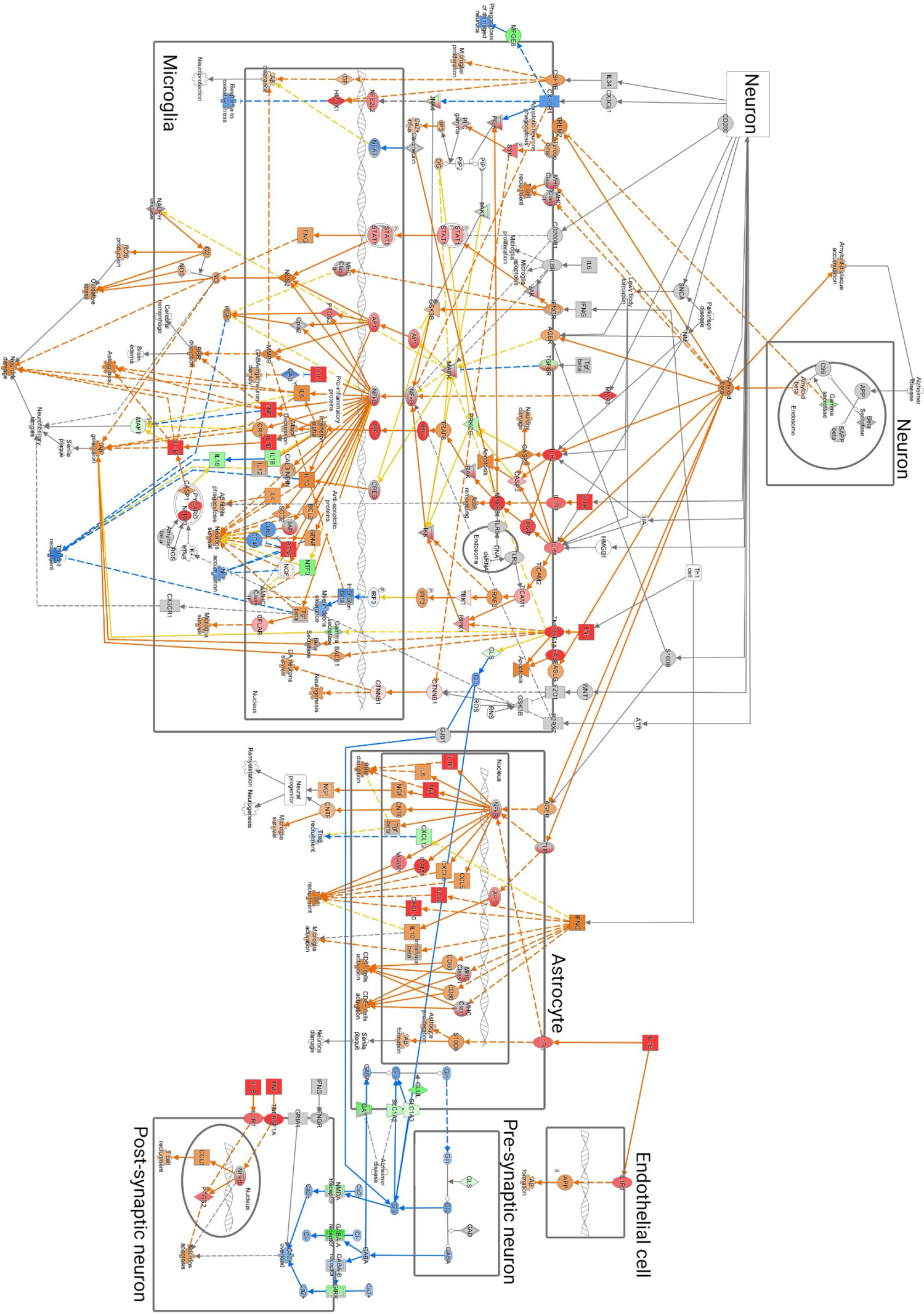
The IPA canonical pathway core analysis revealed dynamic microglial-mediated neuroinflammation as a key regulator of epileptogenesis. Log_2_FC and adjusted p-values of differentially expressed coding genes, as well as Z-scores generated by IPA core analysis at ZT0, were illustrated for the neuroinflammation signaling pathway. Distinct colors (downregulated in green and upregulated in red) determine overlap and common relationships between genes and components of the neuroinflammation signaling pathway. The intensity of the green and red indicates the degree of DEG levels. Orange was predicted as active and blue as inhibited. Yellow arrows highlight the main dysregulated interactions. Molecules in gray are present in the ZT0 DEGs dataset, but could not present statistical significance to predict their activation or inhibition process in the pathway. White molecules are part of the pathway but are not present at ZT0. Sharp arrows represent stimulation; blunt arrows represent inhibition; dashed line indicates indirect interaction. See **Supplementary Fig. S22** for extra legend information.

Specifically, *Cx3cr1,* activated by *Cx3cl1* produced by neurons in response to inflammation, was inactive at ZT0 when the light phase started (**Fig. 5**), but became upregulated from ZT4 to ZT20 (**Supplementary Fig. S9-13**). This progressive activation stimulates the mitogen-activated protein kinase 8 (*Jnk*) associated with T cell proliferation, apoptosis, and differentiation. JNK indirectly regulates nuclear factor, erythroid derived 2, like 2 (*Nfe2l2*), and heme oxygenase 1 (*Hmox1*), both involved in antioxidant response and neuroinflammation, which were upregulated across ZTs. *Cx3cr1* activation likely also influenced the downregulation of milk fat globule EGF and factor V/VIII domain containing (*Mfge8*), involved in apoptotic cell clearance.

*Ifn-γ,* a potent activator of glial cells and major histocompatibility complex (MHC) molecules was active across all ZTs (**Fig. 5** and **Supplementary** Fig. 9-12), except ZT20 (**Supplementary Fig. S13**). In our analysis, *Ifn-γ* activated *Mhc-2*, but also signal transducer and activator of transcription 1 (*Stat1)* and glycogen synthase kinase 3 beta (*Gsk3b),* both involved in apoptotic processes and upregulated at all ZTs.

*Mapk,* a regulator of essential cellular functions such as proliferation, differentiation, and apoptosis, showed erratic rhythms, being upregulated only at ZT0 (**Fig. 5**) and active at ZT20 (**Supplementary Fig. S13**), with downregulation at ZT4, ZT8, and ZT12 (**Supplementary Fig. S9-11**) and inactivation at ZT16 (**Supplementary Fig. S12**). *Mapk* influenced activator protein-1 (*Ap1*) and cAMP-responsive element binding protein 1 (*Creb*), which both showed dynamic profiles (**Supplementary Fig. S12**) and activated nitric oxide synthase 2 (*Nos2*), expressed in neurons in the healthy brain, and in microglia upon inflammation, and interleukin 10 *(Il-10)* respectively, an anti-inflammatory cytokine. Despite *Mapk*’s variable influence, *Nfκb*, remained consistently upregulated at all ZTs in microglia and astrocytes, driving the expression of pro-inflammatory cytokines interleukin 4 (*Il-4*), interleukin 6 (*Il-6*), interleukin 1*β* (*Il-1β),* interleukin 12 (*Il-12),* and *Tnf* across all ZTs (**Fig. 5** and **Supplementary Fig. S9-13**). Interleukin 18 *(Il-18)* showed a unique profile, being downregulated at ZT0, ZT4, and ZT8 (**Fig. 5** and **Supplementary Fig. S9-10**), and activated at ZT12, ZT16, and ZT20 (**Supplementary Fig. S11-13**). Moreover, *Il-18* lost its rhythmicity in epileptogenesis (i.a.KA meta2d_BH.Q=0.02, and meta2d_BH.Q=0.00005 for i.a.PBS).

Lastly, *Tlr2* and *Tlr4* were upregulated during epileptogenesis across all ZTs, predominantly in microglia and astrocytes (**Fig. 5** and **Supplementary Fig. S9-13**). *Tlr4* activation, though not directly linked to high mobility group box-1 (*Hmgb1*) in our analysis, is known to be involved in ictogenesis and epilepsy^30,31^. Our data showed *Hmgb1* losing its rhythmicity (i.a.KA meta2d_BH.Q=1, and meta2d_BH.Q=0.00013 for i.a.PBS) and upregulation at ZT0, ZT4, ZT8, and ZT20 (adj p<0.01). *Tlr4*’s activation led to the upregulation of *Ifn-α*, *Il-6*, *Tnf-α*, and *Myd88,* and the downregulation of *Ifn-β* all the ZT*s* (**Fig. 5** and **Supplementary Fig. S9-13**). *Tlr2* activation, linked to tissue damage, promoted apoptosis at all the ZTs through caspase 8 (*Casp8*) and caspase 3 (*Casp3*), with *Casp3* losing its rhythmicity during epileptogenesis (i.a.KA meta2d_BH.Q=0.6, and meta2d_BH.Q=0.000005 for i.a.PBS). These findings highlight the complex and dynamic role of microglial-mediated neuroinflammation in the early stages of epileptogenesis.

### Astrocytic-neuronal turnover of Glutamate-GABA is dysregulated in epileptogenesis

As previously outlined, a complex crosstalk regulation of neuroinflammation exists between microglia, neurons, and astrocytes (**Fig. 5** and **Supplementary Fig. S9-13**). In astrocytes, Toll-like receptors activate *NFκB* and *Ip1* at all the ZTs. *NFκB*, consistently upregulated, stimulates pro-inflammatory cytokines such as *Il-1β, Il-6,* and *Tnf-α*. *Mhc-I* and *Mhc-II* are also upregulated across all ZTs, facilitating CD4+ and CD8+ cell activation. Conversely, *Cxcl12* shows a dynamic profile, downregulated from ZT0 to ZT16 and active at ZT20 (**Fig. 5** and **Supplementary Fig. S9-13**). In neurons, the neuroinflammatory pathways are less conspicuous with a predicted upregulation of *Tnf-α* and *Il-1β* and activation *NFκB, Ptgs2,* and *Ccl3*.

Our dataset reveals a dynamic dysregulation in glutamate and γ-Aminobutyric acid (GABA) turnover between astrocytes and neurons (**Fig. 5** and **Supplementary Fig. S9-13**). At ZT0, IPA analysis predicted that major components of glutamate and GABA turnover are downregulated or inhibited. In neurons, genes that encode the glutamatergic N-methyl-D-aspartate (NMDA) receptor, γ-Aminobutyric acid type A (GABA-A) receptor are downregulated, while genes that encodes the γ-Aminobutyric acid type B (GABA-B) receptor are inhibited, leading to predicted Ca^2+^ overload and neuronal apoptosis **(Fig. 5)**. G protein-gated inwardly rectifying potassium channel (*Girk)*, which encodes a G protein essential for regulating cellular excitability in the heart and brain^32^, is downregulated.

In astrocytes, genes encoding high-affinity glutamate transporters (solute carrier family members SLC1A3 and SLC1A2) and the GABA transporter (GAT) are also downregulated at ZT0. At ZT12, there is a predicted progressive activation of glutamate production driven by microglial glutaminase (GLS), a mitochondrial enzyme that catalyzes the breakdown of glutamine to form glutamate (**Supplementary Fig. S11**). This consequently results in a progressive activation of the NMDA receptor gene at ZT16 and ZT20 (**Supplementary Fig. S12-13**). The overall dynamic pattern of glutamate and GABA turnover dysregulation suggests the onset of a more excitable cellular state.

### *Bmal1* largely controls neuroinflammation and metabolism during epileptogenesis

To understand the role of *Bmal1* in neuroinflammation during epileptogenesis, we used the upstream regulator IPA core analysis tool to identify its direct targets at different ZTs (see **Supplementary Table 7** for details). *Bmal1* inhibition levels varied across ZTs, with progressive increases from ZT4 to ZT16 (ZT4 z-score=-0.518, p=0.00000499; ZT8 z-score=-1.055, p=0.0000000179; ZT12 z-score=-1.003, p=0.0000374; ZT16 z-score=-1.608; p=0.00000262; ZT20 z-score=0.032, p=0.0000011). While *Bmal1* levels at ZT0 and ZT20 (z-score=0.018, p=0.0000213) were inconclusive, its inhibition was still evident in the circadian rhythm signaling pathway (**Fig. 3** and **Supplementary Fig. S8**).

*Bmal1*’s regulatory influence peaked during the light phase (ZT0 to ZT8; **Fig. 6 a-c**), affecting 38 genes related to inflammation, oxidative stress, metabolism, and cell cycle (**Supplementary Table 7**). During the dark phase (ZT12 to ZT20; **Fig. 6 d-f**), 28 genes were regulated by *Bmal1*, most of them involved in metabolism and transcriptional regulation. The dark phase was less related to inflammatory processes than the light phase. For instance, *Il-1β* and *Nfe2l2* were targeted in the light phase (ZT0 to ZT8) but only at ZT20 in the dark phase.

**Figure 6.**
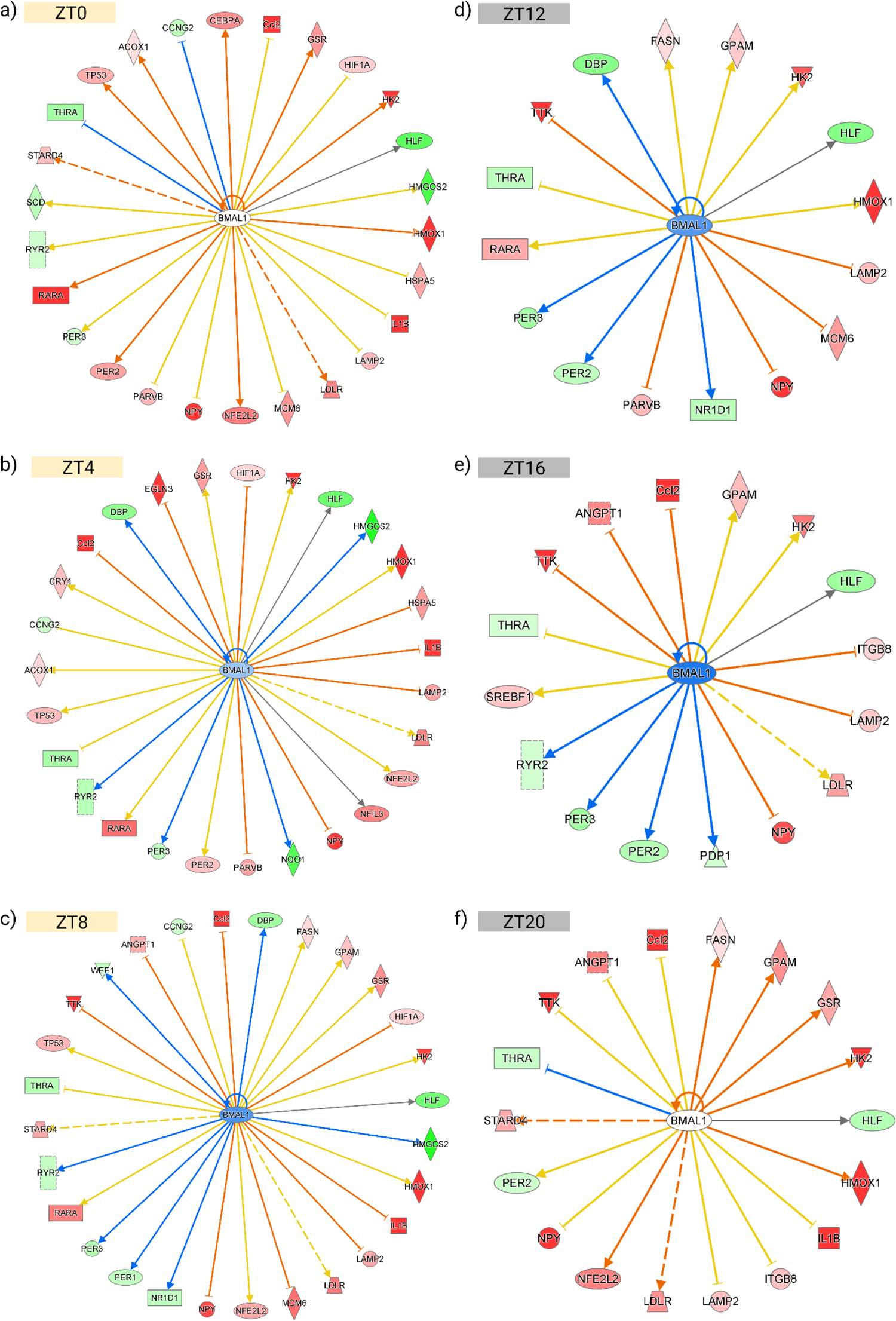
*Bmal1* as an upstream regulator with corresponding activation/inhibition of IPA-predicted target genes during epileptogenesis. The activation/inhibition predicted in this analysis aligns with the RNAseq dataset at **(a)** ZT0, **(b)** ZT4, **(c)** ZT8, **(d)** ZT12, **(e)** ZT16, and **(f)** ZT20. Red indicates upregulated genes, while green signifies downregulated; the intensity of colors (green and red) corresponds to the level of log_2_FC; weaker color indicates a lower expression level, while stronger color indicates a higher expression. Similarly, activation/inhibition of molecule activity, as defined by the activation z-score (orange lines) and inhibition z-score (blue lines), is depicted by color intensity: weaker color represents lower activation/inhibition level, whereas stronger color indicates higher activation/inhibition. Yellow lines represent inconsistency according to the literature; white represents molecules involved in the signaling pathway but absent in our dataset. Sharp arrows represent stimulation; blunt arrows represent inhibition; dashed line indicates indirect interaction. See **Supplementary Fig. S22** for extra legend/symbol information. ZT, *Zeitgeber* time; ZT0, 8 am; ZT4, 12 pm; ZT8, 4 pm; ZT12, 8 pm; ZT16, 12 am; ZT20, 4 am.

Our findings suggest that *Bmal1* dynamically regulates inflammation, metabolism, and cell cycle processes and that its dysregulation disrupts brain homeostasis during epileptogenesis. The full list of *Bmal1* targets per ZT is provided in **Fig.6** and **Supplementary Table 7**.

### Human mTLE-HS transcriptome unveils distinct mechanisms of circadian rhythm and immune system dysregulation

To explore the translational relevance of our findings in humans with mTLE-HS, we conducted RNAseq on hippocampal samples resected from 17 mTLE-HS patients at ∼2 pm (local time; see **Supplementary Table 8** for further information). PCA analysis showed a clear separation between mTLE-HS patients (PC1, 21% variance) and the autopsy controls (PC2, 14% variance), indicating distinct DEGs profiles associated with phenotype (**Supplementary Fig. S14**). RNAseq identified 37,544 transcripts, with 4,271 DEGs (adj p≤0.05) in mTLE-HS samples compared to controls. This corresponds to 3,660 annotated protein-coding genes, which 1,861 are upregulated and 1,799 are downregulated (**Fig. 7a** and **Supplementary Data 2d**).

**Figure 7.**
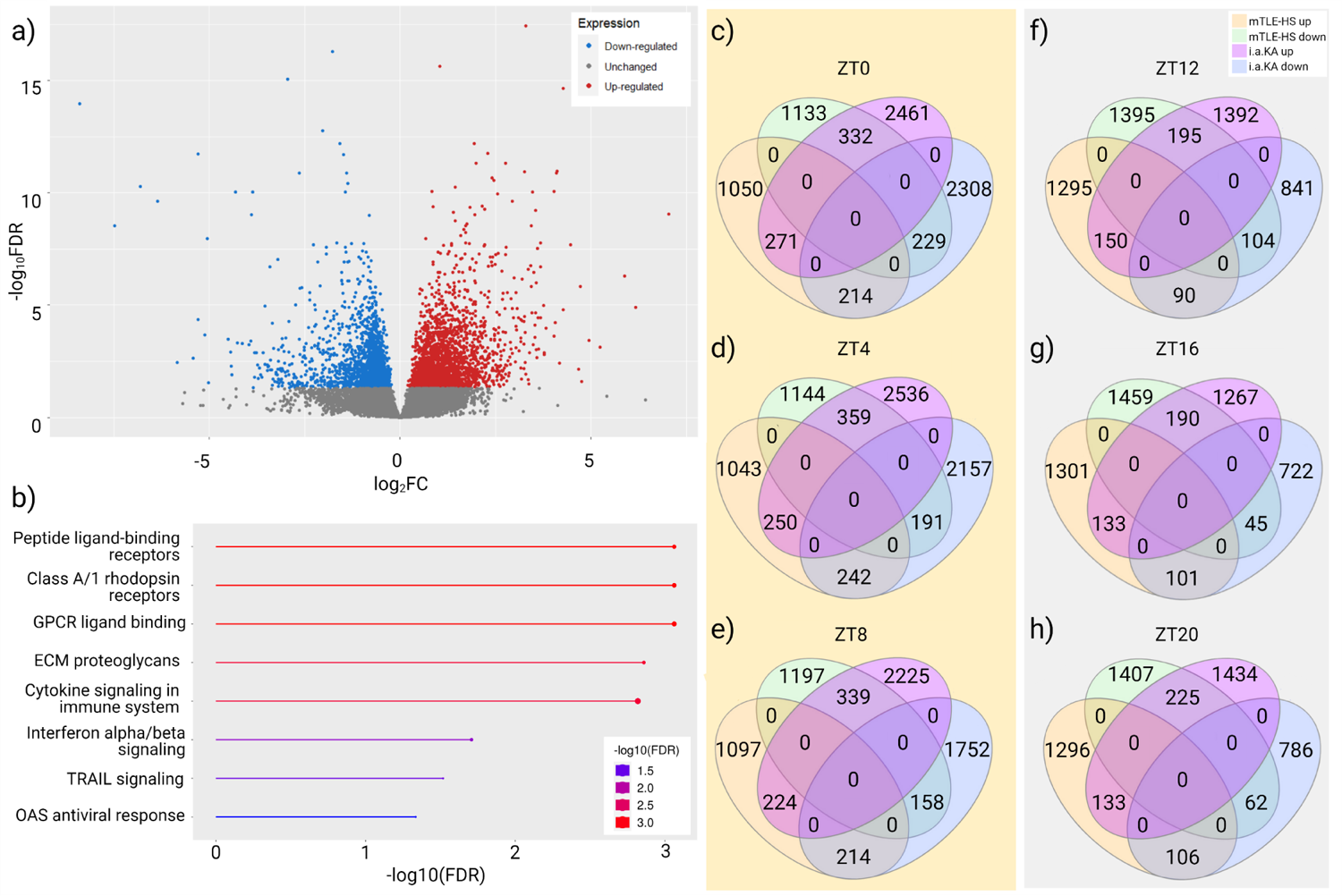
The mTLE-HS patients’ hippocampal transcriptomic analysis resonates with key epileptogenesis findings. **(a)** Volcano plots of differential expression with the hippocampal samples from human mTLE-HS vs control log2FC gene expression and statistical significance (−log10 p-value). Individual genes showing up- or down-regulation are indicated in red and blue, respectively (adj p<0.05). **(b)** Reactome pathway analysis was conducted using ShinyGO to identify the top significantly enriched pathways from the human mTLE-HS RNAseq dataset filtered by log_2_FC <-1 and >1, and adj p<0.01. Pathways are ranked on the y-axis based on FDR enrichment, with colors representing -log_10_FDR values. Dot size reflects the number of genes associated with each pathway. **(c-h)** The Venn diagrams represent the overlaps identified between the up and downregulated genes from the RNAseq in mice and human mTLE-HS for **(c)** ZT0, **(d)** ZT4, **(e)** ZT8, **(f)** ZT12, **(g)** ZT16, and **(h)** ZT20. Yellow background represents the light phase and grey background represents the dark phase. A total of 59 mouse hippocampal samples were used, n=5 per group (except for i.a.PBS at ZT12, where n=4), and 23 dissected hippocampal human samples (17 patients with sporadic mTLE-HS and 6 control individuals). ECM, extracellular matrix; GPCR, G protein– coupled receptors; OAS, 2′-5′ oligoadenylate synthase; TRAIL, TNF-related apoptosis-inducing ligand; ZT, *zeitgeber* time.

Filtering DEGs by log_2_FC ≤ −1 and ≥ 1 yielded 857 upregulated and 461 downregulated genes. Enrichment pathways analysis using the Reactome database identified eight top processes (FDR<0.05): ‘peptide ligand-binding receptor’, ‘class A/1 rhodopsin receptors’, ‘G protein-coupled receptors (GPCR) ligand binding’, ‘extracellular matrix (ECM) proteoglycans’, ‘cytokine signaling and immune system’, ‘interferon alpha, and beta signaling’, ‘TNF-related apoptosis-inducing ligand (TRAIL) signaling’ and ‘2′-5′ oligoadenylate synthase (OAS) antiviral response’ (**Fig. 7b**). KEGG pathway analysis highlighted ‘Cytokine-cytokine receptor interaction’ and ‘TNF signaling pathway’ as significant pathways in mTLE-HS **(Supplementary Fig. S15**).

IPA core analysis (adj p<0.05) of circadian and neuroinflammatory signaling pathways in human mTLE-HS revealed dysregulation of the molecular clock machinery, with mild activation of *BMAL1, CLOCK, PER1, PER2,* and *NR1D1,* while *NR2D2* was inhibited, leading to *NFIL3* inhibition and *CREB* upregulation and activating of *PER1* (**Supplementary Fig. S16)**. The neuroinflammation signaling analysis indicated microglia-mediated neuroinflammation, with *IFN-γ* activating *GSK3B,* leading to *NFKB* upregulation and activation of pro-inflammatory cytokines (e.g. *IL-6*, *IL-12*, *IL-4*). *TLR2* and *TLR4* activation resulted in *CASP8, CASP3,* and *IFN-β* activation, respectively, while *CX3CR1, IL-1β,* and *TNF-α* were downregulated (**Supplementary Fig. S17)**.

We also observed a predicted dysregulation of glutamate and GABA turnover in mTLE-HS. Genes that encode GABA-A receptor were found upregulated, while genes that encode for NMDA receptor were downregulated; potentially due to long-term anti-seizure medication use^33^ or induced by anesthetic drugs during surgery^34^. IPA core analysis of *BMAL1* as an upstream regulator in mTLE-HS showed that *BMAL1* led to upregulation of *EPAS1, FLT1, IL-1R2, NOS3, NQO1,* and *SLC2A,* and downregulation of *IL-17D* (**Supplementary Fig. S18**). Notably, *IL-1R2, IL-17D,* and *NQO1* are involved in cytokines signaling and cell defense^31,35,36^.

### Key transcriptomic targets of experimental epileptogenesis persist in mTLE-HS pathology

Next, we investigated whether the transcriptomic targets identified during experimental epileptogenesis remain relevant in human mTLE-HS. Using the Biomart tool from Ensembl, we converted human genes into their mouse orthologs and performed a cross-dataset analysis. This involved overlaying the DEGs from the RNAseq dataset of experimental epileptogenesis with those from mTLE-HS patients (coding and non-coding DEGs with adj p≤0.05). Out of the 3,660 DEGs identified in mTLE-HS, 3,229 (1,535 upregulated and 1,694 downregulated) were converted to mouse orthologs.

Our analyses revealed the overlap of 668 DEGs in human mTLE-HS and those in experimental epileptogenesis. Specifically, we identified 364 commonly upregulated genes and 304 commonly downregulated genes between the two datasets (see **Fig. 7c-h** and **Supplementary Data 2e** for the list of genes and specific overlaps per ZT). Interestingly, some genes exhibited a reversed expression pattern; i.e. several genes upregulated in murine epileptogenesis were downregulated in mTLE-HS, and vice versa. This inverted expression involved 546 genes at ZT0, 601 at ZT4, 553 at ZT8, 285 at ZT12, 291 at ZT16, and 331 at ZT20 (**Fig. 7c-h**). Out of 12 core clock genes analyzed, only *PER3* (upregulated) and *RORC* (downregulated) overlapped. We also identified that 17 genes that acquired and 301 genes that lost rhythmicity in experimental epileptogenesis overlapped with the identified orthologous DEGs in mTLE-HS (**Supplementary Fig. S19** and **Supplementary Data 2f**).

Using the Reactome database, we further analyzed the 668 DEGs common to both murine and human datasets and identified that the top enriched pathways were ‘immune system’, ‘cytokine signaling’, and ‘signaling by Interleukin’ (**Supplementary Fig. S20**). Progressing with the IPA comparative analyses, we found significant similarities in circadian and neuroinflammatory signaling components. While the molecular clock machinery was inhibited in experimental epileptogenesis, it showed mild activation in mTLE-HS (**Supplementary Fig. S16**). This could be due to differences in pathophysiological conditions, the analysis of a single time point for mTLE-HS, or anesthesia-induced circadian changes^37^. In terms of neuroinflammation, both mTLE-HS and experimental epileptogenesis showed upregulation of *NFKB, TLR2*, *TLR4, IL-6*, *IL-12* and *IL-4*, downregulation of *CX3CR1,* and dysregulation of glutamate-GABA turnover (**Supplementary Fig. S17**).

Finally, we explored the overlap between the predicted *Bmal1* targets in experimental epileptogenesis and the mTLE-HS orthologous dataset (**Supplementary Fig. S21**). We identified 10 orthologous genes downregulated in mTLE-HS, which are involved in metabolism (*ACOX1*, *HK2*, *NPY*, *SCD2, SCD4*), inflammation (*CCL2*, *LAMP2,* and *ANGPT1*), and circadian regulation (*DBP* and *PER3*; **Supplementary Fig. S18** and **Supplementary Table 7**). Additionally, three orthologous genes, associated with oxidative stress response (*HMOX1*, *HSPA5, and NQO1*; **Supplementary Fig. S18** and **Supplementary Table 7**), were upregulated in mTLE-HS.

Together, our findings indicate that key transcriptomic targets identified in experimental epileptogenesis are relevant to mTLE-HS pathology, providing insights into potential therapeutic targets for this condition.

## DISCUSSION

We showed that epileptogenesis triggers large changes in the hippocampal molecular rhythmicity, while the molecular clock machinery is inhibited. These differences are exacerbated in the light phase. Here, we also pinpointed microglia-mediated neuroinflammation and metabolism as key dysregulated pathways in epileptogenesis, processes highly regulated by *Bmal1*. Finally, we explored relevant pathways and targets shared between experimental epileptogenesis and human mTLE-HS, inferring the translational relevance of the identified targets. Our findings revealed novel aspects to the epileptogenesis pathophysiology and suggest potential novel therapeutic targets.

In mammals, different organs and tissues present a unique transcriptome profile and molecular rhythmicity that is largely controlled by the molecular clock^7,38^. The hippocampal circadian oscillatory profile of healthy mice was described by Debski et al. (2020) who identified 1,256 daily oscillating transcripts, a profile that was modified when the experimental epilepsy was induced by pilocarpine^8^. Accordingly, our RNAseq identified 1,489 genes oscillating in the hippocampus of control mice. Our study, on the other hand, comprehensively mapped transcriptomic changes during epileptogenesis, a moment where there is no presence of seizures. We found that 1,468 genes lost their diurnal rhythmicity in i.a.KA-induced epileptogenesis, while 140 acquired new rhythms, and only 21 genes kept oscillating, most of which with an inverted phase. Clearly, these findings highlight a large dysfunction of the diurnal transcriptomic rhythmicity of the hippocampus during epileptogenesis.

BMAL1 is thought as the indispensable and primary regulator of the TTFL-driven circadian rhythmicity in mammals^39,40^. Depletion of *Bmal1* abolishes the circadian rhythms^20^, disarrays the sleep-wake cycle^41^, and disrupts the rhythmicity of other core clock components^42^. Exploring the rhythmicity of the core clock genes, we found that *Bmal1, Nr1d1*, and *Per1* lost their diurnal rhythms in epileptogenesis. The IPA analysis revealed that the core clock machinery was inactive in the mouse hippocampus. Indeed, *Bmal1* malfunction was detected as early as 1h after i.a.KA-induced status epilepticus. Thus, we propose that Bmal1 dysregulation may be key to trigger a widespread dysregulation in signaling pathways essential for brain homeostasis, intensifying the epileptogenic process. This still needs to be further explored.

The TTFL clock functioning is also fundamental to the mammalian organism to adapt its physiology in anticipation of transitions between night and day^38^. In people with epilepsy, nocturnal seizures most often occur in the early morning around 5 to 6 am^43^, which highlights the change in light exposure as periods of higher risk of having seizures. In addition, immune responses are stronger in the second half of the night and early mornings in humans^44,45^ or during the light phase in rodents^46^. Our RNAseq analysis revealed that the transcriptome differences during epileptogenesis are exacerbated in the light phase, with a total of 5,901 dysregulated genes, while a total of 2,635 were dysregulated during epileptogenesis in the dark phase. When analyzing the top disease and biological processes activated/inhibited during epileptogenesis, the comparative IPA core analysis indicated seizure and epilepsy-related processes active during the light phase, but not in the dark phase. The Reactome and top disease and biological processes analysis revealed immune system and neuroinflammation as the main processes in the hippocampus during epileptogenesis. Additional experiments are required to determine whether this phenomenon is occurring as a direct consequence of molecular clock disruption or just due to cellular cascades that follow status epilepticus evolution over time.

The crosstalk between neuroinflammation and metabolism has been described in the pathophysiology of epilepsy^47,48^. Neuroinflammation and microglial hyperactivation seem to be a key mechanism of epileptogenesis process^49,50^. Microglia activity is highly controlled by BMAL1 and the circadian system with several cytokines presenting oscillatory expression^45,51^. Here we further this knowledge by showing that key components of neuroinflammation are active in microglia during early epileptogenesis, and present a distinct rhythmic profile over time. When analyzing *Bmal1* as an upstream regulator in experimental epileptogenesis, we identified a dysregulation relationship of key metabolism and immune homeostasis targets. In fact, the function of microglia seems to largely depend on metabolic adaptations and vice versa^52^. This led us to speculate that the loss of *Bmal1*’s rhythmicity also triggers the dysregulation of essential immunometabolic components. Although experimental evidence of protein binding is required, our findings provide a rationale on the intricate pathophysiological link between microglia-mediated neuroinflammation, metabolism, and circadian regulation during epileptogenesis, hitherto poorly understood. Future work will investigate how much *Bmal1* disruption is required to trigger microglia-mediated neuroinflammatory changes and seizures.

Molecular characterization of human mTLE-HS has revealed large gene expression changes in the hippocampus, as previously reported^53^. Transcriptomic studies have shown microglial dysfunctions and neuronal loss as key characteristics of human mTLE-HS^53–56^. In our study, we found 3,660 DEGs in human mTLE-HS hippocampal samples in comparison to controls. This is aligned with the studies from Salman et al. (2017)^57^ and Mills et al. (2020)^53^ that, regardless of methodological differences, investigated similar cohorts and brain region. Accordingly, these studies respectively identified MAPK signaling, and immune function, cytokine signaling, MAPK signaling and PI3K/AKT network, as main pathways. Dixit et al. (2016) similarly identified neuroinflammation, as well as modulation of neuronal networks and synaptic transmission as key pathways^58^. Our *in silico* predictions identified microglia as the most relevant cell-type in mTLE-HS hippocampal samples. Current single-cell transcriptomic work has been focusing on exploring neocortical neurons and other glial cells^5,59^. Single-cell transcriptome analysis in epileptic perilesional brain tissue from children with drug-refractory found extensive activation of microglia and infiltration of other pro-inflammatory immune cells^56^.

Although homology between a rodent model and human epilepsy cannot be claimed, experimental models must reproduce the main characteristics of the neurological disorder of interest^60^. In this study, we took advantage of the human mTLE-HS cohort to further refine our experimental target selection for future investigations, rather than directly compare the two cohorts. Epileptogenic-related changes will not necessarily be reflected at the established mTLE stage. Moreover, one must consider that the evaluation encompasses not only two completely different scenarios (epileptogenesis vs mTLE-HS) but also different species (mouse vs human). Furthermore, a striking feature of human mTLE is its heterogeneity: there are several differences in forms of HS^61^, symptoms presentation, comorbidities, and response to pharmacological treatments^55^. Differences in time of the autopsy for the human controls may add on variability. Regardless, our findings revealed a vast overlap between the DEGs found in the RNAseq in experimental epileptogenesis with the DEGs in mTLE-HS patients.

The Reactome and KEGG analysis revealed that immune system and cytokine signaling are key pathways in both datasets, corroborating the translational relevance of our findings in experimental epileptogenesis. Despite these similarities, we acknowledge certain limitations by restricting the murine study to a single epileptogenesis model and sex.

In summary, our findings showed that epileptogenesis reshapes the molecular rhythmicity of the mouse hippocampus, which is associated with the inhibition of the molecular clock machinery. We found that microglial neuroinflammation is a key player in epileptogenesis and uniquely dynamic at transcriptional level. We also reinforced the translational relevance of our findings by identifying key processes in human mTLE-HS and cross-analyzing datasets. Finally, our study shines a light on the intricate pathophysiological mechanisms of the epileptogenesis process while uncovering potential new therapeutic targets for human mTLE-HS.

## METHODS

### Epileptogenesis model in mice and temporal sampling

Adult male C57BL/6-OlaHsd mice (∼9 weeks old, 20–25 g) mice were obtained from RCSI’s Biomedical Research Facility (original stock from Harlan, UK) and were housed (up to 5 mice per cage) in on-site barrier-controlled facilities with light/dark cycle (12h/12h; light phase starts at 8 am), with food and water *ad libitum*. All animal procedures were performed in accordance with the European Communities Council Directive (2010/63/EU), and the NIH *Guide for the Care and Use of Laboratory Animals* and followed ARRIVE guidelines. All experiments were approved by the Research Ethics Committee of the RCSI University of Medicine and Health Sciences (REC1587b) under license from the Health Products Regulatory Authority (AE19127/P057), Dublin, Ireland. Sample sizes were defined based on our previous work^62^ and were approved by a biostatistician. Power calculations to determine group sizes were performed using G*Power Software v3.1, in which the group sizes were calculated to provide at least 80% power and alpha=0.05. Experimental designs and sample sizes were not altered during or after study completion, with the exception of one hippocampal sample (i.a.PBS at ZT12) which was excluded from the analysis due to technical issues during the sample processing for RNAseq.

Two sets of mice were used in this study and are represented in **Fig. 1a** and **Supplementary Fig. S1a**, respectively. For the RNA sequencing experiments, a total of 60 mice were anesthetized with isoflurane (5% for induction, 1-2% for maintenance) and subcutaneous bupivacaine (3 mg/kg) and underwent stereotaxic surgery (Day-7) to implant a guide cannula to allow i.a. injections. After local analgesia with lidocaine (2.5%) and prilocaine (2.5%) cream (EMLA, Actavis Pharma, USA), a midline scalp incision and partial craniotomy were made to expose the skull, and place a guide cannula resting on the dura mater (coordinates from the bregma: anterior-posterior (AP), −0.95 mm; lateral (L), −2.85 mm). Surgical site was covered and protected with dental cement (Simplex Powder and Liquid Mc, UK). The body temperature was controlled and kept within the normal physiological range with a feedback-controlled heat pad / rectal thermometer (Harvard Apparatus Ltd, UK). After the procedure, the animals were kept in an observation chamber with controlled temperature until full recovery and then placed back in their respective housing cage. After seven days of recovery (Day 0), mice were intra-amygdalally (i.a.) injected (ventral [V], −2.0 mm below the guide cannula) with either kainic acid (i.a.KA, Sigma-Aldrich, USA; 0.3 µg/2µL; n=30) to induce status epilepticus or sterile phosphate-buffered saline (i.a.PBS; 2µL; n=30) as the vehicle control group. All the injections were done at the same time of day (8 am). After 40 min in an observation chamber, mice received an intraperitoneal injection of lorazepam (0.1 mL of 8 mg/kg solution) to terminate seizures and reduce morbidity and mortality. Status epilepticus was defined as at least 5 min of continuous convulsive behavior, which was closely monitored but not scored. Induction and monitoring of status epilepticus were performed by a researcher blind to their treatment.

From 24 h after the i.a. injections, mice were randomly distributed, transcardially perfused with ice-cold PBS, and had their ipsilateral hippocampi microdissected at their assigned ZT (n=5 for i.a.PBS and n=5 for i.a.KA, per ZT). Samples were collected at 6 different time points according to ZTs. The first time point was ZT0, at 8 am (when lights were turned on) and exactly 24 h after i.a. injections. Next, ZT4 samples were collected at 12 pm (28 h after i.a. injections) and the temporal sampling was followed by similar collections at ZT8 (4 pm; 32 h after i.a. injections), ZT12 (8 pm; 36 h after i.a. injections), ZT16 (12 am; 40 h after i.a. injections) and ZT20 (4 am; 44 h after i.a. injections). The ipsilateral hippocampus was immediately dissected, snap-frozen, and maintained at −80°C until the RNAseq analysis. Microdissections during the dark phase were fully performed under red light. The time of the day was strictly considered for all experimental designs and normal light/dark conditions were kept at all times (lights switching on at 8 am and off at 8 pm). This sampling period is considered ideal to represent epileptogenesis in the i.a.KA model because (i) mice no longer present ictal activity directly elicited by the chemoconsulvant administration; (ii) mice do not present spontaneous recurrent seizures during this period, which occurrence consistently starts after 72h (usually 3-5 days after status epilepticus)^27,28^.

To assess *Bmal1* levels by qPCR, 20 mice were randomly distributed and underwent identical surgical procedures, i.a. and lorazepam injections and monitoring, as described above, with the exception of time of the day (i.e., surgeries and i.a. injections were performed at 12 pm). At 1 h after i.a. injections, 8 mice (n=4 for i.a.PBS; n=4 for i.a.KA) were transcardially perfused and had their ipsilateral hippocampi microdissected, snap-frozen, and maintained at −80°C until the qPCR analysis. Next, the remaining 12 mice (n=7 for i.a.PBS; n=5 for i.a.KA) were accordingly euthanized at 24 h after the i.a. injections (at 12 pm) and samples were stored as before.

### Human samples for gene expression analysis

All patients underwent clinical evaluation at the Epilepsy Service of the University of Campinas Hospital (UNICAMP). Those with well-defined epileptic focus in the medial temporal structures and meeting the criteria for drug-resistant to anti-seizure medications as defined by the ILAE were referred for amygdalohippocampectomy surgery. Informed consent was obtained from all participants and/or guardians as required, and the project was approved by the UNICAMP Research Ethics Committee under protocol number CAAE 12112913.3.0000.5404. Patients with familial history, other neurological/psychiatric disorders, or dual-pathogenesis were excluded. Furthermore, the exclusion criteria for control samples were the occurrence of any neurological disease, any viral cause of death such as HIV and hepatitis, or any other reason that would directly/indirectly affect the brain tissue.

Seventeen samples of human hippocampal tissue biopsies were collected during amygdalohippocampectomy surgeries at 2 pm (Brasilia local time, GMT-3). The tissue was immediately frozen in isopentane (−60°C) over dry ice and transported to the Molecular Genetics Laboratory for storage at −80°C. Hippocampal tissue from six autopsied individuals was used as control. All tissues were diagnosed by a neuropathologist (F.R.) using the ILAE classification^61^. The hippocampus was examined and separately rated for the presence and severity of neuronal loss and gliosis, following ILAE consensus classification of hippocampal sclerosis.

All data related to patients and human biological samples were handled in accordance with the General Data Protection Regulation (GDPR) of the European Union (EU) 2016/679 and were processed and stored as per trust policies. All personal data from participants were anonymized at the source and identified by a code number during processing to ensure their protection. No biological material was transferred and gene expression analyses were performed at the local Institutions.

### RNA sequencing in mouse samples

#### RNA extraction

First, 800 μL of Trizol™ (Sigma-Aldrich, USA) was added to each ipsilateral hippocampi sample for the tissue lysis, followed by homogenization in the bioruptor (TissueLyser LT, Qiagen Ltd, Germany) for 20 s. Next, the samples were centrifuged at 12,000 g for 10 min at 4°C. The supernatant was collected, and after 5 min of incubation at room temperature, 200 μl of chloroform (Sigma-Aldrich, USA) was added to each sample to induce the phase separation. The samples were vigorously mixed for 15 s and incubated at room temperature for 3 min, followed by centrifugation at 20,800 g for 15 min at 4°C. For RNA precipitation, the upper phase was removed and 450 μl of isopropanol (Sigma-Aldrich, USA) was added to each sample. Samples were centrifuged at 21,300 g for 30 min at 4°C, then the supernatants were discarded, and 500 μl of ethanol 75% (VWR International, LLC, USA) was used to wash the pellet. Samples were centrifuged at 21,300 g for 10 min at 4°C and the ethanol was removed. Finally, pellets were left to dry completely and resuspended in 25 μl of nuclease/RNase-free water (Thermo Fisher Scientific, USA). The concentration of RNA in each sample was measured using a Nanodrop Spectrophotometer (Thermo Fisher Scientific, USA). Nuclease/RNase-free water (Thermo Fisher Scientific, USA) was added in each sample to have a final concentration of 25 ng/μl in all the samples.

#### Library preparation

The samples were rRNA depleted and prepared for sequencing using SMARTer Stranded Total RNA Sample Prep Kit - HI Mammalian (Takara Bio, France). In brief, this kit first removes ribosomal RNA (rRNA) using RiboGone technology specifically depletes nuclear rRNA sequences (5S, 5.8S, 18S, and 28S) and mitochondrial rRNA 12S. RiboGone oligos are hybridized to rRNA, which is cleaved using RNase H. First-strand synthesis is performed using random priming, adding an anchor for use with later qPCR steps. Template switching is utilized during the RT step, adding additional non-templated nucleotides to the 3’ end of the newly formed cDNA. qPCR is performed leveraging the non-templated nucleotides and the added anchor sequence to produce Illumina-compatible libraries. Prepared libraries were quality-controlled using the Bioanalyzer 2100 (Agilent Technologies, USA) and qPCR-based concentration measurements. Libraries were equimolarly pooled and sequenced, including 20% PhiX in-lane control as 150 bp paired-end reads on an Illumina HiSeq sequencer.

#### Quantification and detection of rhythmic gene expression

Sequencing data were pre-processed by trimming away low-quality bases with a Phred score below 20, removing adapter sequence, and removing the first three nucleotides from reads in FASTQ file one for each sample, as per kit manufacturer’s instructions, using Trim Galore. Quality control was performed using FASTQC to ensure high-quality data. Differential expression analysis was performed by mapping the filtered reads to the mouse genome (mm10) using Tophat2. The software featureCounts was used to quantify the number of reads mapping to each gene using gene annotation from the Ensembl release 19. The DESeq2 Likelihood ratio test (LRT) was used to detect temporal expression changes that are significantly different between i.a.KA and i.a.PBS animals. This time-resolved differential expression analysis is used to produce the heatmap. DESeq2 Wald tests were conducted to detect significant i.a.KA vs i.a.PBS gene expression changes at individual time points, which were used for the volcano plots. The R package metaCycle, which incorporates ARSER, JTK_CYCLE, and Lomb-Scargle to detect rhythmic signals from time-series datasets, was used on the gene expression data. The metaCycle function meta2d was run with each biological replicate analyzed as successive days in the analysis. This analysis was used to demonstrate the rhythmicity of the gene expression levels.

### Quantitative Polymerase Chain Reaction (qPCR) in mouse samples

For the cDNA synthesis, the High Capacity cDNA Kit (Applied Biosystems, Thermo Fisher Scientific, USA) was used. To prepare the mix solution 0.8 μl of dNTPs, 2 μl of 10× buffer, 1 μl of reverse transcriptase enzyme, 2 μl of random primers, and 6.5 μl of nuclease/RNase-free water (Thermo Fisher Scientific, USA) were used to each reaction well. 12 μl of the master mix and 8 μl of RNA samples were added into MicroAmp PCR tubes (Applied Biosystems, Thermo Fisher Scientific, USA), followed by microcentrifugation at a low speed for 5 s. The samples were placed overnight in the thermocycler (Veriti Thermocycler, Thermo Fisher Scientific, USA) for the cDNA synthesis (25°C for 10 min, 37°C for 120 min, 85 °C for 5 min, and 4 °C for ∞). Finally, the cDNA samples were diluted (1:2) with nuclease/RNase-free water (Thermo Fisher Scientific, USA) and microcentrifuged for 30 s. The qPCR was performed to evaluate the gene expression in the cDNA samples. Each reaction tube contained 2 μl of each cDNA sample, 5 μl Power up SYBR Green Master mix (Applied biosciences, Thermo Fisher Scientific, USA), 0.5 μl of the reverse primer (5 mM), and 0.5 μl of the forward primer (5 mM; Eurofins, Luxembourg), and 2 μl nuclease/RNase-free water (Thermo Fisher Scientific, USA) to a final volume of 10 μl. The plate was centrifuged at 200g, 4 °C, for 2 min and covered with a MicroAmp Optical Adhesive Film (Applied Biosystems, Thermo Fisher Scientific, USA). Finally, the plate was placed in the 7900HT Fast Real-Time PCR System (Applied Biosystems, Thermo Fisher Scientific, USA) qPCR machine to evaluate the gene expression. Data were analyzed and normalized compared to the expression of β-actin. All the primers used (Eurofins, Luxembourg) are described in **Supplementary Table 9**. The results are presented as a fold change of each gene of interest against unstimulated control conditions and the housekeeping gene (β-actin).

### RNA sequencing in human mTLE-HS samples

#### RNA extraction

RNA was extracted using Trizol™ (Life Technologies, USA) from four histological sections of 60 µm thickness. The quantification and quality assessment of RNA was conducted using the Bioanalyzer 2100 RNA Nano (Agilent Technologies, USA). Subsequently, cDNA libraries were constructed with the TruSeq Stranded Total RNA LT Kit (Illumina, USA) following the manufacturer’s instructions. The libraries’ quality and quantity were evaluated using the Bioanalyzer 2100 DNA High Sensitive assay, yielding satisfactory results. Libraries were sequenced on the Illumina Hiseq 4000 platform with 100 paired-end reads. Quality assessment of reads was conducted using FASTQC v0.11.8 software developed by the Babraham Institute (Babraham, UK), followed by adapter and low-quality read filtering using TrimGalore v0.6.6^63^. An index was constructed using human GRCh38 cDNA from GENCODE release 36 as a reference for aligning all FASTQ files. Subsequently, Salmon (version 1.4.0) was employed to quantify transcript abundances with the pre-built index and align FASTQ files^64,65^. Variance Stabilizing Transformation (VST) was then applied to the count data using the R package DESeq2 to normalize the data. PCA was performed on the VST-transformed data to visualize the overall variance and identify any batch effects. Differential expression analysis was performed utilizing the R package DESeq2, incorporating age as a covariate^66^. The FDR was controlled using the Benjamini-Hochberg correction, and gene expression changes with an adj p≤0.05 were considered significant.

### Overlap analysis mTLE-HS and experimental epileptogenesis

To explore the translational mechanisms underlying epileptogenesis, mirroring those of mTLE-HS in humans, we comprehensively analyzed DEGs from our epileptogenesis RNAseq dataset overlaid with DEGs from mTLE-HS. Initially, we utilized the Biomart tool from Ensembl to convert human genes into their mouse orthologs. To ensure a comprehensive analysis, we adjusted our filters to include all DEGs with adj p ≤ 0.05 in both datasets, regardless of gene product type (coding or non-coding) or log_2_FC values. Subsequently, we conducted an overlaps analysis using Venn diagrams using the InteractiVenn^67^ to identify correlations between upregulated and downregulated DEGs at each of the 6 ZTs. This allowed us to identify DEGs with similar expression patterns in both datasets, as well as those with opposing expressions. Furthermore, we explored mouse orthologous DEGs to overlap with (i) identified targets of *Bmal1* as an upstream regulator, (ii) relevant genes involved in the neuroinflammation signaling pathway, and (iii) the correlation between acquired or lost rhythmicity in the mouse hippocampus during epileptogenesis.

### In silico analyses

#### Gene ontology (GO) analysis

The GO analysis was performed for mouse data only. The analysis was carried out to identify the GO terms most relevant for genes displaying temporal expression changes in both i.a.KA and i.a.PBS animals. The DESeq2 Likelihood ratio test (LRT) was used separately for i.a.KA and i.a.PBS animals to test for significant gene expression differences between any time points examined in this study. The top 500 most significant DEGs from both conditions (all with p adj p<0.05) were submitted to The Database for Annotation, Visualization, and Integrated Discovery (DAVID)^29^ to assay for GO enrichment analysis. All genes expressed above 2 FPKM were used as background for DAVID.

#### Reactome and KEGG pathway analyses

Enrichment pathway analyses were performed using ShinyGo^68^ and Reactome databases. The analysis aimed to identify the top 20 significantly enriched pathways, employing an FDR cutoff of 0.05 for both mouse and human datasets. For the mouse model, analysis was conducted for the common DEGs across all 6 ZTs, employing the following filters: log_2_FC <-1.5 and > 1.5, and adj p <0.01. For DEGs in humans, the following filters were applied: log_2_FC <-1 and > 1, and adj p <0.05. The y-axis displays pathways ranked by FDR, with the color plot indicating -log(FDR) values. For pathway enrichment analysis using mTLE-HS DEG overlaps orthologous in mice, we applied more permissive filters, considering all mTLE-HS genes with an adjusted p-value < 0.05, regardless of log_2_FC levels and coding state.

#### Ingenuity Pathway Analysis

The RNAseq data from each of the 6 ZTs, as well as the human mTLE-HS RNA-seq data, were analyzed using QIAGEN’s Ingenuity Pathway Analysis (IPA, QIAGEN Inc., Germany, https://digitalinsights.qiagen.com/IPA). Briefly, for each 6 ZT dataset, the differentially expressed genes (DEGs) encoding proteins were imported into the IPA server, and gene identification numbers (gene IDs) were assigned based on Ensembl for *Mus musculus*. For mTLE-HS data, all genes were imported, regardless of coding protein status, with gene IDs based on Ensembl for Homo sapiens. The core analysis was conducted based on expression analysis using the logarithmic expression ratio. Genes were filtered according to a p-value adjustment threshold of ≤ 0.01 for each of the ZTs and ≤ 0.05 for mTLE-HS. The reference set utilized was the Ingenuity Knowledge Base (Genes Only), considering both direct and indirect relationships. Within these parameters, three main outputs were evaluated: (i) *Enrichment of canonical pathways*: Canonical pathways were identified based on interactions among DEGs present in our dataset from each ZT and mTLE-HS and known biological pathways in the IPA database. Only pathways with -log_10_ p-value and -log_10_ BH-adj ≥ 1.3 were considered. Canonical pathways were classified as activated/inhibited based on an absolute z-score cutoff of 2. (ii) *Upstream regulator analysis:* Potential regulatory factors that could be controlling the observed changes in our data were identified. This allowed us to identify *Bmal1* as an upstream regulator and the regulation of its key targets at each of the 6 ZT, as well as its corresponding Aryl Hydrocarbon Receptor Nuclear Translocator (*ARNT*) in mTLE-HS. Furthermore, the main molecular regulators that may be involved in processes related to epileptogenesis and mTLE-HS were predicted. (iii) *Comparison IPA Core Analysis related to Diseases and Functions:* We also conducted a comparative analysis using IPA, filtered for diseases and functions related exclusively to neurological diseases, inflammatory responses, and inflammatory diseases. Additionally, only significant activation/inhibition processes with an absolute z-score >2 and -log_10_ p-value and -log_10_ BH-adj p=1.3, respectively, were considered. The comparison analysis was only for the mouse dataset, comparing the diseases and functions of the 6 ZTs between them.

### Statistics and reproducibility

The number of significantly differentiated genes in the volcano plots and Venn diagrams were defined with the adj p≤0.01 for experimental epileptogenesis data and adj p≤0.05 for mTLE-HS data. Pearson correlation test was performed using the “stats” R software package. For the qPCR statistical analysis, GraphPad Prism 9.4.1 software for Windows 10 was used to perform the statistical tests and lay out the graphics. The normality of the data was confirmed using Kolmogorov-Smirnov and Shapiro-Wilk normality tests. Parametric data were analyzed using multiple unpaired two-tailed Student’s t-tests. Outliers were identified by ROUT test and were excluded from the analysis when appropriate. Statistically significant differences between the experimental groups were considered when p<0.05. All plots from the RNAseq dataset were generated in R software version 4.3.1 with packages ggplot2^69^, ggVennDiagram^70^, and “stats”^71^. All graphs show mean values ± SEM as well as individual values as dot plots. All bar graphs are overlaid with dot plots in which each dot represents the value for one animal’s sample to show the distribution of data and the number (n) of animals per group. The data were analyzed by a researcher blind to the treatment group and group distribution. The statistical test used and specific information for each experiment are indicated in the figure legends and/or Supplementary information file.

### Data Availability

The authors declare that all the data that support the findings of this study are available in the Gene Expression Omnibus (GEO) database (GSE270801 for mice RNAseq and GSE269633 for human RNAseq), within this article and Supplementary Information files or available from the corresponding author upon request.

## ACKNOWLEDGMENTS

The present study was supported by the CURE Epilepsy Cameron Boyce Foundation Taking Flight Award by the Citizens United for Research in Epilepsy, doing business as CURE Epilepsy, a fellowship from the Irish Research Council in partnership with Epilepsy Ireland (to R.B; EPSPD/2022/112) and Science Foundation Ireland (SFI; to C.R.R; 17/TIDA/5002). This publication has emanated from research supported in part by a research grant from SFI under Grant Number 21/RC/10294 and co-funded under the European Regional Development Fund and by FutureNeuro industry partners. I.L.C, F.C., D.C.F.B., E.M.B., and F.R. were supported by the Fundação de Amparo à Pesquisa do Estado de São Paulo (FAPESP), Brazil (Grants Number 2013/07559-3 to I.L.C and F.C.; 2017/26167-0 to D.C.F.B.; 18/03254-7 to E.M.B.; and 2019/08259-0 to F.R.). I.L.C. and C.L.Y. were supported by the Conselho Nacional de Pesquisa (CNPq), Brazil (Grants number 311923/2019-4 to I.L.C; and 315953/2021-7 to C.L.Y.). I.L.C. was supported by the Coordenação de Aperfeiçoamento de Pessoal de Nível Superior (CAPES), Brazil (Grant number 001). The funding bodies had no role in the study design, collection of samples, data analysis or interpretation, or in the writing of the manuscript. We would like to thank the staff in the Biomedical Research Facilities at RCSI and the operations team at FutureNeuro. We also thank Dr. Niamh M. Connolly and Dr. James O’Siorain for insights into the bioinformatics analysis, Dr. Jennifer Dowling for the discussions, and Dr Remsha Afzal for the technical support. We thank all physicians involved in this work and with the patient care. Foremost, we thank all the people with epilepsy who generously contributed by donating their tissue and data for this study.

## AUTHOR CONTRIBUTIONS

C.R.R. designed the study with the support of A.C. (murine experiments) and I.L.C. (mTLE-HS experiments). C.R.R., R.B., T.S performed the mice research, and sample collection and preparation. R.B., M.V., T.S., M.G.L., A.R., Y.J., A.W., and A.S.R. performed the molecular analyses for mice research. D.C.F.B., M.V., and M.G.L performed the bioinformatics analyses with the support of L.S. E.M.B., M.K.M.A., C.L.Y., F.C, F.R., I.L.C, D.C.F.B, and EMB. performed the collection, preparation, and analysis of the human mTLE-HS samples. C.R.R., D.H., I.L.C, and F.C. provided reagents and/or materials. R.B., D.C.F.B, K.K. and C.R.R combined the data analyses and wrote the manuscript. All authors have edited and accepted the final version of the manuscript.

