## Supplementary figures and images for "EPILEPTOGENESIS INHIBITS THE CIRCADIAN CLOCK AND RESHAPES THE DIURNAL TRANSCRIPTOMIC RHYTHMICITY IN THE MOUSE HIPPOCAMPUS"

### Supplementary Data 1

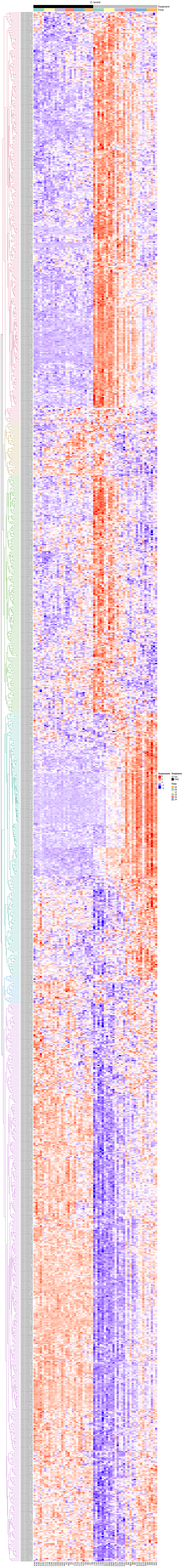
