## Supplementary Figures for "EPILEPTOGENESIS INHIBITS THE CIRCADIAN CLOCK AND RESHAPES THE DIURNAL TRANSCRIPTOMIC RHYTHMICITY IN THE MOUSE HIPPOCAMPUS"

*Running title: Epileptogenesis reshapes the molecular rhythmicity in the mouse hippocampus*

Radharani Benvenutti[1,2], Danielle C. F. Bruno[3,4,5], Matheus Gallas-Lopes[6], Morten T. Venø[7], Estela Maria Bruxel[4,5], Tammy Strickland[1,2], Arielle Ramsook[1], Aditi Wadgaonkar[1], Yiyue Jiang[1,2], Amaya Sanz-Rodriguez[2,8], Lasse Sinkkonen[3], Marina K.M. Alvim[5,9], Clarissa L. Yasuda[5,9], Fabio Rogerio[5,10], Fernando Cendes[5,9], David C. Henshall[2,8], Annie M. Curtis[1,11], Katja Kobow[12], Iscia Lopes-Cendes[4,5], Cristina R. Reschke[1,2*]

[1] School of Pharmacy and Biomolecular Sciences, RCSI University of Medicine and Health Sciences, D02 YN77, Dublin, Ireland
[2] FutureNeuro SFI Research Centre, RCSI University of Medicine and Health Sciences, D02 YN77, Dublin, Ireland
[3] Department of Life Sciences and Medicine, Faculty of Science, Technology and Medicine, University of Luxembourg, 4367, Esch-Belval Esch-sur-Alzette, Luxembourg
[4] Department of Translational Medicine, School of Medical Sciences, University of Campinas (UNICAMP), 13083-888, Campinas, Brazil
[5] Brazilian Institute of Neuroscience and Neurotechnology (BRAINN), Brazil.
[6] Pharmacology Department, Federal University of Rio Grande do Sul, 90035-003, Porto Alegre, Brazil
[7] Omiics ApS, Aarhus, Denmark
[8] Department of Physiology and Medical Physics, RCSI University of Medicine and Health Sciences, D02 YN77, Dublin, Ireland
[9] Department of Neurology, School of Medical Sciences, University of Campinas (UNICAMP), 13083-888, Campinas, Brazil
[10] Department of Pathology, School of Medical Sciences, University of Campinas (UNICAMP), 13083-888, Campinas, Brazil
[11] Tissue Engineering Research Group (TERG), RCSI University of Medicine and Health Sciences, D02 YN77, Dublin, Ireland
[12] Department of Neuropathology, Universitätsklinikum Erlangen, Friedrich-Alexander University Erlangen-Nuremberg (FAU), Erlangen, Germany

*** Corresponding author:**
Cristina Ruedell Reschke, PhD
Chrono-Epilepsy Laboratory, School of Pharmacy and Biomolecular Sciences,
RCSI University of Medicine and Health Sciences, D02 YN77, Dublin, Ireland.


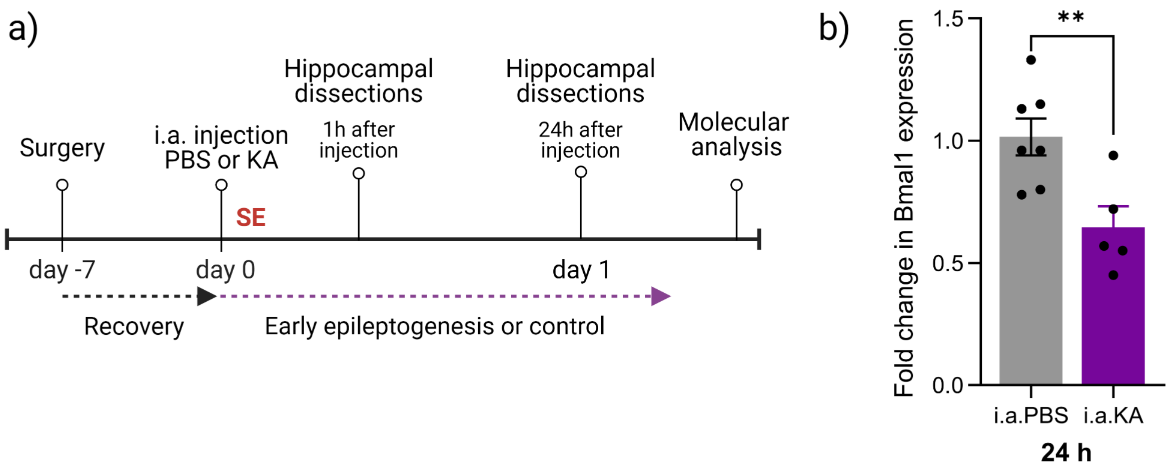


**Supplementary Figure S1. Epileptogenesis disrupts the key component of molecular clock maintenance. (a)** Schematic experimental design (created with BioRender). A surgery for cannula implantation was performed seven days before the intra-amygdala (i.a.) injection. The i.a. injection of kainic acid (KA) was used to trigger status epilepticus. Control mice were i.a. injected with phosphate-buffered saline (PBS) and subjected to the same conditions. Ipsilateral hippocampi were collected for molecular analysis 24 h after i.a. injections. **(b)** Relative gene expression (2−ΔΔCT) of *Bmal1* by qPCR. n=7 mice in i.a.PBS group and n=5 in i.a.KA group. The hippocampal dissections were performed at 12 pm. Two-tailed Student's t-test ± SEM. **p<0.01. i.a.PBS, intra-amygdala PBS injection; i.a.KA, intra-amygdala kainic acid injection; SE, status epilepticus.

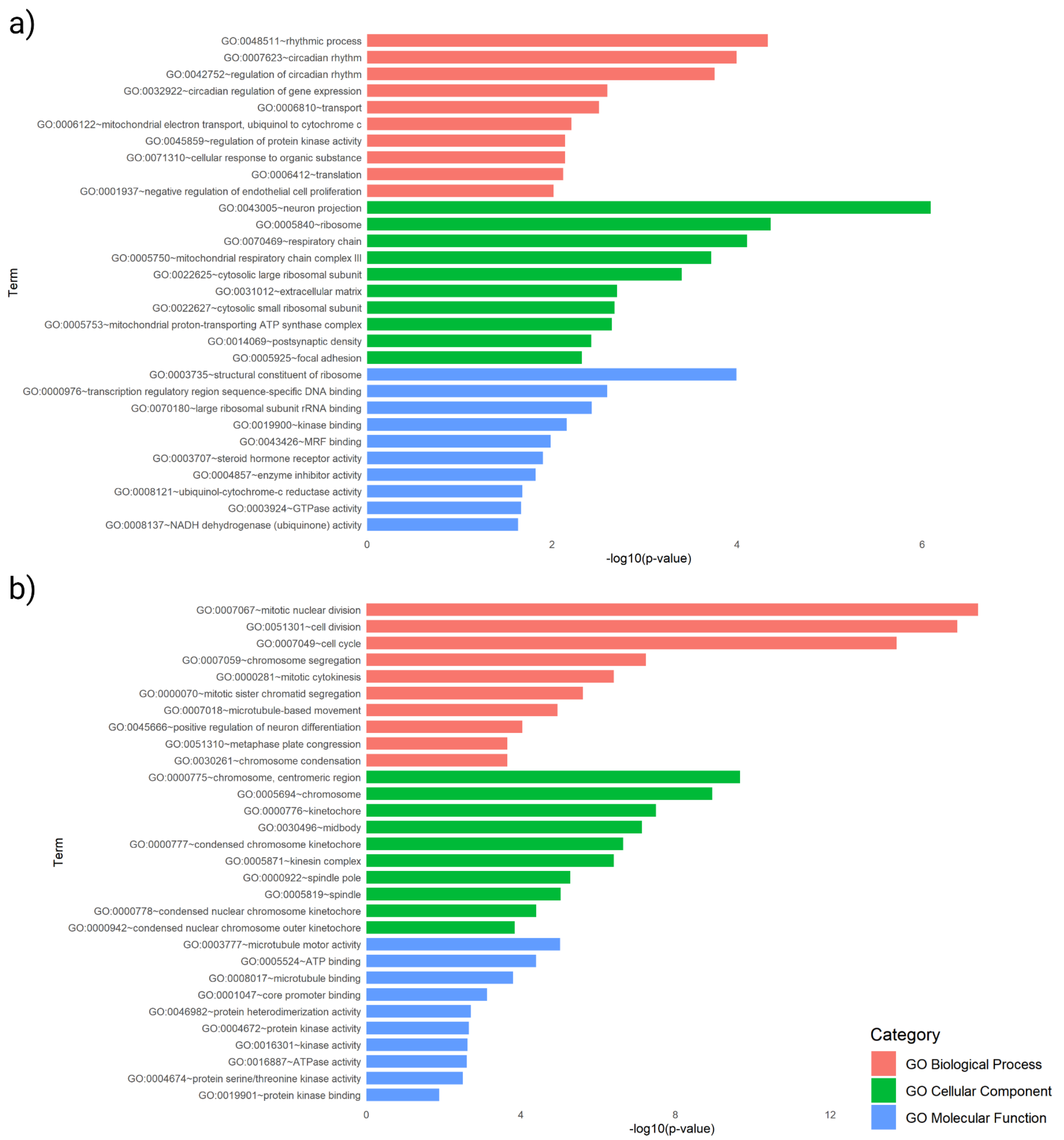


**Supplementary Figure S2. GO terms analysis from the RNAseq dataset.** The 500 top DEGs from the RNAseq dataset for **(a)** i.a.PBS and **(b)** i.a.KA were used for this analysis. The GO terms were divided into three categories: biological process, cellular component, and molecular function, represented by the colors red, green, and blue, respectively.

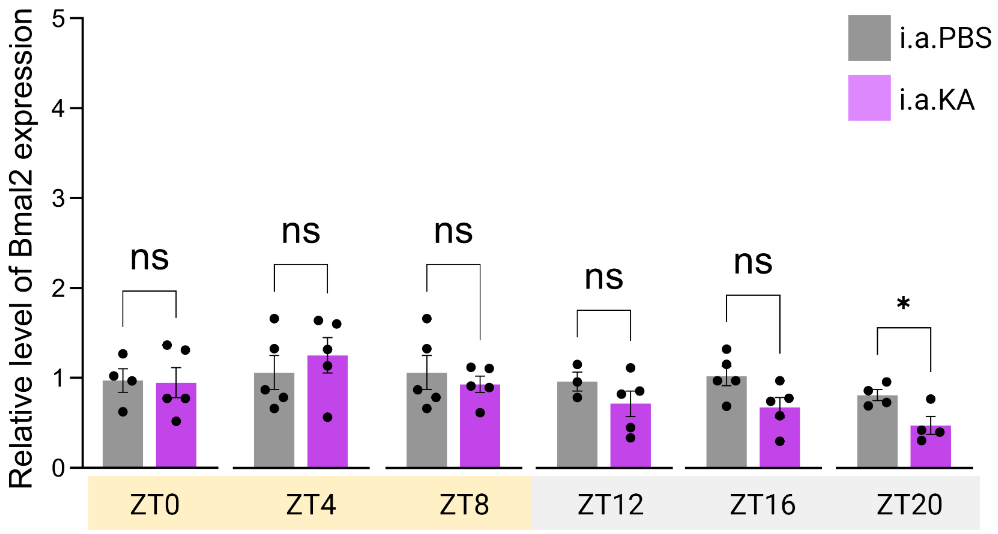


**Supplementary Figure S3.** ***Bmal2*** **is downregulated at ZT20.** Relative gene expression analysis of *Bmal2* by qPCR in all the 6 ZTs in the hippocampus during epileptogenesis. Multiple Student's t-tests ± SEM, *p <0.05. A total of 59 hippocampal samples were used, n=5 per group (except for some samples that were excluded from the analysis based on technical issues: i.a.PBS at ZT0 and ZT20, where n=4, and ZT12, where n=3 and i.a.KA at ZT20, where n=4).

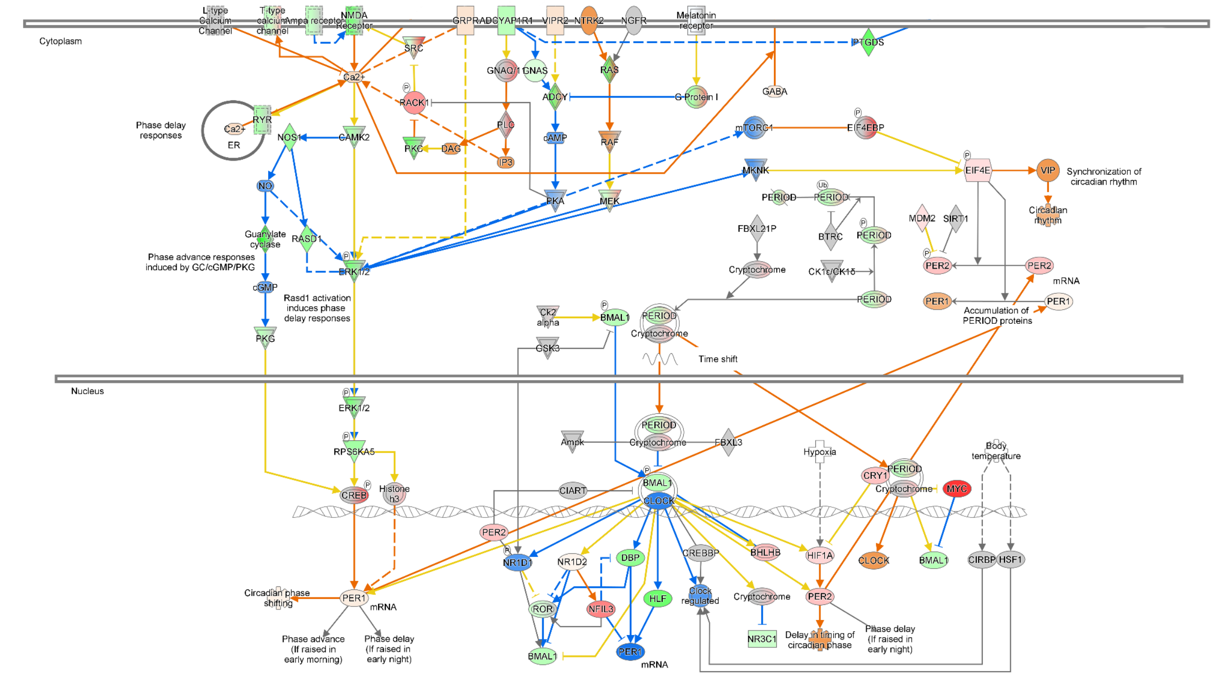


© 2000-2023 QIAGEN. All rights reserved.

**Supplementary Figure S4. The IPA signaling pathway analysis depicts the circadian molecular machinery in the hippocampus during epileptogenesis at ZT4 (12 pm).** Predictions of activation/inhibition of the pathway were identified by IPA analysis of the RNAseq dataset. The intensity of colors (green and red) corresponds to the level of $log_2FC$; weaker color indicates a lower expression level, while stronger color indicates a higher expression. Similarly, activation/inhibition of molecule activity, as defined by the activation z-score (orange) and inhibition z-score (blue), is depicted by color intensity: weaker color represents lower activation/inhibition level, whereas stronger color indicates higher activation/inhibition. Targets and/or molecular interactions marked in yellow lines represent inconsistency according to the literature; white represents molecules involved in the signaling pathway but absent in our dataset; gray represents that the genes and/or interactions are present in the dataset but the database was unable to predict their expression and/or their contribution to the pathway activation/inhibition. This figure was simplified from the IPA to focus only on the intracellular processes.

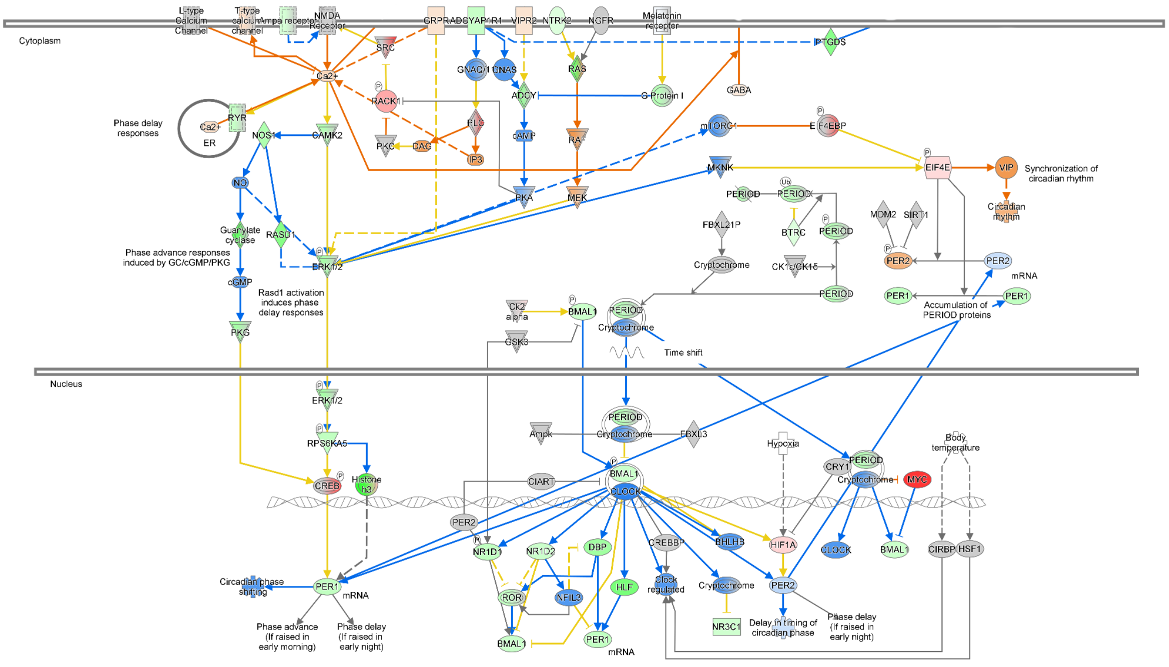


© 2000-2023 QIAGEN. All rights reserved.

**Supplementary Figure S5. The IPA signaling pathway analysis depicts the circadian molecular machinery in the hippocampus during epileptogenesis at ZT8 (4 pm).**

Predictions of activation/inhibition of the pathway were identified by IPA analysis of the RNAseq dataset. The intensity of colors (green and red) corresponds to the level of $log_2FC$; weaker color indicates a lower expression level, while stronger color indicates a higher expression. Similarly, activation/inhibition of molecule activity, as defined by the activation z-score (orange) and inhibition z-score (blue), is depicted by color intensity: weaker color represents lower activation/inhibition level, whereas stronger color indicates higher activation/inhibition. Targets and/or molecular interactions marked in yellow lines represent inconsistency according to the literature; white represents molecules involved in the signaling pathway but absent in our dataset; gray represents that the genes and/or interactions are present in the dataset but the database was unable to predict their expression and/or their contribution to the pathway activation/inhibition. This figure was simplified from the IPA to focus only on the intracellular processes.

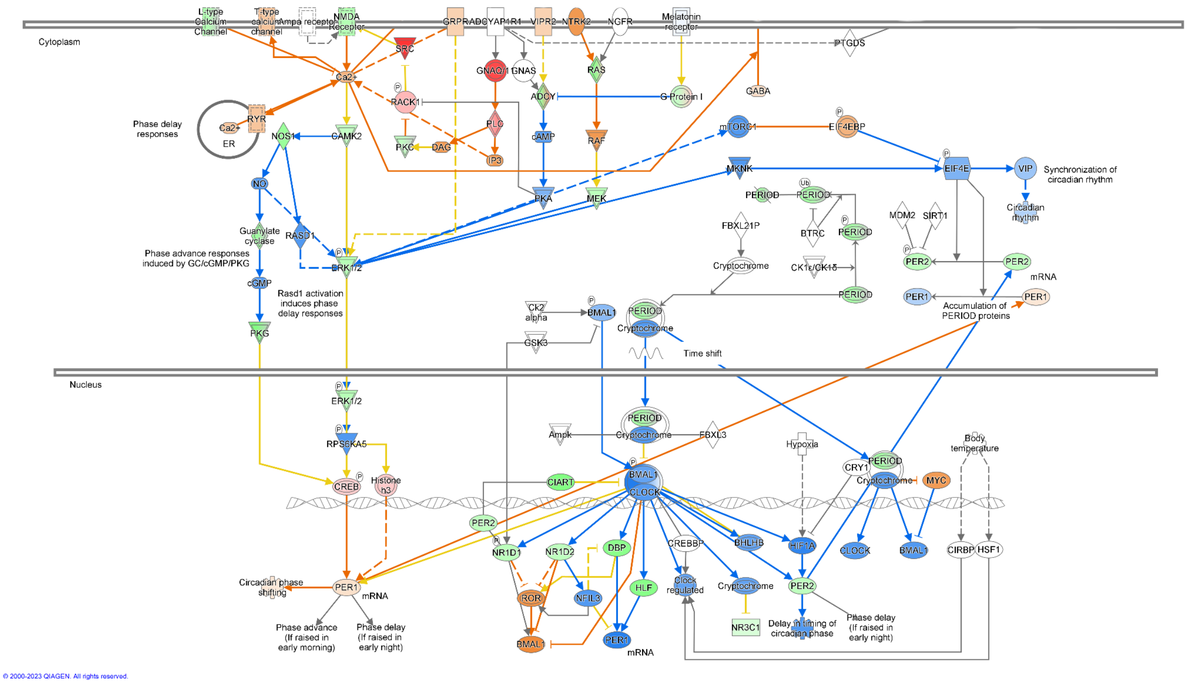


**Supplementary Figure S6. The IPA signaling pathway analysis depicts the circadian molecular machinery in the hippocampus during epileptogenesis at ZT12 (8 pm).** Predictions of activation/inhibition of the pathway were identified by IPA analysis of the RNAseq dataset. The intensity of colors (green and red) corresponds to the level of $log_2FC$;

weaker color indicates a lower expression level, while stronger color indicates a higher expression. Similarly, activation/inhibition of molecule activity, as defined by the activation z-score (orange) and inhibition z-score (blue), is depicted by color intensity: weaker color represents lower activation/inhibition level, whereas stronger color indicates higher activation/inhibition. Targets and/or molecular interactions marked in yellow lines represent inconsistency according to the literature; white represents molecules involved in the signaling pathway but absent in our dataset; gray represents that the genes and/or interactions are present in the dataset but the database was unable to predict their expression and/or their contribution to the pathway activation/inhibition. This figure was simplified from the IPA to focus only on the intracellular processes.

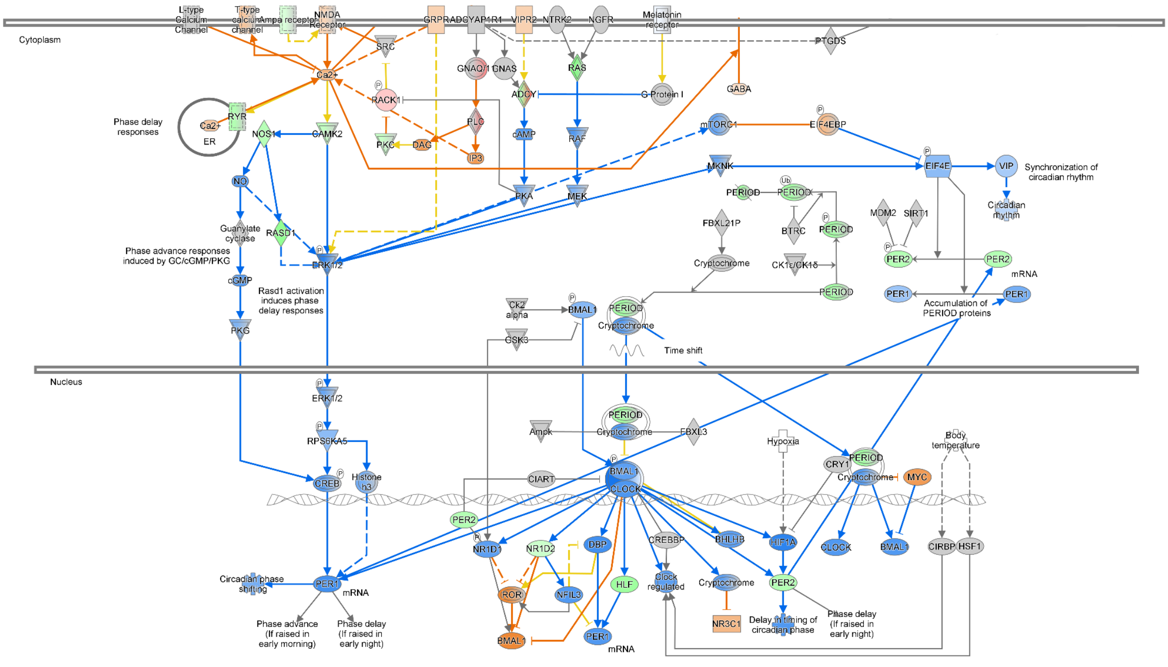

© 2000-2023 QIAGEN. All rights reserved.

**Supplementary Figure S7. The IPA signaling pathway analysis depicts the circadian molecular machinery in the hippocampus during epileptogenesis at *ZT*16 (12 am).** Predictions of activation/inhibition of the pathway were identified by IPA analysis of the RNAseq dataset. The intensity of colors (green and red) corresponds to the level of $log_2$FC; weaker color indicates a lower expression level, while stronger color indicates a higher expression. Similarly, activation/inhibition of molecule activity, as defined by the activation z-score (orange) and inhibition z-score (blue), is depicted by color intensity: weaker color represents lower activation/inhibition level, whereas stronger color indicates higher activation/inhibition. Targets and/or molecular interactions marked in yellow lines represent inconsistency according to the literature; white represents molecules involved in the signaling pathway but absent in our dataset; gray represents that the genes and/or interactions are present in the dataset but the database was unable to predict their expression and/or their contribution to the pathway activation/inhibition. This figure was simplified from the IPA to focus only on the intracellular processes.

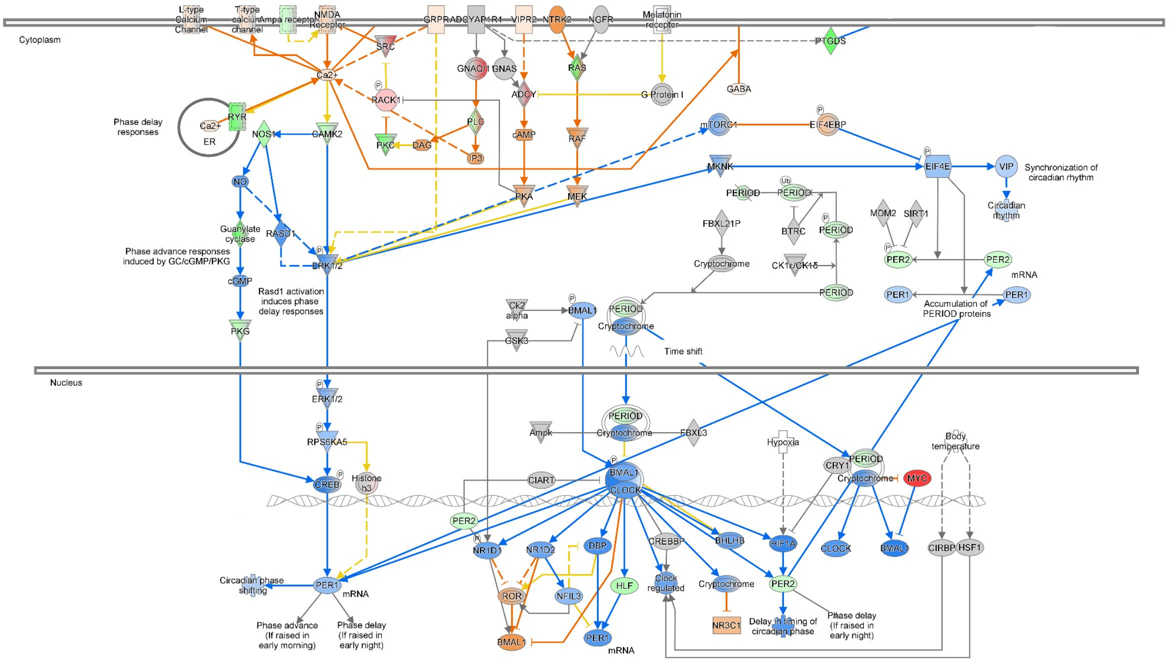


© 2000-2023 QIAGEN. All rights reserved.

**Supplementary Figure S8. The IPA signaling pathway analysis depicts the circadian molecular machinery in the hippocampus during epileptogenesis at ZT20 (4 am).** Predictions of activation/inhibition of the pathway were identified by IPA analysis of the RNAseq dataset. The intensity of colors (green and red) corresponds to the level of $log_2FC$; weaker color indicates a lower expression level, while stronger color indicates a higher expression. Similarly, activation/inhibition of molecule activity, as defined by the activation z-score (orange) and inhibition z-score (blue), is depicted by color intensity: weaker color represents lower activation/inhibition level, whereas stronger color indicates higher activation/inhibition. Targets and/or molecular interactions marked in yellow lines represent inconsistency according to the literature; white represents molecules involved in the signaling pathway but absent in our dataset; gray represents that the genes and/or interactions are present in the dataset but the database was unable to predict their expression and/or their contribution to the pathway activation/inhibition. This figure was simplified from the IPA to focus only on the intracellular processes**.**

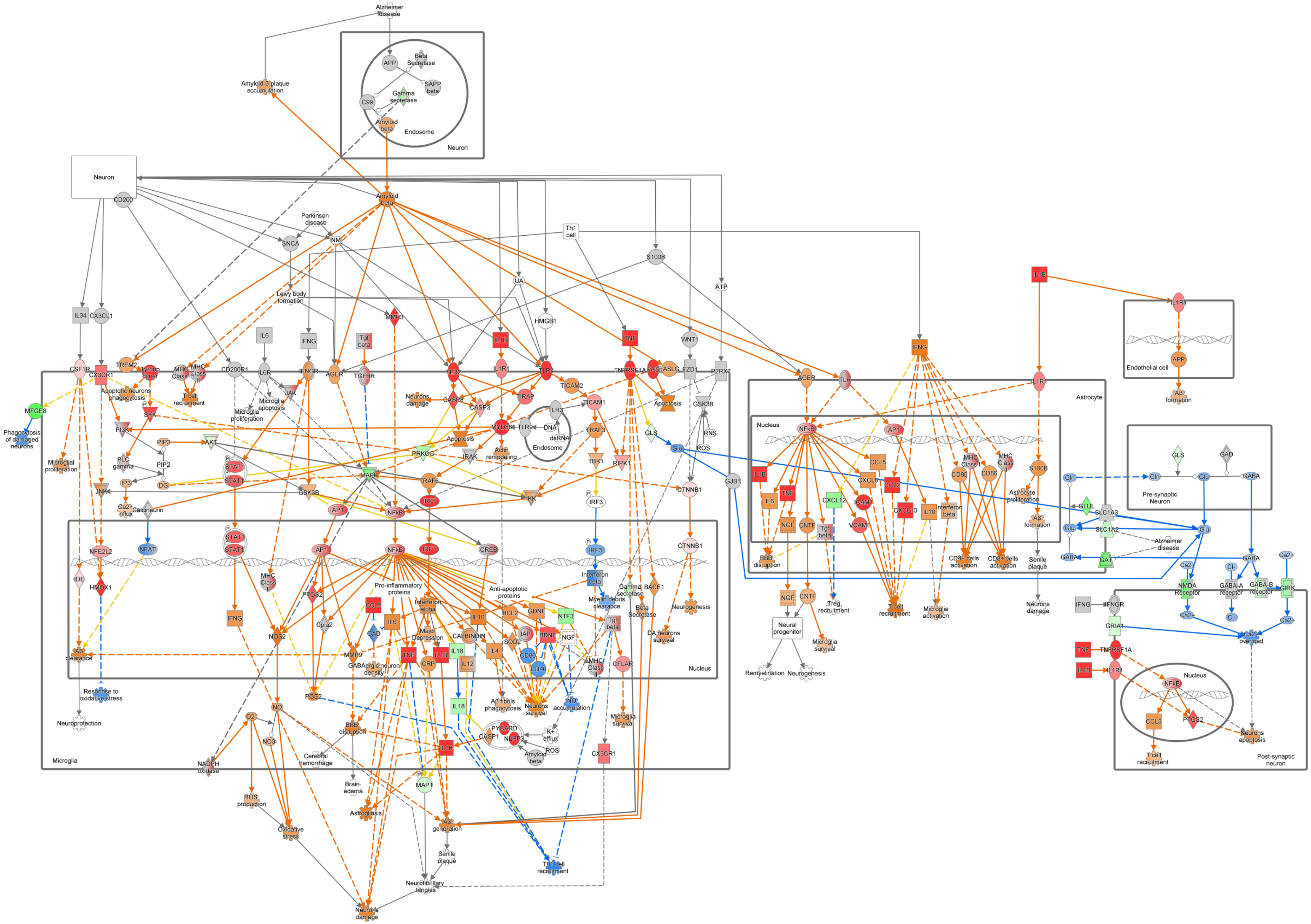
Alzheimer disease
Amyloid-β plaque accumulation
Neuron
Endosome
APP
Beta Secretase
SAPP beta
Gamma secretase
C99
Amyloid beta
CD200
Parkinson disease
SNCA
NM
Lewy body formation
Th1 cell
S100B
UA
ATP
HMGB1
MMP3
IL34
CX3CL1
IL6
IFNG
Tgf beta
IL1B
TNF
WNT1
CSF1R
CX3CR1
TREM2
CD200R1
IL6R
IFNGR
AGER
TGFBR
TLR2
IL1R1
TLR4
TNFRSF1A
FAS
FASLG
FZD1
P2RX7
MFGE8
Phagocytosis of damaged neurons
Microglial proliferation
Apoptotic neurons phagocytosis
T cell recruitment
Microglia apoptosis
Microglia proliferation
JAK
SYK
PI3K
PIP3
AKT
PLC gamma
PIP2
IP3
DG
Ca2+ influx
Calcineurin
JNK
STAT1
GSK3B
PRKCG
MAPK
Apoptosis
IRAK
Actin remodeling
MYD88
TLR9
DNA
dsRNA
TLR3
TIRAP
TICAM2
TICAM1
TRAF6
TRAF3
TBK1
IRF3
RIPK1
IKK
IRF7
AP1
NFkB
CASP8
CASP3
Neurons damage
GLS
Glu
RNS
ROS
CTNNB1
GJB1
Microglia
Nucleus
NFE2L2
NFAT
IDE
HMOX1
Pro-inflammatory proteins
Anti-apoptotic proteins
Interferon alpha
CREB
Interferon beta
Myelin debris clearance
Gamma secretase
BACE1
Beta Secretase
Neurogenesis
DA neurons survival
PTGS2
Cpla2
NOS2
MMP9
CCL2
GAD
GABAergic neuron density
Major Depression
CALBINDIN
IL10
BCL2
GDNF
NTF3
SOD2
IAP
BDNF
NGF
CD80
CD40
IL18
IL12
IL4
CRP
CFLAR
MHC Class II
Aβ clearance
Response to oxidative stress
Neuroprotection
PGE2
NO
O2
NO3
NADPH oxidase
ROS production
Oxidative stress
BBB disruption
Cerebral hemorrhage
Brain edema
Astrogliosis
MAPT
Aβ generation
Senile plaque
Neurofibrillary tangles
Aβ fibrils phagocytosis
Neurons survival
NO accumulation
PYCARD
CASP1
NLRP3
K+ efflux
Amyloid beta
Microglia survival
Th1 cell recruitment
Astrocyte
IL1R1
Endothelial cell
Aβ formation
TLR
CCL5
CXCL8
CXCL12
ICAM1
VCAM1
CXCL10
CNTF
MHC Class I
CD86
Astrocyte proliferation
CD4+ cells activation
CD8+ cells activation
Treg recruitment
Microglia activation
Neural progenitor
Remyelination
Gln
GLUL
SLC1A3
SLC1A2
GAT
GABA
Pre-synaptic Neuron
Ca2+
Cl-
NMDA Receptor
GABA-A receptor
GABA-B receptor
GIRK
IFNGR
GRIA1
Ca2+ overload
CCL3
Post-synaptic neuron
Neurons apoptosis

**Supplementary Figure S9. The IPA canonical pathway core analysis revealed dynamic microglial-mediated neuroinflammation as a key regulator of epileptogenesis at ZT4.** Log2FC and adj $p<0.01$ of differentially expressed coding genes, as well as Z-scores generated by IPA core analysis from ZT4 (12 pm), were imported into the neuroinflammation signaling pathway. Distinct colors (downregulated in green and upregulated in red) determine overlap and common relationships between genes and components of the neuroinflammation signaling pathway. The shades of green and red refer to the DEG level. Orange predicted as active and blue as inhibited. Yellow arrows highlight the main dysregulated interactions. Molecules in gray are present in the ZT4 DEGs dataset, but could not present statistical significance to predict their activation or inhibition process in the pathway. White molecules are part of the pathway but are not present in ZT4. n=5 per group.

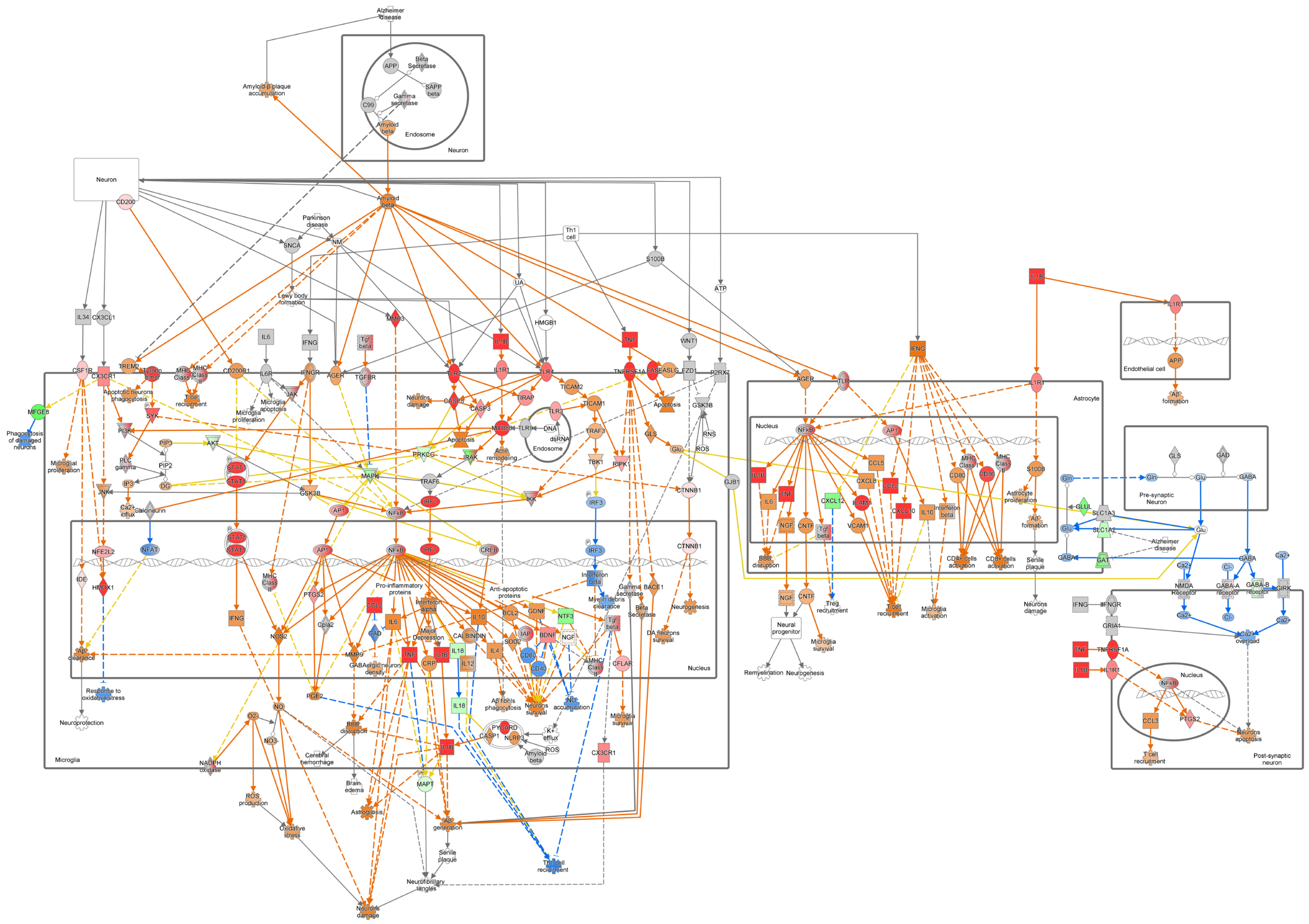
Alzheimer disease
Neuron
Endosome
Microglia
Nucleus
Astrocyte
Endothelial cell
Pre-synaptic Neuron
Post-synaptic neuron

**Supplementary Figure S10. The IPA canonical pathway core analysis revealed dynamic microglial-mediated neuroinflammation as a key regulator of epileptogenesis at ZT8.** Log2FC and adj $p<0.01$ of differentially expressed coding genes, as well as Z-scores generated by IPA core analysis from ZT8 (4 pm), were imported into the neuroinflammation signaling pathway. Distinct colors (downregulated in green and upregulated in red) determine overlap and common relationships between genes and components of the neuroinflammation signaling pathway. The shades of green and red refer to the DEG level. Orange predicted as active and blue as inhibited. Yellow arrows highlight the main dysregulated interactions. Molecules in gray are present in the ZT8 DEGs dataset, but could not present statistical significance to predict their activation or inhibition process in the pathway. White molecules are part of the pathway but are not present in ZT8. n=5 per group.

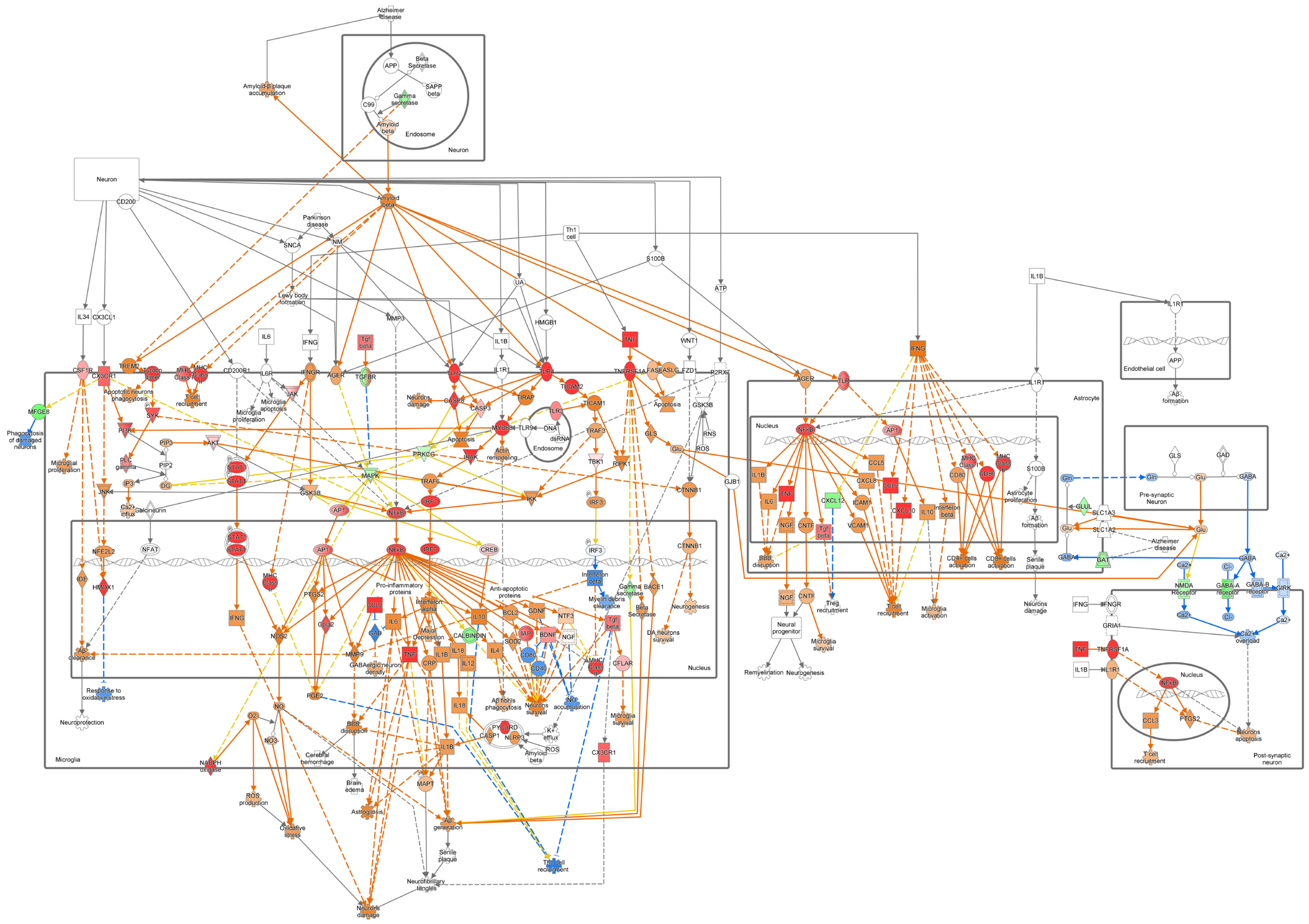

Alzheimer disease
Amyloid-β plaque accumulation
APP
Beta Secretase
SAPP beta
Gamma secretase
C99
Amyloid beta
Endosome
Neuron
CD200
Parkinson disease
SNCA
NM
Lewy body formation
Th1 cell
S100B
ATP
UA
MMP3
HMGB1
IL34
CX3CL1
IL6
IFNG
Tgf beta
IL1B
TNF
WNT1
CSF1R
CX3CR1
TREM2
Tyrobp dimer
MHC Class I
MHC Class II
CD200R1
IL6R
IFNGR
AGER
TGFBR
TLR2
IL1R1
TLR4
TICAM2
TNFRSF1A
FAS
FASLG
FZD1
P2RX7
MFGE8
Phagocytosis of damaged neurons
Apoptotic neurons phagocytosis
T cell recruitment
JAK
Microglia apoptosis
Microglia proliferation
Neurons damage
CASP8
CASP3
TIRAP
TICAM1
Apoptosis
GSK3B
RNS
SYK
PI3K
PIP3
AKT
PIP2
PLC gamma
Microglial proliferation
MYD88
TLR9
DNA
dsRNA
TLR3
TRAF3
TBK1
RIPK1
GLS
Glu
ROS
GJB1
PRKCG
IRAK
Actin remodeling
STAT1
MAPK
TRAF6
IKK
IRF7
IRF3
CTNNB1
JNK
IP3
DG
Ca2+ influx
Calcineurin
AP1
NFkB
CREB
NFE2L2
NFAT
IDE
HMOX1
Pro-inflammatory proteins
Anti-apoptotic proteins
Interferon beta
Myelin debris clearance
Gamma secretase
BACE1
Beta Secretase
Neurogenesis
DA neurons survival
PTGS2
Gfap2
CCL2
GAD
IL6
Interferon alpha
Major Depression
IL10
BCL2
GDNF
NTF3
CALBINDIN
IL4
SOD2
IAP
BDNF
NGF
CD80
CD40
MHC Class
CFLAR
NOS2
MMP9
GABAergic neuron density
TNF
CRP
IL1B
IL18
IL12
Aβ clearance
Nucleus
Response to oxidative stress
Neuroprotection
NO
O2
PGE2
NO3
BBB disruption
Cerebral hemorrhage
Brain edema
Aβ fibrils phagocytosis
Neurons survival
NO accumulation
Microglia survival
PYCARD
CASP1
NLRP3
K+ efflux
ROS
Amyloid beta
CX3CR1
Microglia
NADPH oxidase
ROS production
Oxidative stress
Astrogliosis
MAPT
Aβ generation
Senile plaque
Th2 cell recruitment
Neurofibrillary tangles
Neurons damage
AGER
TLR
IL1R
Astrocyte
NFkB
AP1
CCL5
CXCL8
CCL2
MHC Class I
MHC Class II
CD80
CD86
S100B
Astrocyte proliferation
Aβ formation
Senile plaque
CXCL12
ICAM1
VCAM1
CXCL10
IL10
Interferon beta
Tgf beta
CNTF
NGF
CD4+ cells activation
CD8+ cells activation
Treg recruitment
T cell recruitment
Microglia activation
Neurons damage
Neural progenitor
Remyelination
Neurogenesis
Microglia survival
IL1R1
Endothelial cell
Aβ formation
Gln
GLS
GAD
GABA
Pre-synaptic Neuron
GLUL
SLC1A3
SLC1A2
GAT
Alzheimer disease
Ca2+
Cl-
NMDA Receptor
GABA-A receptor
GABA-B receptor
GIRK
Ca2+ overload
IFNGR
GRIA1
TNFRSF1A
CCL3
PTGS2
Neurons apoptosis
Post-synaptic neuron

**Supplementary Figure S11. The IPA canonical pathway core analysis revealed dynamic microglial-mediated neuroinflammation as a key regulator of epileptogenesis at ZT12.** Log2FC and adj $p<0.01$ of differentially expressed coding genes, as well as Z-scores generated by IPA core analysis from ZT12 (8 pm), were imported into the neuroinflammation signaling pathway. Distinct colors (downregulated in green and upregulated in red) determine overlap and common relationships between genes and components of the neuroinflammation signaling pathway. The shades of green and red refer to the DEG level. Orange predicted as active and blue as inhibited. Yellow arrows highlight the main dysregulated interactions. Molecules in gray are present in the ZT12 DEGs dataset, but could not present statistical significance to predict their activation or inhibition process in the pathway. White molecules are part of the pathway but are not present in ZT12. n=4 for i.a.PBS at ZT12 and n=5 for i.a.KA.

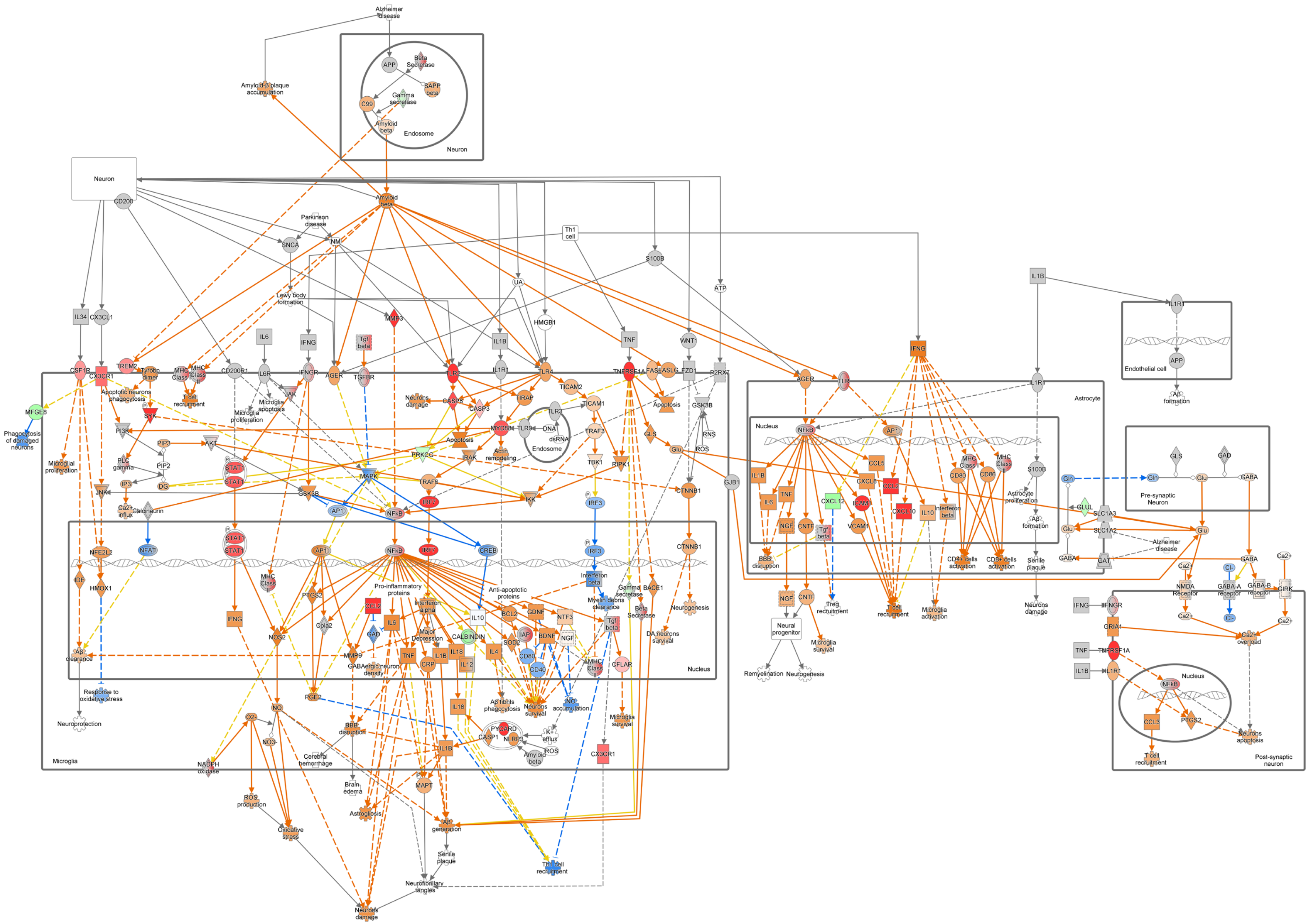
Alzheimer disease
Amyloid-β plaque accumulation
APP
Beta Secretase
SAPP beta
Gamma secretase
C99
Amyloid beta
Endosome
Neuron
CD200
Parkinson disease
SNCA
NM
Lewy body formation
Th1 cell
S100B
ATP
UA
HMGB1
IL34
CX3CL1
IL6
IFNG
Tgf beta
MMP3
IL1B
TNF
WNT1
CSF1R
TREM2
CX3CR1
Tyrobp dimer
MHC Class I
MHC Class II
CD200R1
IL6R
IFNGR
AGER
TGFBR
TLR2
IL1R1
TLR4
TNFRSF1A
FAS
FASLG
FZD1
P2RX7
MFGE8
Phagocytosis of damaged neurons
Apoptotic neurons phagocytosis
T cell recruitment
JAK
Microglia apoptosis
Microglia proliferation
Neurons damage
CASP8
CASP3
TIRAP
TICAM2
TICAM1
Apoptosis
GSK3B
SYK
PI3K
PIP3
AKT
MYD88
TLR3
DNA
dsRNA
TRAF3
GLS
Glu
RNS
ROS
Microglial proliferation
PLC gamma
PIP2
PRKCG
IRAK
Actin remodeling
TBK1
RIPK1
IP3
DG
STAT1
MAPK
TRAF6
IKK
CTNNB1
GJB1
JNK
Ca2+ influx
Calcineurin
AP1
NFkB
IRF7
CREB
IRF3
NFE2L2
NFAT
IDE
HMOX1
MHC Class II
Pro-inflammatory proteins
Anti-apoptotic proteins
Interferon beta
Gamma secretase
BACE1
Myelin debris clearance
Beta Secretase
Neurogenesis
PTGS2
CCL2
Interferon alpha
IL10
BCL2
GDNF
NTF3
Cpla2
IL6
Major Depression
CALBINDIN
IAP
SOD2
BDNF
NGF
DA neurons survival
NOS2
GAD
MMP9
TNF
IL18
CD80
CD40
CFLAR
Aβ clearance
GABAergic neuron density
CRP
IL12
Nucleus
Response to oxidative stress
PGE2
NO
Aβ fibrils phagocytosis
Neurons survival
NO accumulation
Microglia survival
Neuroprotection
O2-
NO3-
BBB disruption
PYCARD
CASP1
NLRP3
K+ efflux
Cerebral hemorrhage
Brain edema
CX3CR1
Microglia
NADPH oxidase
ROS production
Astrogliosis
Aβ generation
MAPT
Senile plaque
Oxidative stress
Th1 cell recruitment
Neurofibrillary tangles
Neurons damage
TLR
Astrocyte
CCL5
CXCL8
CXCL12
ICAM1
CXCL10
VCAM1
CNTF
MHC Class I
CD86
Interferon beta
Astrocyte proliferation
Aβ formation
CD4+ cells activation
CD8+ cells activation
Treg recruitment
Neural progenitor
Microglia survival
Remyelination
Microglia activation
Endothelial cell
IL1R1
Gln
GLUL
SLC1A3
SLC1A2
GAT
GABA
GAD
Pre-synaptic Neuron
NMDA Receptor
GABA-A receptor
GABA-B receptor
GIRK
Ca2+
Cl-
Ca2+ overload
GRIA1
CCL3
Neurons apoptosis
Post-synaptic neuron

**Supplementary Figure S12. The IPA canonical pathway core analysis revealed dynamic microglial-mediated neuroinflammation as a key regulator of epileptogenesis at ZT16.** Log2FC and adj $p<0.01$ of differentially expressed coding genes, as well as Z-scores generated by IPA core analysis from ZT16 (12 am), were imported into the neuroinflammation signaling pathway. Distinct colors (downregulated in green and upregulated in red) determine overlap and common relationships between genes and components of the neuroinflammation signaling pathway. The shades of green and red refer to the DEG level. Orange predicted as active and blue as inhibited. Yellow arrows highlight the main dysregulated interactions. Molecules in gray are present in the ZT16 DEGs dataset, but could not present statistical significance to predict their activation or inhibition process in the pathway. White molecules are part of the pathway but are not present in ZT16. n=5 per group.

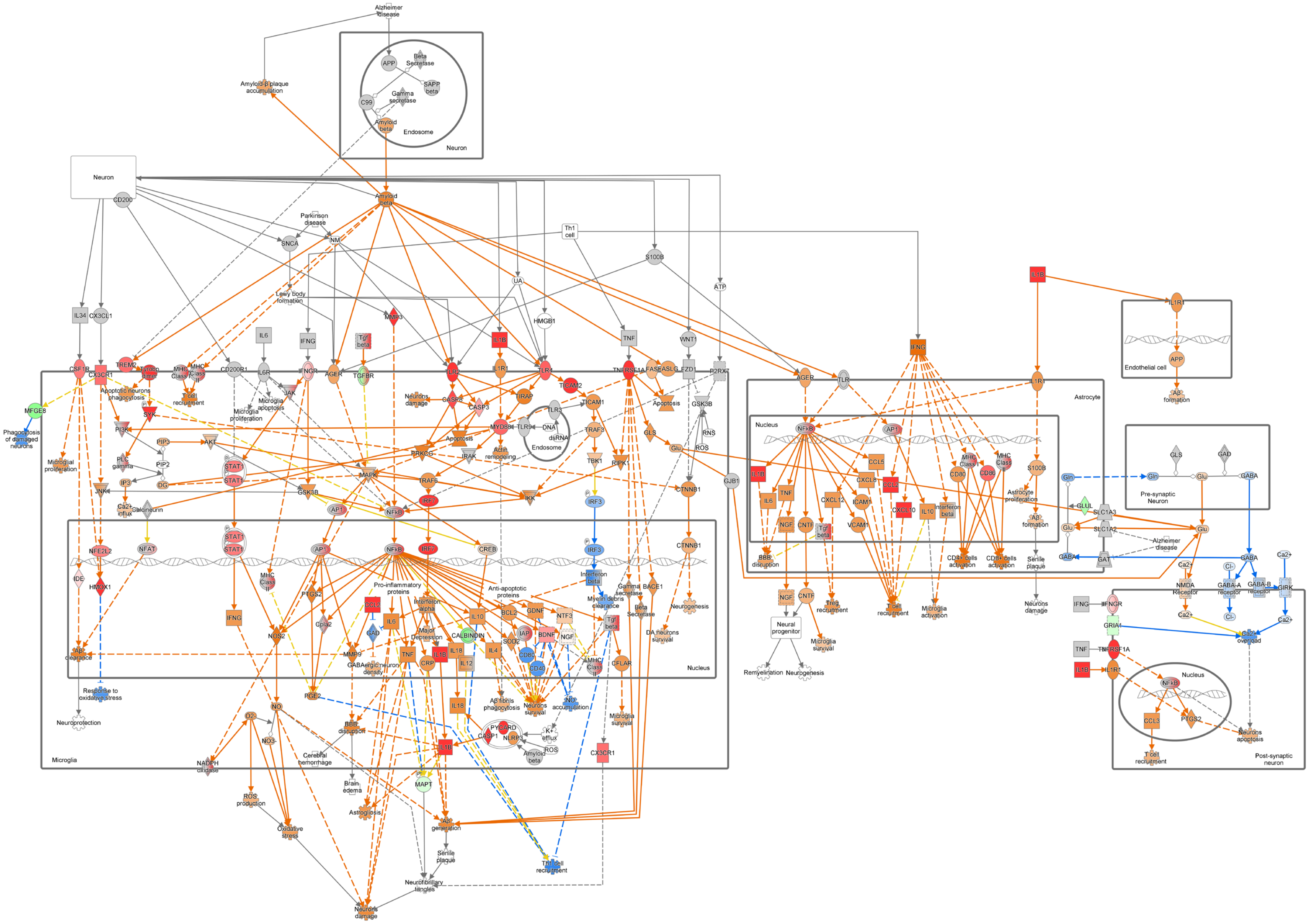

Alzheimer disease
Neuron
Endosome
Amyloid-β plaque accumulation
Amyloid beta
Parkinson disease
Lewy body formation
Microglia
Nucleus
Astrocyte
Endothelial cell
Pre-synaptic Neuron
Post-synaptic neuron
Neural progenitor
Remyelination
Neurogenesis
Neuroprotection
Neurofibrillary tangles
Neurons damage
Senile plaque
Oxidative stress
ROS production
Phagocytosis of damaged neurons
Microglial proliferation
Pro-inflammatory proteins
Anti-apoptotic proteins
Neurons survival
Aβ generation
Brain edema
Cerebral hemorrhage
Astrogliosis
Th1 cell

**Supplementary Figure S13. The IPA canonical pathway core analysis revealed dynamic microglial-mediated neuroinflammation as a key regulator of epileptogenesis at ZT20.** Log2FC and adj p<0.01 of differentially expressed coding genes, as well as Z-scores generated by IPA core analysis from ZT20 (4 am), were imported into the neuroinflammation signaling pathway. Distinct colors (downregulated in green and upregulated in red) determine overlap and common relationships between genes and components of the neuroinflammation signaling pathway. The shades of green and red refer to the DEG level. Orange predicted as active and blue as inhibited. Yellow arrows highlight the main dysregulated interactions. Molecules in gray are present in the ZT20 DEGs dataset, but could not present statistical significance to predict their activation or inhibition process in the pathway. White molecules are part of the pathway but are not present in ZT20. n=5 per group.

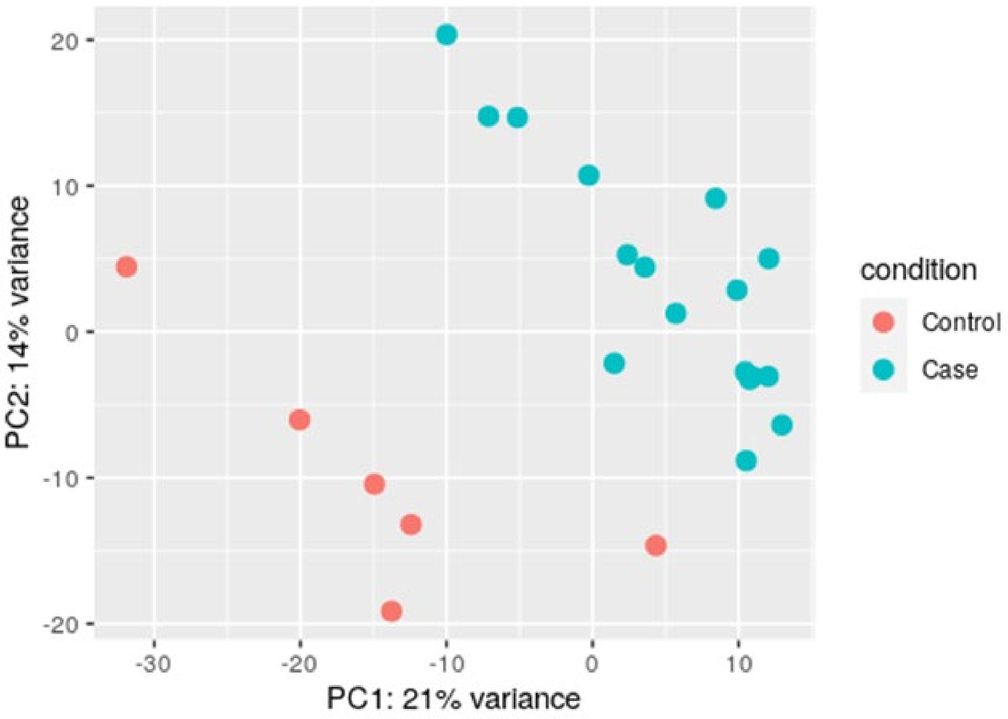


**Supplementary Figure S14. Principal component analysis (PCA) distinctly separated the control group from mTLE-HS patients.** The analysis indicates distinct hippocampal DEG profiles associated with the disease phenotype. The orange circles represent the autopsy samples from the control group (n=6), while the blue circles represent the samples from mTLE-HS patients (n=17).

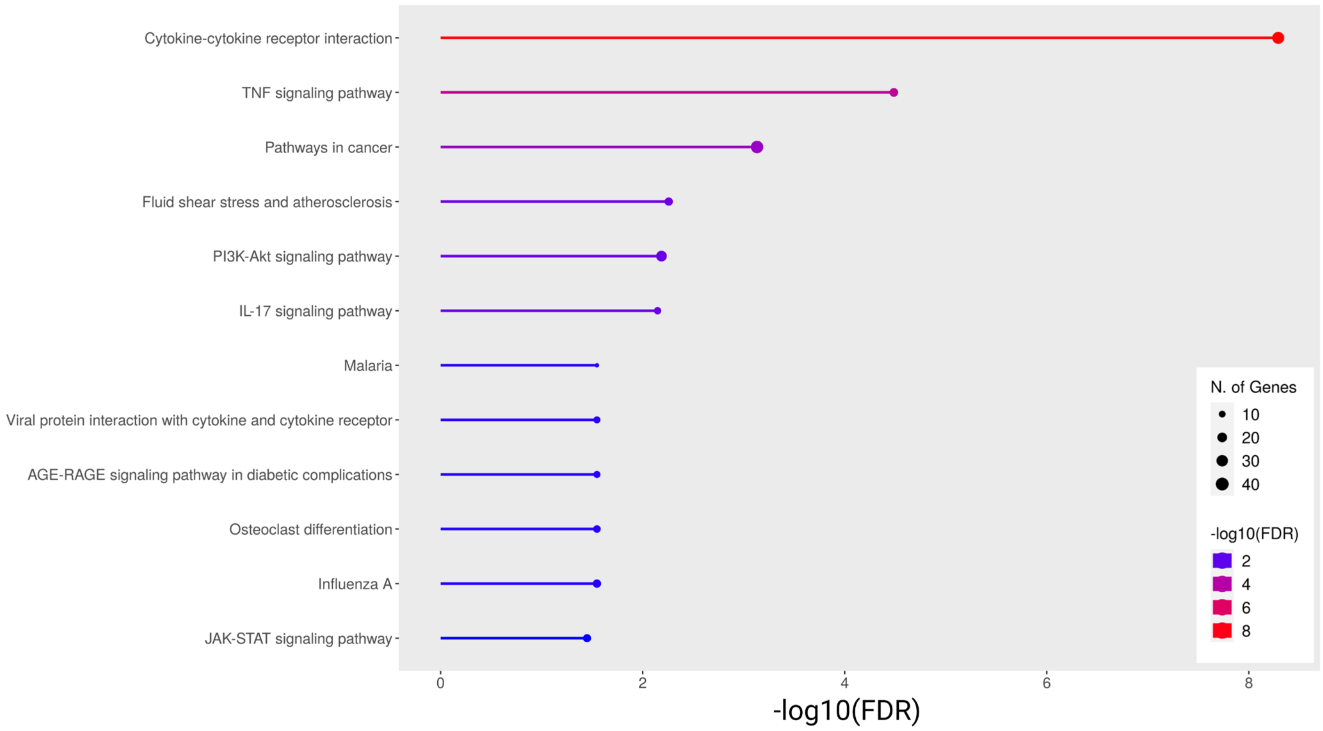


**Supplementary Figure S15. The Kyoto Encyclopedia of Genes and Genomes (KEGG) analysis revealed the most relevant pathways in human mTLE-HS.** This analysis was performed using the RNAseq dataset from human mTLE-HS (n=17) in comparison with the control group (n=6). Colors depict –log10(FDR) and circles depict the number of genes.

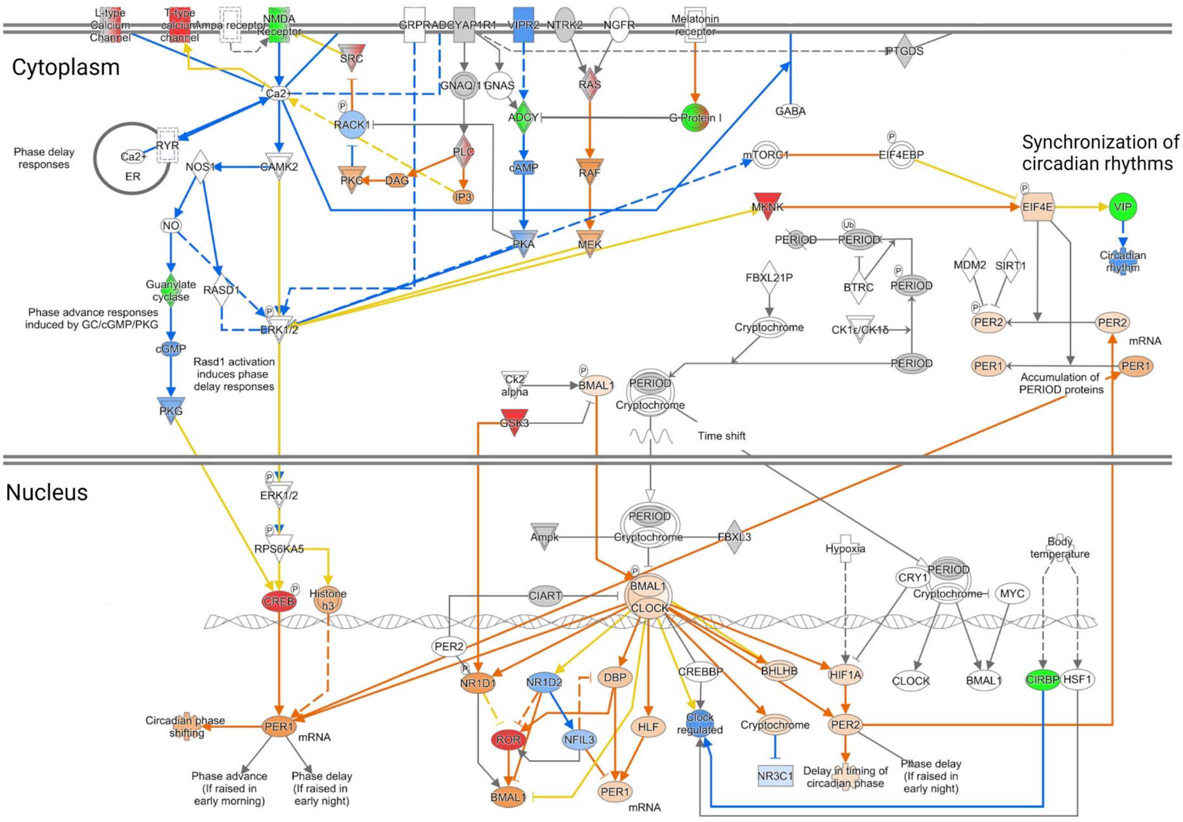


**Supplementary Figure S16. The IPA signaling pathway analysis depicts the circadian molecular machinery of hippocampal samples dissected from mTLE-HS patients.** Predictions of activation/inhibition of the pathway were identified by IPA analysis of the mTLE-HS patients' RNAseq dataset. The intensity of colors (green and red) corresponds to the level of $log_2FC$; weaker color indicates a lower expression level, while stronger color indicates a higher expression. Similarly, activation/inhibition of molecule activity, as defined by the activation z-score (orange) and inhibition z-score (blue), is depicted by color intensity: weaker color represents lower activation/inhibition level, whereas stronger color indicates higher activation/inhibition. Targets and/or molecular interactions marked in yellow lines represent inconsistency according to the literature; white represents molecules involved in the signaling pathway but absent in our dataset; gray represents that the genes and/or interactions are present in the dataset but the database was unable to predict their expression and/or their contribution to the pathway activation/inhibition.

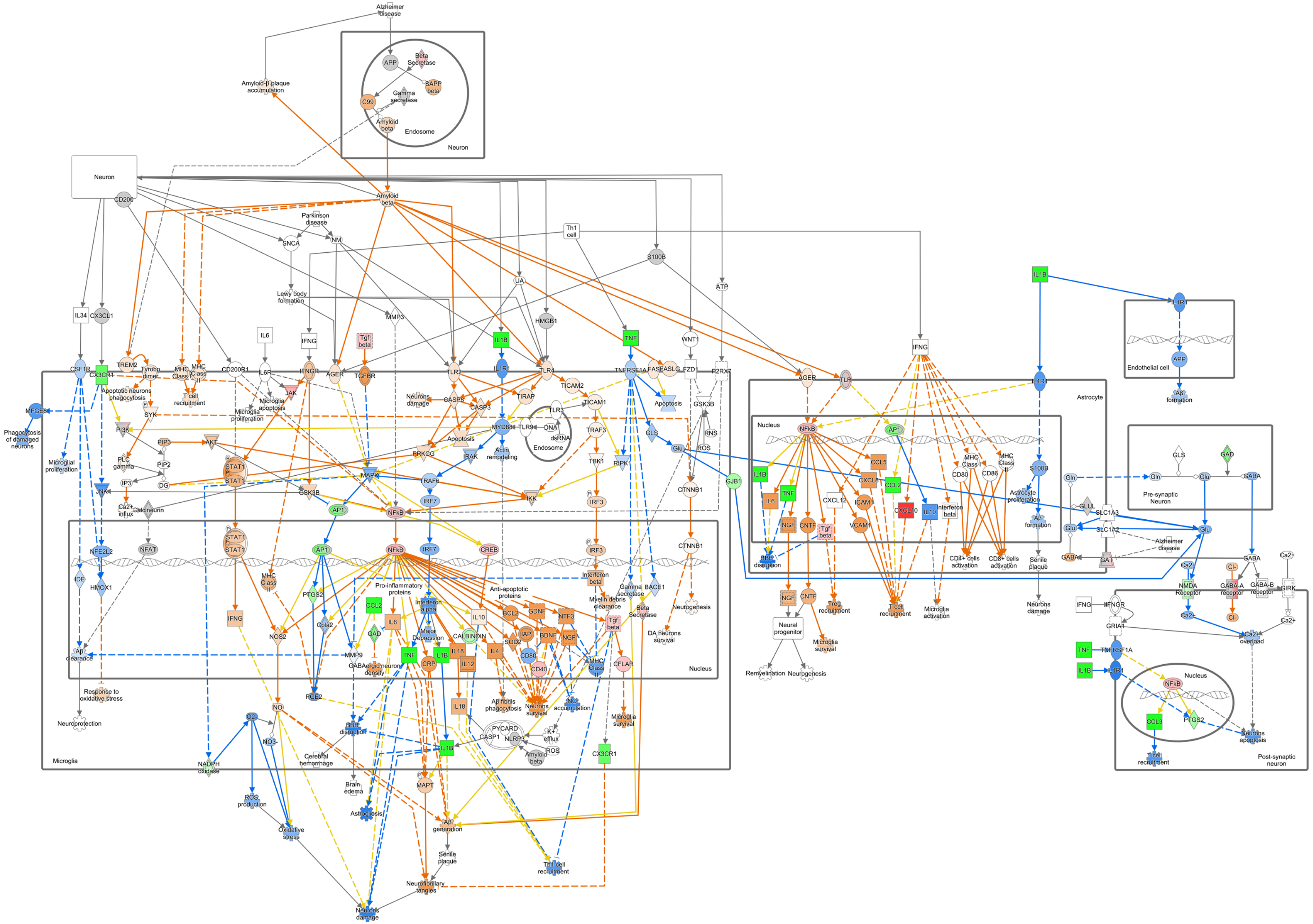
Alzheimer disease
Amyloid-β plaque accumulation
Neuron
Endosome
APP
Beta Secretase
SAPP beta
Gamma secretase
C99
Amyloid beta
CD200
Parkinson disease
SNCA
NM
Lewy body formation
Th1 cell
S100B
ATP
UA
HMGB1
MMP3
IL34
CX3CL1
IL6
IFNG
Tgf beta
WNT1
TNF
IL1B
CSF1R
CX3CR1
TREM2
Tyrobp dimer
MHC Class I
MHC Class II
T cell recruitment
CD200R1
IL6R
JAK
IFNGR
AGER
TGFBR
TLR2
IL1R1
TLR4
TICAM2
TNFRSF1A
FASLG
FAS
FZD1
P2RX7
MFGE8
Phagocytosis of damaged neurons
Apoptotic neurons phagocytosis
SYK
Microglia apoptosis
Microglial proliferation
Neurons damage
CASP8
CASP3
TIRAP
TICAM1
Apoptosis
GSK3B
PI3K
PIP3
AKT
MYD88
TLR9
TLR3
DNA
dsRNA
TRAF3
GLS
RNS
ROS
Glu
PLC gamma
PIP2
IP3
DG
STAT1
PRKCG
IRAK
Actin remodeling
TBK1
RIPK1
MAPK
TRAF6
IKK
JNK
Ca2+ influx
Calcineurin
IRF7
IRF3
CTNNB1
AP1
NFkB
CREB
NFE2L2
NFAT
IDE
HMOX1
Pro-inflammatory proteins
Anti-apoptotic proteins
Interferon beta
Gamma secretase
BACE1
Myelin debris clearance
Beta Secretase
Neurogenesis
DA neurons survival
PTGS2
CCL2
Interferon alpha
IL10
BCL2
GDNF
NTF3
NOS2
Cpla2
GAD
IL6
Major Depression
CALBINDIN
IAP
SOD2
BDNF
NGF
MHC Class I
MMP9
GABAergic neuron density
CD80
TNF
CRP
IL18
IL12
IL4
CD40
CFLAR
Aβ clearance
Response to oxidative stress
Neuroprotection
O2
NO
NO3
PGE2
BBB disruption
Aβ fibrils phagocytosis
Neurons survival
Na+ accumulation
Microglia survival
PYCARD
CASP1
NLRP3
K+ efflux
Amyloid beta
Cerebral hemorrhage
Brain edema
NADPH oxidase
ROS production
Oxidative stress
MAPT
Astrogliosis
Aβ generation
Senile plaque
Th1 cell recruitment
Neurofibrillary tangles
Neurons damage
Microglia
Nucleus
GJB1
Astrocyte
TLR
CCL5
CXCL8
ICAM1
CXCL12
CXCL10
VCAM1
CNTF
MHC Class I
MHC Class II
CD86
S100B
Astrocyte proliferation
Aβ formation
CD4+ cells activation
CD8+ cells activation
Senile plaque
Neurons damage
Treg recruitment
T cell recruitment
Microglia activation
Neural progenitor
Microglia survival
Remyelination
Neurogenesis
Endothelial cell
Gln
GLS
GAD
GABA
Pre-synaptic Neuron
GLUL
SLC1A3
SLC1A2
GAT
Ca2+
NMDA Receptor
Cl-
GABA-A receptor
GABA-B receptor
GIRK
IFNGR
GRIA1
Ca2+ overload
CCL3
Neurons apoptosis
Post-synaptic neuron

**Supplementary Figure S17. The IPA canonical pathway core analysis revealed microglial-mediated neuroinflammation as a key regulator of human mTLE-HS.** Active neuroinflammation canonical pathway using IPA core analysis. Log2FC and adj $p<0.01$ of differentially expressed coding genes, as well as Z-scores generated by IPA core analysis, were imported into the neuroinflammation signaling pathway. Distinct colors (downregulated in green and upregulated in red) determine overlap and common relationships between genes and components of the neuroinflammation signaling pathway. The shades of green and red refer to the DEG level. Orange predicted as active and blue as inhibited. Yellow arrows highlight the main dysregulated interactions. Molecules in gray are present in the dataset, but could not present statistical significance to predict their activation or inhibition process in the pathway. White molecules are part of the pathway but are not present in the dataset.

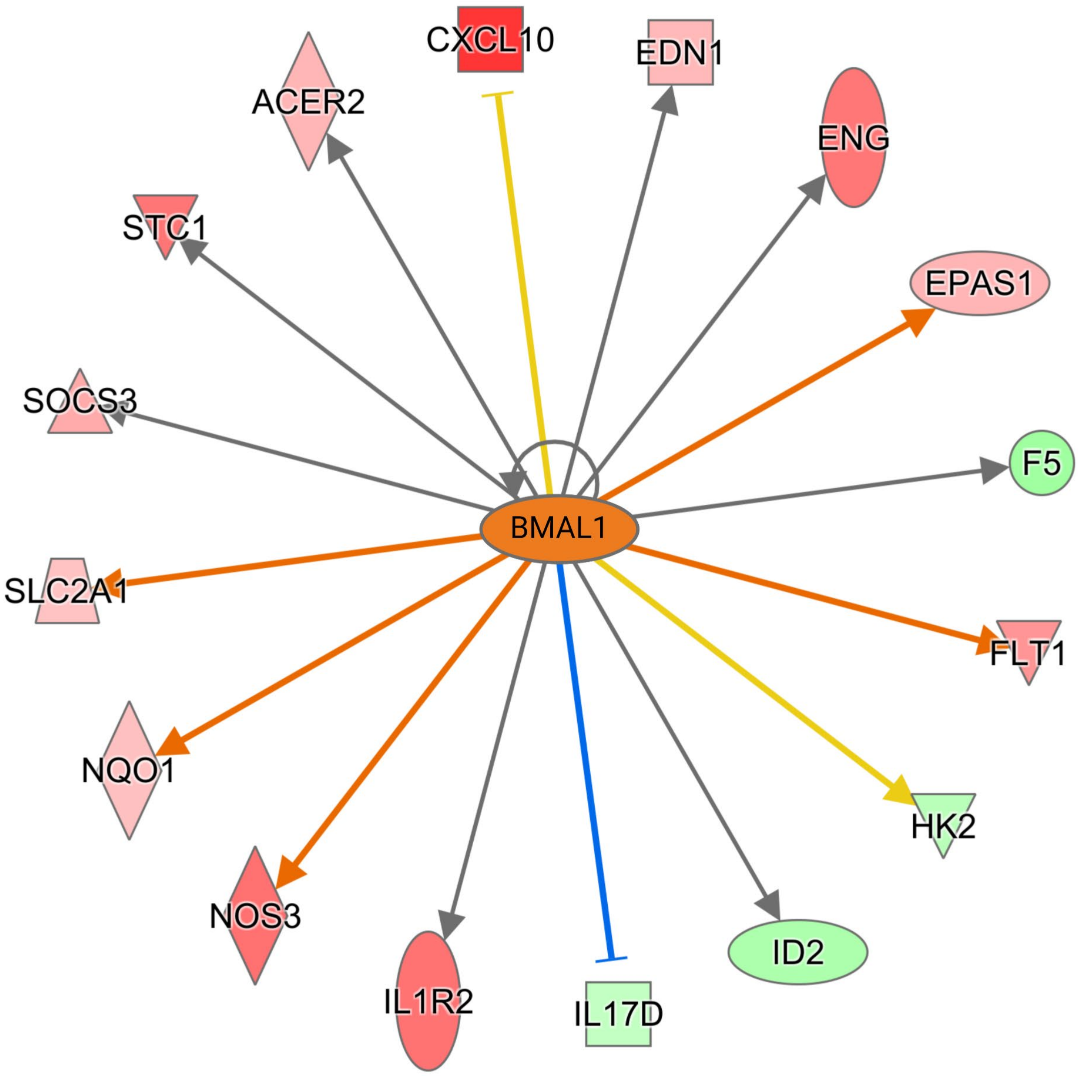


**Supplementary Figure S18. *Bmal1* regulates genes involved with cytokines signaling and cell defense in human mTLE-HS.** IPA analysis of *Bmal1* as an upstream regulator with corresponding activation/inhibition of predicted target genes in dissected hippocampal samples from human mTLE-HS. The activation/inhibition predicted in this analysis aligns with the RNAseq dataset. Red indicates upregulated genes, while green signifies downregulated; the intensity of colors (green and red) corresponds to the level of log2FC; weaker color indicates a lower expression level, while stronger color indicates a higher expression. Similarly, activation/inhibition of molecule activity, as defined by the activation z-score (orange lines) and inhibition z-score (blue lines), is depicted by color intensity: weaker color represents lower activation/inhibition level, whereas stronger color indicates higher activation/inhibition. Yellow lines represent inconsistency according to the literature; white represents molecules involved in the signaling pathway but absent in our dataset. Sharp arrows represent stimulation; blunt arrows represent inhibition; dashed line indicates indirect interaction.

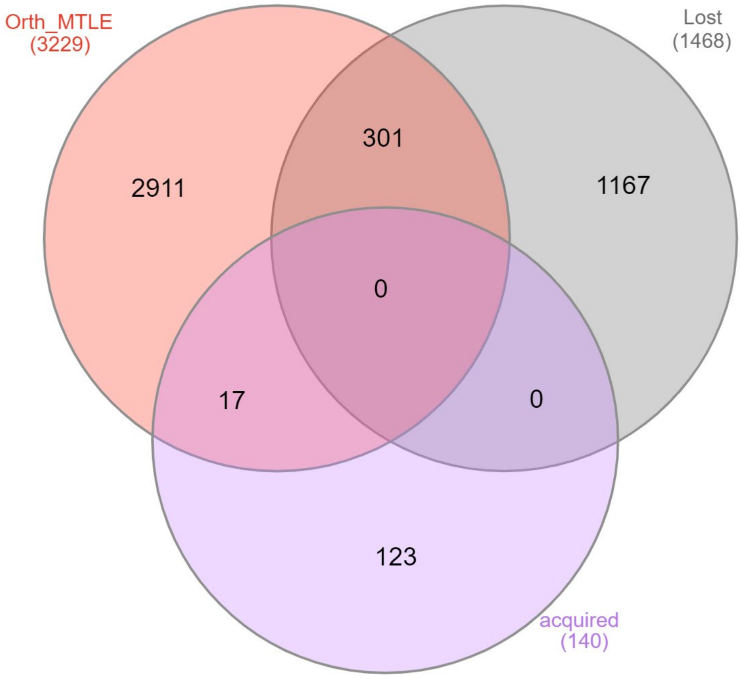


**Supplementary Figure S19. 17 orthologous genes in mTLE-HS overlapped with the ones that acquired rhythmicity, while 301 genes overlapped those that lost rhythmicity in experimental epilepsy.** An overlap analysis was performed between orthologues genes that were dysregulated in human mTLE-HS and genes that acquired or lost rhythmicity in mice hippocampus during epileptogenesis.

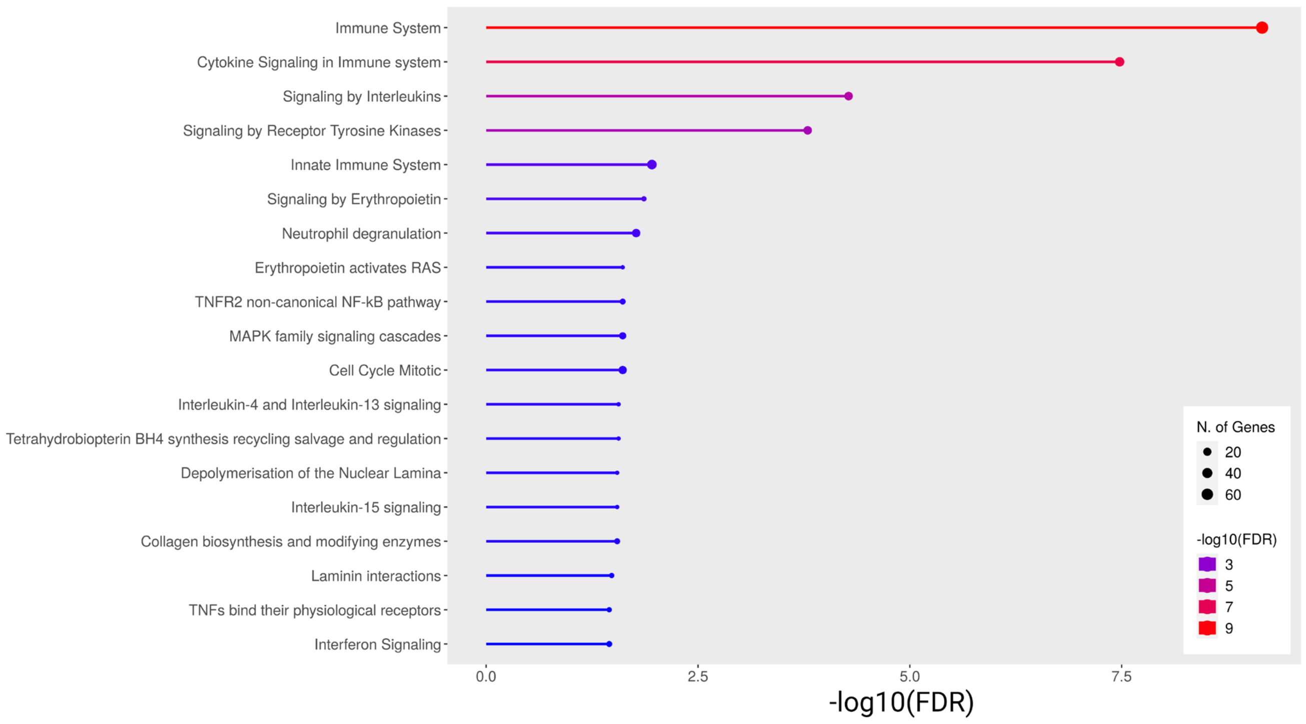


**Supplementary Figure S20. The Reactome analysis revealed the common relevant pathways in human mTLE-HS and experimental epileptogenesis.** The Reactome analysis was performed using the 668 DEGs (364 genes commonly upregulated and 304 downregulated) in both human mTLE-HS and experimental epileptogenesis RNAseq datasets.

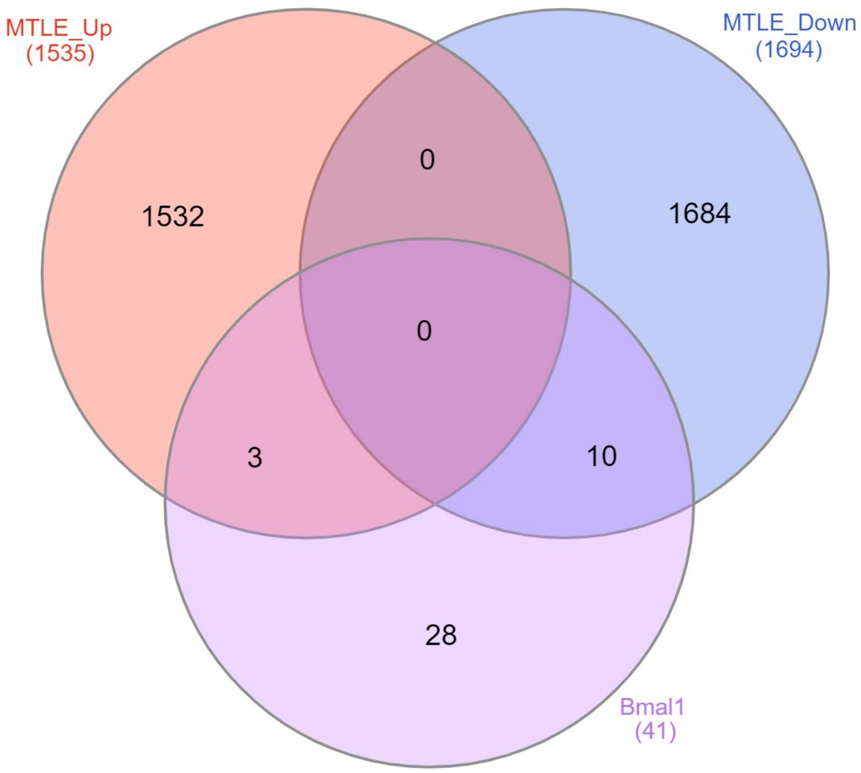


**Supplementary Figure S21. 3 upregulated genes and 10 downregulated genes in mTLE-HS were overlapping with the 41 *Bmal1*'s targets. An** overlap analysis was performed between the orthologues genes upregulated or downregulated in mTLE-HS patients and those identified as *Bmal1*'s target genes in mice hippocampus during epileptogenesis.

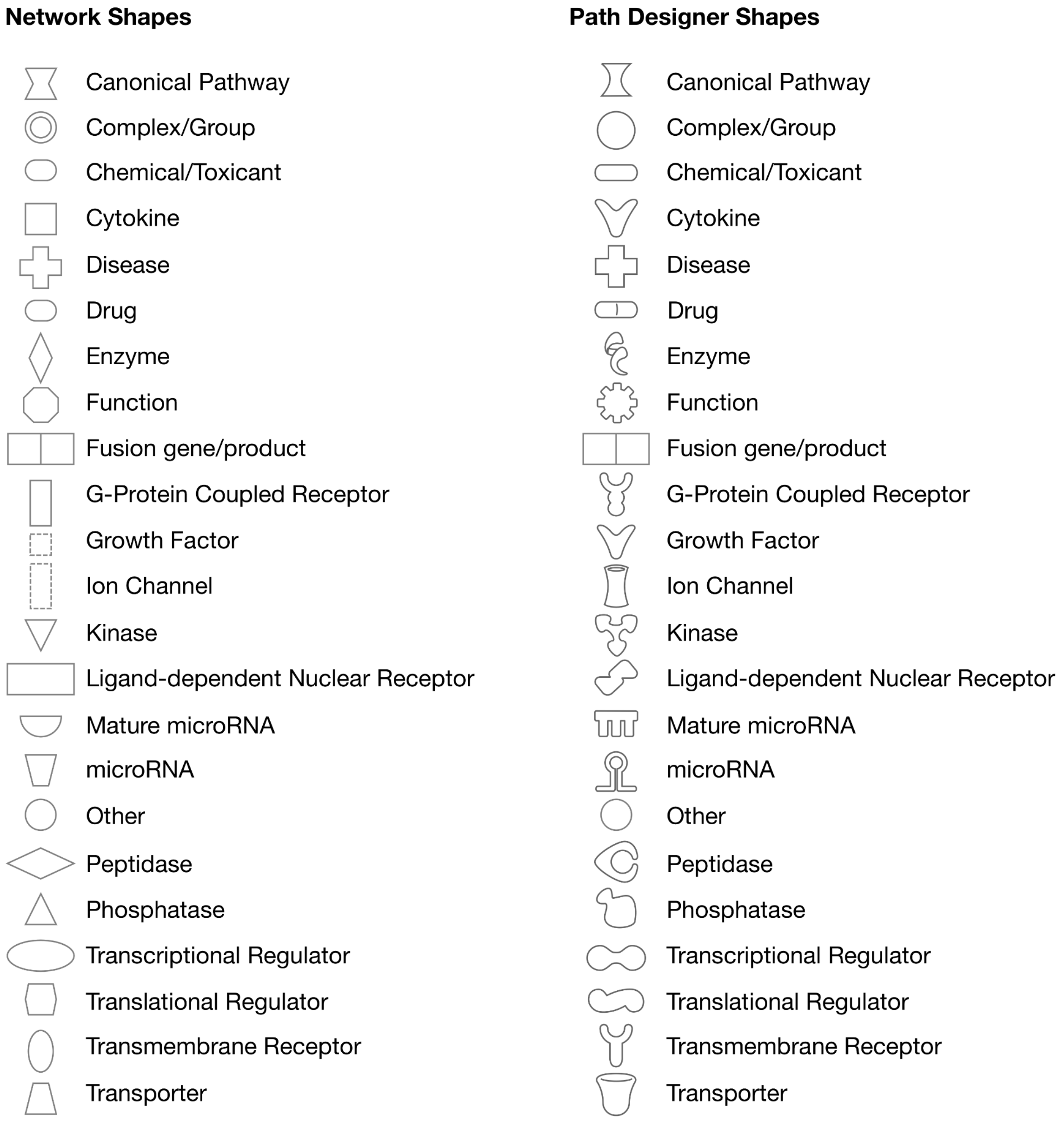


**Supplementary Figure S22.** Legend information for the shape of symbols that may appear in the IPA analysis figures.
