## Supplementary Tables for "EPILEPTOGENESIS INHIBITS THE CIRCADIAN CLOCK AND RESHAPES THE DIURNAL TRANSCRIPTOMIC RHYTHMICITY IN THE MOUSE HIPPOCAMPUS"

*Running title: Epileptogenesis reshapes the molecular rhythmicity in the mouse hippocampus*

Radharani Benvenutti[1,2], Danielle C. F. Bruno[3,4,5], Matheus Gallas-Lopes[6], Morten T. Venø[7], Estela Maria Bruxel[4,5], Tammy Strickland[1,2], Arielle Ramsook[1], Aditi Wadgaonkar[1], Yiyue Jiang[1,2], Amaya Sanz-Rodriguez[2,8], Lasse Sinkkonen[3], Marina K.M. Alvim[5,9], Clarissa L. Yasuda[5,9], Fabio Rogerio[5,10], Fernando Cendes[5,9], David C. Henshall[2,8], Annie M. Curtis[1,11], Katja Kobow[12], Iscia Lopes-Cendes[4,5], Cristina R. Reschke[1,2*]

[1] School of Pharmacy and Biomolecular Sciences, RCSI University of Medicine and Health Sciences, D02 YN77, Dublin, Ireland
[2] FutureNeuro SFI Research Centre, RCSI University of Medicine and Health Sciences, D02 YN77, Dublin, Ireland
[3] Department of Life Sciences and Medicine, Faculty of Science, Technology and Medicine, University of Luxembourg, 4367, Esch-Belval Esch-sur-Alzette, Luxembourg
[4] Department of Translational Medicine, School of Medical Sciences, University of Campinas (UNICAMP), 13083-888, Campinas, Brazil
[5] Brazilian Institute of Neuroscience and Neurotechnology (BRAINN), Brazil.
[6] Pharmacology Department, Federal University of Rio Grande do Sul, 90035-003, Porto Alegre, Brazil
[7] Omiics ApS, Aarhus, Denmark
[8] Department of Physiology and Medical Physics, RCSI University of Medicine and Health Sciences, D02 YN77, Dublin, Ireland
[9] Department of Neurology, School of Medical Sciences, University of Campinas (UNICAMP), 13083-888, Campinas, Brazil
[10] Department of Pathology, School of Medical Sciences, University of Campinas (UNICAMP), 13083-888, Campinas, Brazil
[11] Tissue Engineering Research Group (TERG), RCSI University of Medicine and Health Sciences, D02 YN77, Dublin, Ireland
[12] Department of Neuropathology, Universitätsklinikum Erlangen, Friedrich-Alexander University Erlangen-Nuremberg (FAU), Erlangen, Germany

*** Corresponding author:**
Cristina Ruedell Reschke, PhD
Chrono-Epilepsy Laboratory, School of Pharmacy and Biomolecular Sciences,
RCSI University of Medicine and Health Sciences, D02 YN77, Dublin, Ireland.


**Supplementary Table 1. The Top20 upregulated genes per ZT.** The identified genes' names and biological function in mice (*Mus musculus*) were determined based on the NCBI National Center for Biotechnology Information (https://www.ncbi.nlm.nih.gov/gene) tool. The log2Fold Change and p-values were extracted from the RNAseq dataset.

| Gene identified | ZT | log2Fold Change | p-value | Biological function |
|---|---|---|---|---|
| ***Gfap*** | 0 | 3.1500809 | 3.67E-39 | It is a constituent of the structural cytoskeleton. It is involved in the regulation of autophagy and acts upstream of several processes, including long-term synaptic potentiation, intermediate filament organization and neurogenesis. |
| ***Serpina3n*** | 0 | 4.4020706 | 4.92E-38 | It is a member of the serpin proteins family, responsible for serine proteases inhibition. |
| ***Nptx2*** | 0 | 3.0206063 | 4.91E-36 | Predicted to enable carbohydrate and metal ion binding and may be active in the glutamatergic synapse, expressed in the embryo, gut and the heart. |
| ***Slc14a1*** | 0 | 1.9354264 | 4.22E-33 | It encodes a membrane transporter that mediates urea and water transport in erythrocytes. |
| ***Sv2c*** | 0 | 2.5840129 | 5.24E-33 | Associated with transmembrane transporter activity. May be involved in chemical synaptic transmission, neurotransmitter transport, and transmembrane transport. |
| ***Fgfr1*** | 0 | 0.9687246 | 6.08E-32 | It encodes protein members of the fibroblast growth factor receptor family and is involved in forebrain development and regulation of the cell cycle. |
| ***Megf11*** | 0 | 1.9288319 | 1.26E-31 | It seems to play a role in the development of the retinal layers and homotypic cell-cell adhesion. Presumably located in the basolateral plasma membrane and may be a crucial part of the membrane. |
| ***Lrp8*** | 0 | 0.6331335 | 5.22E-30 | It encodes a member of the low density lipoprotein receptor family. The expressed protein enables binding activity for amyloid-beta and apolipoprotein and is involved in development of the nervous system and CREB regulation. |
| ***Bdnf*** | 0 | 1.5920165 | 8.25E-30 | It encodes a nerve growth factor family protein. It is involved in growth and differentiation of neurons in the developing nervous system. |
| ***Stk40*** | 0 | 1.5950376 | 8.94E-30 | It seems to enable ATP binding activity, protein serine kinase activity, and protein serine/threonine kinase activity. |
| ***Trib2*** | 0 | 1.7891013 | 2.07E-29 | It encodes one of the Tribbles family's three members. It is involved in the negative regulation of differentiation of fat cells and positively regulates catabolic protein processes. |

| | | | | |
|---|---|---|---|---|
| ***Vgf*** | 0 | 2.0249758<br>08 | 1.77E-27 | This gene, which is activated by nerve growth factor enables neuropeptide hormone activity which is involved in glucose homeostasis and insulin secretion. |
| ***C4b*** | 0 | 1.6602503<br>17 | 4.62E-27 | It is predicted to be involved in carbohydrate and complement component binding which mediates immunoglobulin immune responses. |
| ***Gldn*** | 0 | 4.4991453<br>85 | 4.08E-25 | It enables protein binding involved in cell-cell adhesion of heterotypic cells. It acts within clustering of voltage gated sodium channels and is involved in microvillus organisation. |
| ***Aldh1l2*** | 0 | 1.7669550<br>68 | 3.41E-26 | It encodes a member of the formyl transferase and aldehyde dehydrogenase superfamilies. This enzyme converts 10-formyltetrahydrofolate to tetrahydrofolate and $CO_2$ in a NADP(+)-dependent reaction and is the mitochondrial form of 10-formyltetrahydrofolate dehydrogenase. |
| ***Kif18a*** | 0 | 2.2115949 | 5.43E-25 | KIF18A belongs to the kinesin superfamily of microtubule-associated molecular motors, which employ ATP hydrolysis to generate force and movement along microtubules. |
| ***P4ha3*** | 0 | 3.3381435 | 8.37E-25 | It encodes a prolyl 4-hydroxylase component, a crucial enzyme in collagen production. The encoded protein is involved in peptidyl-proline hydroxylation to 4-hydroxy-L-proline. |
| ***Skil*** | 0 | 1.0083227 | 1.11E-25 | It encodes for a family of proteins involved in cellular response to extracellular growth signals. The encoded protein regulates growth factor beta signalling and is hence highly expressed in cancer cells. |
| ***Odc1*** | 0 | 2.1085110 | 1.88E-25 | Enables ornithine decarboxylase activity and protein homodimerization activity. Involved in putrescine biosynthetic process from ornithine and regulation of protein catabolic process. |
| ***Scube1*** | 0 | 1.7610603 | 4.06E-25 | This gene produces a SCUBE (signal peptide, CUB domain, EGF (epidermal growth factor)-like protein) family cell surface glycoprotein. This protein is expressed in endothelial cells and platelets, and it may be crucial for positive regulation of smoothened signalling pathway. |
| ***Gfap*** | 4 | 3.4166438 | 8.39E-46 | Already described in ZT0. |
| ***Serpina3n*** | 4 | 4.3947633 | 2.74E-38 | Already described in ZT0. |
| ***Nptx2*** | 4 | 3.0523869 | 3.45E-37 | Already described in ZT0. |
| ***Slc14a1*** | 4 | 2.0267802 | 6.56E-37 | Already described in ZT0. |
| ***Aldh1l2*** | 4 | 1.9691703 | 8.90E-33 | Already described in ZT0. |
| ***Gldn*** | 4 | 5.1962601 | 2.18E-32 | Already described in ZT0. |

| | | | | |
|---|---|---|---|---|
| ***Gbp2*** | 4 | 4.6454606 | 1.09E-30 | It belongs to the guanine-binding protein (GBP) family. It is predicted to enable Hsp90 protein and guanyl ribonucleotide binding which provides a cellular response to interferon beta. |
| ***S100a10*** | 4 | 2.7410069 | 1.20E-30 | Predicted to enable calcium-dependent protein binding and protein homodimerization. It is involved in the extrinsic component of the plasma membrane and is expressed in several organ systems. |
| ***C4b*** | 4 | 1.6985545 | 3.99E-29 | Already described in ZT0. |
| ***Tvp23a*** | 4 | 1.3342704 | 2.45E-24 | It encodes a membrane protein associated with the Golgi apparatus, which plays a crucial role in intracellular vesicular transport and protein secretion. |
| ***Sv2c*** | 4 | 2.3528175 | 6.94E-28 | Already described in ZT0. |
| ***Aqp4*** | 4 | 1.5360840 | 3.09E-27 | It encodes a member of the aquaporin family of intrinsic membrane proteins that function as water-selective channels in the plasma membranes of many cells. This protein is the predominant aquaporin found in the brain and has an important role in brain water homeostasis. |
| ***Stk40*** | 4 | 1.4664819 | 4.21E-26 | Already described in ZT0. |
| ***Ankrd28*** | 4 | 0.7107844 | 1.14E-25 | Not available. |
| ***Msn*** | 4 | 2.9974733 | 2.77E-25 | It is predicted to enable actin and double-stranded RNA binding which positively regulates the podosome assembly. |
| ***Wwtr1*** | 4 | 1.6892813 | 3.28E-25 | This gene encodes a protein which regulates cell cycle progression, differentiation and apoptosis. This protein is also acts as a transcriptional co-activator which binds to the PPXY motif. |
| ***Efhd2*** | 4 | 1.3147823 | 6.21E-25 | Enables cadherin binding activity. |
| ***Megf11*** | 4 | 1.6769055 | 7.93E-25 | Already described in ZT0. |
| ***Tubb2b*** | 4 | 1.4345832 | 2.47E-24 | It enables GTP binding and protein heterodimerization and is predicted to be a structural part of the cytoskeleton. |
| ***C4b*** | 8 | 1.9965143 | 4.95E-40 | Already described in ZT0. |
| ***Gfap*** | 8 | 3.1480151 | 3.76E-39 | Already described in ZT0. |
| ***Serpina3n*** | 8 | 4.3298321 | 3.86E-37 | Already described in ZT0. |
| ***Slc14a1*** | 8 | 1.8374734 | 1.20E-30 | Already described in ZT0. |
| ***Gbp2*** | 8 | 4.5528441 | 8.92E-30 | Already described in ZT4. |
| ***Parp3*** | 8 | 3.4308104 | 6.81E-29 | The protein encoded by this gene belongs to the PARP family. These enzymes modify nuclear proteins by poly-ADP-ribosylation, and is involved in the negative regulation of isotype switching. |
| ***Aldh1l2*** | 8 | 1.8473960 | 9.71E-29 | Already described in ZT0. |
| ***Aqp4*** | 8 | 1.4990315 | 4.91E-26 | Already described in ZT4. |
| ***Gldn*** | 8 | 4.3927440 | 2.30E-25 | Already described in ZT0. |

| | | | | |
|---|---|---|---|---|
| ***Fam111a*** | 8 | 3.1046910 | 6.16E-25 | It enables single-stranded DNA binding and is predicted to be involved in DNA metabolic processes and negative regulation of replication of the viral genome. |
| ***Nptx2*** | 8 | 2.4428702 | 2.10E-24 | Already described in ZT0. |
| ***Npas3*** | 8 | 1.0690024 | 1.13E-22 | The encoded protein is localized to the nucleus and may regulate protein heterodimerization and affect the startle response and other locomotory behaviour. |
| ***S100a10*** | 8 | 2.3136473 | 1.97E-22 | Already described in ZT4. |
| ***Msn*** | 8 | 2.7998579 | 2.63E-22 | Already described in ZT4. |
| ***Vim*** | 8 | 3.2088054 | 3.43E-22 | This gene enables RNA binding and is part of the cytoskeleton and eye lens structure. It is also involved in organisation of the intermediate filament and regulation of gene expression. It acts within astrocyte differentiation and cellular response to interferon-gamma. |
| ***Stat3*** | 8 | 1.9609594 | 3.87E-21 | It encodes a member of the STAT protein family. This protein is activated through phosphorylation in response to various cytokines and growth factors including IFNs, EGF, IL5, IL6, HGF, LIF, and BMP2. It also mediates the expression of a variety of genes in response to cell stimuli and plays a key role in many cellular processes such as cell growth and apoptosis. |
| ***Pros1*** | 8 | 1.9100276 | 5.61E-21 | It encodes a vitamin K-dependent plasma protein that is involved in coagulation, apoptosis and vasculogenesis. If this gene is not present, it can result in fulminant coagulopathy and death. |
| ***Col16a1*** | 8 | 2.1197971 | 4.36E-20 | Already described in ZT0. |
| ***Megf11*** | 8 | 1.5247302 | 1.80E-20 | Already described in ZT0. |
| ***Aspg*** | 8 | 3.9938469 | 2.78E-20 | Predicted to be involved in asparagine metabolic process and phospholipid metabolic process. |
| ***C4b*** | 12 | 1.9941400 | 3.20E-35 | Already described in ZT0. |
| ***Gfap*** | 12 | 3.1302837 | 1.32E-34 | Already described in ZT0. |
| ***Serpina3n*** | 12 | 4.2605049 | 5.45E-32 | Already described in ZT0. |
| ***Aldh1l2*** | 12 | 1.8445050 | 5.56E-25 | Already described in ZT0. |
| ***Gbp2*** | 12 | 4.5438872 | 5.78E-25 | Already described in ZT4. |
| ***Slc14a1*** | 12 | 1.7611588 | 3.68E-24 | Already described in ZT0. |
| ***Ly86*** | 12 | 2.4825380 | 1.06E-23 | Acts upstream of or within positive regulation of lipopolysaccharide-mediated signalling pathway. |
| ***Olfml3*** | 12 | 1.6491940 | 4.65E-23 | It is expressed in several organ systems such as the GI tract, genitourinary system and the neural ectoderm. |
| ***Parp3*** | 12 | 3.1212340 | 3.57E-22 | Already described in ZT8. |

| | | | | |
|---|---|---|---|---|
| ***Top2a*** | 12 | 4.0834101 | 1.81E-21 | It encodes a DNA topoisomerase, an enzyme that is involved female meiotic nuclear division and circadian rhythm regulation. |
| ***Gldn*** | 12 | 4.3751198 | 1.88E-21 | Already described in ZT0. |
| ***Vim*** | 12 | 3.3409208 | 2.03E-21 | Already described in ZT8. |
| ***Nptx2*** | 12 | 2.3679153 | 1.53E-20 | Already described in ZT0. |
| ***Gpnmb*** | 12 | 2.9137644 | 7.44E-20 | Enables heparin binding activity and integrin binding activity. Involved in several processes, including negative regulation of tumor necrosis factor production; positive regulation of ERK1 and ERK2 cascade; and positive regulation of protein phosphorylation. |
| ***Mki67*** | 12 | 3.8908982 | 1.26E-19 | Enables protein C-terminus binding activity. Involved in regulation of chromosome segregation and regulation of mitotic nuclear division. |
| ***C1qb*** | 12 | 1.8156083 | 7.90E-19 | It enables identical protein binding and is involved in synapse pruning which acts upstream of the development of the inner ear. |
| ***Msn*** | 12 | 2.6611368 | 3.59E-18 | Already described in ZT4. |
| ***S100a10*** | 12 | 2.1877156 | 3.93E-18 | Already described in ZT4. |
| ***Aqp4*** | 12 | 1.2987461 | 7.46E-18 | Already described in ZT4. |
| ***Mtmr11*** | 12 | 2.2132902 | 1.48E-17 | It seems to be involved in phosphatidylinositol dephosphorylation. |
| ***C4b*** | 16 | 1.9873579 | 6.24E-39 | Already described in ZT0. |
| ***Gfap*** | 16 | 2.8929298 | 2.62E-33 | Already described in ZT0. |
| ***Mki67*** | 16 | 4.6668661 | 3.68E-30 | Already described in ZT12. |
| ***Olfml3*** | 16 | 1.7376127 | 1.26E-28 | Already described in ZT12. |
| ***Top2a*** | 16 | 4.5115013 | 3.83E-28 | Already described in ZT12. |
| ***Serpina3n*** | 16 | 3.6388530 | 1.24E-26 | Already described in ZT0. |
| ***Parp3*** | 16 | 3.2951799 | 1.23E-24 | Already described in ZT8. |
| ***Vim*** | 16 | 3.2776047 | 4.93E-23 | Already described in ZT8. |
| ***Knl1*** | 16 | 3.5981281 | 8.48E-22 | Predicted to be involved in attachment of spindle microtubules to kinetochore and protein localization to kinetochore. Predicted to act upstream of or within cell division and chromosome segregation. |
| ***Laptm5*** | 16 | 1.1817494 | 2.18E-21 | It enables enzyme binding activity and protein sequestering and is involved in lysosomal transport and regulation of the macromolecule metabolic process. |
| ***Ly86*** | 16 | 2.1461845 | 2.80E-21 | Already described in ZT12. |
| ***Gpnmb*** | 16 | 2.7745918 | 3.78E-21 | Already described in ZT12. |
| ***Nusap1*** | 16 | 3.4173157 | 4.18E-21 | NUSAP1 is a nucleolar-spindle-associated protein that plays a role in spindle microtubule organization via localisation of the mitotic spindle. |
| ***Lgmn*** | 16 | 1.1818507 | 4.08E-20 | It encodes a member of the cysteine peptidase family which is involved in the endosome/lysosomal degradation system. The encoded protein undergoes autocatalytic removal of the C-terminal inhibitory peptide |

| | | | | |
|---|---|---|---|---|
| | | | | which generates the active endopeptidase to cleave the protein substrates on the asparagine residues. |
| ***Cd9*** | 16 | 1.5699501 | 5.34E-20 | It enables integrin binding, which is involve in skeletal muscle regeneration, paranodal junction assembly and single fertilisation. It also acts upstream of cell proliferation. |
| ***Aldh1l2*** | 16 | 1.5240174 | 1.08E-19 | Already described in ZT0. |
| ***Kif11*** | 16 | 2.5455673 | 1.38E-19 | It encodes a motor protein that belongs to the kinesin-like protein family, involved in various kinds of spindle dynamics including spindle assembly and possible colocalization with the spindle pole. |
| ***Gldn*** | 16 | 3.8427471 | 4.98E-19 | Already described in ZT0. |
| ***Bin2*** | 16 | 1.9214970 | 5.36E-19 | It enables phospholipid binding activity. Involved in several processes, including phagocytosis, engulfment; plasma membrane tubulation; and podosome assembly. |
| **Slc14a1** | 16 | 1.4513412 | 7.65E-19 | Already described in ZT0. |
| **C4b** | 20 | 2.1693282 | 1.16E-40 | Already described in ZT0. |
| **Gfap** | 20 | 3.0721792 | 2.24E-33 | Already described in ZT0. |
| **Olfml3** | 20 | 1.8191193 | 2.61E-27 | Already described in ZT12. |
| **Mki67** | 20 | 4.6446803 | 1.27E-26 | Already described in ZT12. |
| **Laptm5** | 20 | 1.4008313 | 3.50E-26 | Already described in ZT16. |
| **Serpina3n** | 20 | 3.7867513 | 1.32E-25 | Already described in ZT0. |
| **Ly86** | 20 | 2.5956213 | 1.02E-24 | Already described in ZT12. |
| **Vim** | 20 | 3.5297707 | 1.21E-23 | Already described in ZT8. |
| ***Bin2*** | 20 | 2.3403029 | 7.82E-23 | Already described in ZT16. |
| ***Top2a*** | 20 | 4.1773089 | 9.50E-23 | Already described in ZT12. |
| ***Lgmn*** | 20 | 1.2982995 | 2.09E-21 | Already described in ZT16. |
| ***Csf1r*** | 20 | 0.9552793 | 2.29E-21 | It encodes a receptor for colony stimulating factor 1, a cytokine which controls the production, differentiation, and function of macrophages. |
| ***Knl1*** | 20 | 3.8887635 | 4.55E-21 | Already described in ZT16. |
| ***Ctss*** | 20 | 1.4636258 | 7.03E-21 | The preproprotein encoded by this gene, a member of the peptidase C1 family generates enzyme secretion by antigen-presenting cells during inflammation. It is predicted to induce pain and itch via G-protein receptor activiation. |
| ***Nusap1*** | 20 | 3.8374753 | 1.31E-20 | Already described in ZT16. |
| ***Gpnmb*** | 20 | 2.9523067 | 3.30E-20 | Already described in ZT12. |
| ***Lair1*** | 20 | 0.9595944 | 1.35E-19 | Predicted to be in the membrane and and plasma membrane and may be an integral part of the membrane. Exact function is unknown |
| ***Aldh1l2*** | 20 | 1.6154823 | 2.39E-19 | Already described in ZT0. |
| ***Ccr5*** | 20 | 1.9032156 | 5.53E-19 | Enables C-C chemokine binding activity and C-C chemokine receptor activity. Involved in negative regulation of macrophage apoptosis. |

| | | | | |
|---|---|---|---|---|
| ***C1qc*** | 20 | 1.8643514 | 5.75E-19 | Involved in synapse pruning. Located in postsynapse. Is expressed in blood vessel and retina. |

**Supplementary Table 2. The Top20 downregulated genes per ZT.** The identified genes' names and biological function in mice (*Mus musculus*) were determined based on the NCBI National Center for Biotechnology Information (https://www.ncbi.nlm.nih.gov/gene) tool. The log2Fold Change and p-values were extracted from the RNAseq dataset.

| Gene identified | ZT | log2Fold Change | p-value | Biological function |
|---|---|---|---|---|
| ***Rps6ka5*** | 0 | -0.970493 | 5.60E-27 | Involved in a number of processes, including DNA-template transcription control, and protein phosphrylation. Permits protein serine/threonine kinase and ATP binding activity. |
| ***Aifm3*** | 0 | -1.318458 | 8.85E-26 | This gene is predicted to enable oxidoreductase which acts on NAD(P)H. It is also involved in the execution of apoptosis. |
| ***Cdh12*** | 0 | -1.133677 | 1.04E-25 | This gene encodes a member of the cadherin family of calcium-dependent glycoproteins that mediate cell adhesion and regulate many morphogenetic events during development. |
| ***Glul*** | 0 | -0.971055 | 9.20E-25 | It may enable anion binding, dynein light chain binding and metal ion binding. It is also involved in endothelial cell migration, sprouting angiogenesis and works upstream of the cellular response to both glucose and starvation. |
| ***Extl1*** | 0 | -1.178023 | 1.51E-24 | It seems to enable acetylglucosaminyl-transferase activity and glucuronosyl-transferase activity and may be involved in protein glycosylation. |
| ***Alcam*** | 0 | -0.881179 | 2.31E-24 | It seems to enable protein binding activity and acts upstream of motor neuron axon guidance. |
| ***Lzts3*** | 0 | -0.942842 | 5.19E-24 | It is thought to be involved in protein homooligomerization and the regulation of dendritic spine formation. It is active in the dendritic spine. |
| ***Lgi3*** | 0 | -0.921118 | 6.28E-24 | It is involved in catalytic activity and play a role in the regulation of exocytosis. |
| ***Hspa12a*** | 0 | -0.739589 | 1.90E-23 | It seems to enable ATP binding activity and is located in the extracellular exosome. |
| ***Jph4*** | 0 | -0.808120 | 5.07E-23 | It encodes a transmembrane protein from the junctophilin family, which is acts as an upstream of several processes, including learning; neuromuscular process controlling balance; and regulation of calcium entry. |
| ***Phyhip*** | 0 | -0.866918 | 2.46E-22 | Allows protein tyrosine kinase activity and is involved in protein localisation. |

| | | | | |
|---|---|---|---|---|
| ***Ppm1k*** | 0 | -0.592468 | 3.80E-22 | This gene is predicted to be involved in metal ion binding, protein serine phosphatase activity and protein threonine phosphatase activity. |
| ***Mturn*** | 0 | -0.935697 | 1.68E-21 | It positively regulates the differentiation of megakaryocytes and is expected to be found in the cytoplasm. |
| ***March1*** | 0 | -0.663883 | 6.32E-21 | Enables ubiquitin protein ligase activity. Involved in antigen processing and presentation of peptide antigen via MHC class II; immune response; and protein polyubiquitination. |
| ***Ivd*** | 0 | -0.721872 | 7.16E-21 | The mitochondrial matrix enzyme isovaleryl-CoA dehydrogenase catalyzes the third stage in leucine catabolism. |
| ***Amot*** | 0 | -0.860989 | 8.58E-21 | Predicted to enable angiostatin binding activity and signaling receptor activity. |
| ***Egfem1*** | 0 | -1.065736 | 1.39E-20 | Predicted to enable calcium ion binding activity. |
| ***Smarca2*** | 0 | -0.546901 | 2.04E-20 | It encodes a protein that enables several functions, including chromatin binding activity; transcription cis-regulatory region binding activity; and transcription coactivator activity. |
| ***Rasal1*** | 0 | -1.149095 | 3.16E-20 | It is expected to facilitate metal ion binding, phospholipid binding, and GTPase activator activity and may play a role in the regulation of dendritic extension positively. It is also expected to function either inside or ahead of cell differentiation. |
| ***Nsg2*** | 0 | -0.574935 | 3.26E-20 | It is expected to allow clathrin light chain binding activity and plays a role in clathrin coat assembly and endosomal trafficking. |
| ***Aifm3*** | 4 | -1.408897 | 5.13E-29 | Already described in ZT0. |
| ***Ivd*** | 4 | -0.737019 | 1.17E-21 | Already described in ZT0. |
| ***Pde7b*** | 4 | -1.415089 | 2.42E-21 | It permits the phosphodiesterase activity of 3',5'-cyclic AMP and is expected to play a role in signalling mediated by cAMP. |
| ***Sowaha*** | 4 | -1.044119 | 2.86E-21 | There is no specific known function. It is expressed in brain; liver lobe; and olfactory epithelium. |
| ***Alcam*** | 4 | -0.798526 | 3.55E-20 | Already described in ZT0. |
| ***Lzts3*** | 4 | -0.828192 | 7.77E-19 | Already described in ZT0. |
| ***Glul*** | 4 | -0.836822 | 8.48E-19 | Already described in ZT0. |
| ***Ppfia4*** | 4 | -0.635570 | 1.89E-18 | It is predicted to be involved in synapse organisation. |
| ***Nsg2*** | 4 | -0.546973 | 1.92E-18 | Already described in ZT0. |
| ***Appl2*** | 4 | -1.024823 | 2.24E-18 | It is predicted to facilitate the binding of small GTPases, proteins homodimerization, and phospholipids and is involved in the control of vesicle-mediated transport, the cellular response to hepatocyte growth factor stimulation, and the adiponectin-activated signalling pathway. |

| | | | | |
|---|---|---|---|---|
| ***Hspa12a*** | 4 | -0.646686 | 2.78E-18 | Already described in ZT0. |
| ***Jph4*** | 4 | -0.707230 | 5.44E-18 | Already described in ZT0. |
| ***Unc80*** | 4 | -0.549172 | 6.13E-18 | It enables cation channel activity and it is involved in cation homeostasis. |
| ***Efnb3*** | 4 | -0.929523 | 1.35E-17 | It is expected to facilitate the binding of ephrin receptors and is involved in regulating synaptic transmission through negative regulation of axonogenesis and trans-synaptic signalling by trans-synaptic complex. |
| ***Tprkb*** | 4 | -1.313157 | 2.35E-17 | It is involved in protein kinase binding activity and in tRNA threonylcarbamoyladenosine modification and telomere maintenance via recombination. |
| ***Lgi3*** | 4 | -0.762713 | 3.22E-17 | Already described in ZT0. |
| ***Ezh1*** | 4 | -0.599275 | 4.32E-17 | A member of the Polycomb-group (PcG) family is encoded by this gene. As a major element of the polycomb repressive complex 2 (PRC2), which methylates histone H3 at lysine 27 and causes the transcriptional suppression of impacted target genes, the encoded protein is interchangeable with the related Enhancer of zeste 2 (Ezh2) protein. |
| ***Abcc8*** | 4 | -1.047263 | 7.23E-17 | This protein regulates ATP-sensitive potassium channels and transmembrane transporter activity. It is predicted to be involved in learning and memory as well as negative regulation of blood-brain-barrier permeability. |
| ***Ralgapa2*** | 4 | -0.598307 | 1.10E-16 | It is expected to play a role in the activation of GTPase activity and may be involve in exocyst localisation. |
| ***Cdon*** | 4 | -0.811006 | 1.57E-16 | It functions either within or upstream of several activities, such as the growth of animal organs, the positive control of intracellular signal transduction, and the metabolic process of nitrogen compounds. |
| ***Glul*** | 8 | -0.877885 | 1.43E-20 | Already described in ZT0. |
| ***Appl2*** | 8 | -1.055409 | 2.16E-19 | Already described in ZT4. |
| ***Utp14b*** | 8 | -0.870280 | 3.74E-18 | It is thought to be involved in spermatogenesis and is part of small-subunit processome. |
| ***Ralgapa2*** | 8 | -0.593773 | 8.92E-17 | Already described in ZT4. |
| ***Tprkb*** | 8 | -1.272113 | 1.33E-16 | Already described in ZT4. |
| ***Gldc*** | 8 | -1.214399 | 3.84E-16 | It is involved in anion binding activity; glycine dehydrogenase (decarboxylating) activity; and protein homodimerization activity. |
| ***Aifm3*** | 8 | -0.991449 | 1.02E-15 | Already described in ZT0. |

| | | | | |
|---|---|---|---|---|
| ***Acsl3*** | 8 | -0.732361 | 3.17E-15 | Expected to facilitate protein kinase binding, protein domain specific binding, and arachidonate-CoA ligase activity. It may play a role in the metabolic process of fatty acids, positive regulation of the biosynthesis of phosphatidylcholine, positive regulation of transport and fatty acid metabolism process. |
| ***Pla2g7*** | 8 | -0.908399 | 1.13E-14 | Predicted to facilitate the activities of phospholipid binding, calcium-independent phospholipase A2, and 1-alkyl-2-acetylglycerophosphocholine esterase and may play a role in the remodelling of plasma lipoprotein particles, the glycerophospholipid catabolic process, and the positive control of monocyte chemotaxis. |
| ***Cyp2j9*** | 8 | -1.165757 | 1.88E-14 | Expected to facilitate the activities of monooxygenase, isomerase, and heme binding and may be involved in the metabolism of xenobiotics, linoleic acid, and the epoxygenase P450 pathway. |
| ***Mfge8*** | 8 | -1.075163 | 3.51E-14 | It allows for the binding activities of integrin, phosphatidylethanolamine, and phosphatidylserine and is involved in the removal of apoptotic cells and the processes of phagocytosis engulfment, phagocytosis, recognition, and positive phagocytosis control. |
| ***Cbx7*** | 8 | -0.652566 | 4.53E-14 | It enables chromatin, RNA and methylated histone binding and is invovled in sabecous gland development. |
| ***Rasd2*** | 8 | -0.997235 | 5.31E-14 | Predicted to enable several functions, including G-protein beta-subunit binding activity; GTP binding activity; and ubiquitin conjugating enzyme binding activity. Involved in dopaminergic synaptic transmission and locomotory behavior. |
| ***Ndrg2*** | 8 | -0.757332 | 7.18E-14 | This gene is involved in the regulation of the ERK1 and ERk2 cascade, smooth muscle cell proliferation and cytokine production. |
| ***Cldn10*** | 8 | -1.703698 | 9.35E-14 | This gene encodes a claudin family member. Tight junction strands act as a physical barrier to prevent solutes and water from freely moving through the paracellular gap between epithelial or endothelial cell sheets, and they also play important roles in cell polarity and signal transmission. |
| ***Nwd1*** | 8 | -1.012466 | 1.04E-13 | It is involved in ATP binding activity and regulation of NF-kB activity and positive regulation of gene expression. |
| ***Slc39a12*** | 8 | -1.281532 | 1.15E-13 | It encodes the protein that enables zinc ion transmembrane transport and regulation of microtubule polymerization and neuron projection development. |

| | | | | |
|---|---|---|---|---|
| ***Hlf*** | 8 | -0.900990 | 1.34E-13 | This gene enables DNA-binding transcription activation, RNA polymerase II- specific and cis-regulatory region, sequence specific DNA-binding activity. It allows for skeletal muscle cell differentiation. |
| ***Wasf3*** | 8 | -0.509016 | 2.07E-13 | It seems to enable Arp2/3 complex binding and protein kinase A regulatory subunit binding and is involved in postsynaptic modification of the actin cytoskeleton. |
| ***Tub*** | 8 | -0.547551 | 2.47E-13 | Predicted to enable G protein-coupled receptor binding activity and intraciliary transport particle A binding activity. Acts upstream of or within several processes, including phagocytosis, recognition; photoreceptor cell maintenance; and protein localization to non-motile cilium. |
| ***Ralgapa2*** | 12 | -0.570330 | 5.05E-14 | Already described in ZT4. |
| ***Lypd1*** | 12 | -0.945970 | 3.88E-12 | It is involved in the acetylcholine receptor signaling pathway. Several processes, including behavioral fear response, cholinergic synaptic transmission. |
| ***Camkk2*** | 12 | -0.633188 | 1.49E-11 | Predicted to enable calcium ion binding activity; calmodulin binding activity; and calmodulin-dependent protein kinase activity. Predicted to be involved in CAMKK-AMPK signaling cascade; activation of protein kinase activity; and protein autophosphorylation. |
| ***Erbb4*** | 12 | -0.648683 | 5.39E-11 | Expected to facilitate many activities, such as the binding of the ErbB-3 class receptor, the Hsp90 protein, and enzymes. It is also expected to support the binding activity of growth factors and is involved in both the positive control of protein phosphorylation and the activity of GTPase. |
| ***Mfge8*** | 12 | -0.961608 | 9.40E-11 | Already described in ZT8. |
| ***Dpf1*** | 12 | -0.492725 | 1.99E-10 | Allows for sequence-specific double-stranded DNA binding and transcription coregulator activity. It is also involved in nervous system development. |
| ***Otof*** | 12 | -2.022231 | 5.45E-10 | Enables AP-2 adaptor complex binding activity and calcium ion binding activity. Involved in synaptic vesicle priming. |
| ***Hlf*** | 12 | -0.797796 | 5.66E-10 | Already described in ZT8. |
| ***Trp53i11*** | 12 | -0.790880 | 8.71E-10 | It is predicted to be an integral component of the membrane and is expressed in the nervous system and neural retina. The exact function is unknown. |
| ***St8sia4*** | 12 | -0.559585 | 1.20E-09 | Enables alpha-N-acetylneuraminate alpha-2,8-sialyltransferase activity. Predicted to be involved in ganglioside biosynthetic process; glycoprotein metabolic process; and oligosaccharide metabolic process. |
| ***Glul*** | 12 | -0.603670 | 1.55E-09 | Already described in ZT0. |

| | | | | |
|---|---|---|---|---|
| ***Per3*** | 12 | -0.610468 | 1.82E-09 | This gene belongs to the Period family of genes and is expressed in the suprachiasmatic nucleus, the major circadian pacemaker in the mammalian brain, in a circadian pattern. This family of genes encodes components of circadian rhythms of locomotor activity, metabolism, and behavior. This gene is activated by CLOCK/ARNTL heterodimers, but it is then repressed in a feedback loop by PER/CRY heterodimers interacting with CLOCK/ARNTL. |
| ***Alcam*** | 12 | -0.533846 | 5.10E-09 | Already described in ZT0. |
| ***Aldh6a1*** | 12 | -0.563059 | 5.11E-09 | It encodes a protein from the aldehyde dehydrogenase family. The encoded protein is a mitochondrial methylmalonate semialdehyde dehydrogenase involved in the catabolic pathways of valine and pyrimidine. |
| ***Fat4*** | 12 | -0.643396 | 5.18E-09 | This gene is predicted to enable calcium ion binding and is involved in the heterophilic cell-cell adhesion via plasma membrane cell adhesion molecules. It also plays a part in hippocampus signalling and nervous system development. |
| ***Neurl1b*** | 12 | -0.698603 | 5.57E-09 | It is expected to allow ubiquitin protein ligase function. It is expected to play a role in ubiquitin-dependent endocytosis. It is found in the actin cytoskeleton and the cytoplasm. |
| ***Kcnk9*** | 12 | -0.620532 | 7.20E-09 | This gene enables the activity of voltage-gated potassium channels, allowing for potassium ion importation across the plasma membrane. |
| ***Nos1*** | 12 | -0.672388 | 7.88E-09 | This gene enables nitric-oxide synthase activity and is involved in several processes including the nitric oxide biosynthetic process, positive regulation of adenylate cyclase-activating adrenergic receptor signalling pathway and nitrogen compound metabolic processes. |
| ***Cntfr*** | 12 | -0.712336 | 1.02E-08 | It encodes the alpha subunit of the ciliary neurotrophic factor (CNTF) receptor which is responsible for the assembly of a trimolecular complex when bound to CNTF. The encoded protein also undergoes proteolytic processing to form a glycosylphosphatidylinositol-linked cell surface protein. |
| ***Nr1d2*** | 12 | -0.435223 | 1.02E-08 | It encodes a member of the nuclear hormone receptor family, specifically the NR1 receptor subfamily. The encoded protein is a transcriptional repressor that is involved in circadian rhythms regulation. |
| ***Ralgapa2*** | 16 | -0.505875 | 1.39E-12 | Already described in ZT4. |
| ***Per3*** | 16 | -0.628247 | 4.24E-11 | Already described in ZT12. |
| ***Rasd2*** | 16 | -0.847474 | 9.69E-11 | Already described in ZT8. |

| | | | | |
|---|---|---|---|---|
| ***Camkk2*** | 16 | -0.547192 | 8.34E-10 | Already described in ZT12.fat |
| ***Per2*** | 16 | -0.508918 | 1.39E-09 | This is a core clock gene that belongs to the Period family of genes it is activated by CLOCK/ARNTL heterodimers, but it is then repressed in a feedback loop by PER/CRY heterodimers interacting with CLOCK/ARNTL. |
| ***Tsc22d1*** | 16 | -0.176749 | 2.42E-08 | This gene encodes a leucine zipper transcription factor from the TSC22 domain family. It seems to enable DNA-binding transcription activator, RNA polymerase II-specific and cis-regulatory region, sequence specific DNA-binding activity. It is also regulates the apoptotic process and cell proliferation. |
| ***Mkx*** | 16 | -0.803805 | 2.61E-08 | It is predicted to enable DNA-binding transcription activator, RNA polymerase II-specific and cis-regulatory region, sequence specific DNA-binding activity. It is also involved RNA transcription regulation and tendon development. |
| ***Doc2b*** | 16 | -0.743070 | 2.87E-08 | The protein encoded by this gene is a calcium sensor that is involved in insulin secretion. |
| ***Adamts20*** | 16 | -0.495392 | 3.44E-08 | This gene encodes a member of a multi-domain matrix-associated metalloendopeptidases (ADAMTS) which are involved in tissue morphogenesis and pathophysiological remodelling in inflammation and general vascular biology. |
| ***Itpr1*** | 16 | -0.416508 | 4.42E-08 | This gene encodes an inositol 1,4,5-triphosphate intracellular receptor. This receptor mediates calcium release from the endoplasmic reticulum in response to inositol 1,4,5-triphosphate activation. |
| ***Kmt2a*** | 16 | -0.297782 | 5.06E-08 | It enables binding activity of DNA, chromatin and histone methyltransferase. It is also involved in haemopoiesis and regulation of gene expression and histone modification. |
| ***Gmps*** | 16 | -0.200890 | 8.12E-08 | Enables GMP synthase (glutamine-hydrolyzing) activity. Involved in GMP biosynthetic process. |
| ***Hlf*** | 16 | -0.645267 | 1.05E-07 | Already described in ZT8. |
| ***Atxn1*** | 16 | -0.291975 | 1.42E-07 | It enables RNA and chromatin binding and is involved in brain development, learning and memory and social behaviour. It also regulates insulin-like growth factor receptor signalling. |
| ***Fat4*** | 16 | -0.542354 | 1.73E-07 | Already described in ZT12. |
| ***Gldc*** | 16 | -0.745857 | 2.94E-07 | Already described in ZT8. |
| ***Elfn2*** | 16 | -0.348373 | 3.82E-07 | It enables protein phosphatase inhibitor activity. Predicted to be located in membrane and to be active in extracellular matrix and extracellular space. |

| | | | | |
|---|---|---|---|---|
| ***B3gat1*** | 16 | -0.350035 | 5.19E-07 | It enables UDP- galactose: beta-N-acetylglucosamine beta-1,3-galactosyltransferase activity and is involved carbohydrate metabolism and the chondroitin sulphate proteoglycan biosynthetic process. |
| ***Ppp1r13b*** | 16 | -0.341961 | 5.43E-07 | This gene increases DNA binding and transactivation of p53-family proteins on proapoptotic gene promoters. |
| ***Prdm8*** | 16 | -0.617642 | 6.47E-07 | It encodes a protein that enables chromatin binding and histone methyltransferase activity. It also acts within neuron axonogenesis and central nervous projection. |
| ***Ralgapa2*** | 20 | -0.561896 | 1.26E-13 | Already described in ZT4. |
| ***Fat4*** | 20 | -0.771786 | 2.76E-12 | Already described in ZT12. |
| ***Rasd2*** | 20 | -0.920033 | 3.62E-11 | Already described in ZT8. |
| ***Atp2b3*** | 20 | -0.660608 | 3.89E-11 | This gene encodes a protein that belongs to the P-type primary ion transport ATPase family. These enzymes are important in intracellular calcium homeostasis. |
| ***Ahcyl2*** | 20 | -0.547129 | 1.13E-10 | This gene enables hydrolase activity and in involved in S-adenosylmethionine cycle. It acts upstream of one-carbon metabolic processes. |
| ***Amer2*** | 20 | -0.538079 | 1.25E-10 | Allows for the binding of phosphatidylinositol-4,5-bisphosphate. Negatively regulates the canonical Wnt signaling pathway. |
| ***Ryr1*** | 20 | -0.896041 | 3.21E-10 | It enables protease binding and ryanodine-sensitive calcium-release channel activity. It is involved in animal organ development and the release of sequestered calcium ions into the cytosol. |
| ***Appl2*** | 20 | -0.777438 | 3.47E-10 | Already described in ZT4. |
| ***Fam160a1*** | 20 | -1.233639 | 4.62E-10 | Predicted to be involved in protein localization to perinuclear region of cytoplasm. |
| ***Kcng3*** | 20 | -0.954983 | 2.02E-09 | It enables delayed rectifier potassium channel activity and involved in potassium ion transmembrane transport. Predicted to act upstream of or within potassium ion transport. |
| ***Vwa3a*** | 20 | -1.481303 | 2.40E-09 | It is expressed in the $4^{th}$ ventricle, choroid plexus and lateral ventricle. The exact function is unknown |
| ***Mast3*** | 20 | -0.604044 | 4.86E-09 | Protein serine/threonine kinase activity is predicted. It is thought to be involved in the organization of the cytoskeleton, intracellular signal transduction, and peptidyl-serine phosphorylation. |
| ***Itpr1*** | 20 | -0.470595 | 5.60E-09 | Already described in ZT16. |

| | | | | |
|---|---|---|---|---|
| ***Ddn*** | 20 | -0.664282 | 8.36E-09 | DNA-binding transcription factor activity, RNA polymerase II-specific, and RNA polymerase II cis-regulatory region sequence-specific DNA binding activity are all predicted. It is predicted that RNA polymerase II will work upstream of or within the positive control of transcription. |
| ***Ecm2*** | 20 | -1.081347 | 1.04E-08 | It enables collagen V and heparin binding. It is also a positive regulator of cell-substrate adhesion. |
| ***Calb1*** | 20 | -0.811442 | 1.09E-08 | This gene enables calcium ion binding activity and is involved in the pre- and post-synaptic regulation of cytosolic calcium ion concentration and binding activity. |
| ***Aldh6a1*** | 20 | -0.550903 | 1.45E-08 | Already described in ZT12. |
| ***Grid1*** | 20 | -0.378266 | 1.46E-08 | This gene seems to allow for glutamate receptor activity, identical protein binding and transmitter-gated ion channel activity which regulates the postsynaptic membrane potential. |
| ***Tnxb*** | 20 | -0.822109 | 2.52E-08 | This gene enables collagen and heparin binding activity. It is involved in collagen fibril organisation, regulation of JUN kinase and triglyceride metabolism. |
| ***Glul*** | 20 | -0.554963 | 2.92E-08 | Already described in ZT0. |

**Supplementary Table 3. The Top100 genes that lost rhythmicity.** The identified genes' names and biological function in mice (*Mus musculus*) were determined based on the NCBI National Center for Biotechnology Information (https://www.ncbi.nlm.nih.gov/gene) tool.

| Gene identified | PBS meta2d_BH.Q | KA meta2d_BH.Q | Biological function |
|---|---|---|---|
| ***Gm26891*** | 1.89959E-06 | 1 | Not available. |
| ***Ppp1r14b*** | 1.89959E-06 | 1 | Predicted to enable protein serine/threonine phosphatase inhibitor activity. Predicted to be involved in innate immune response. |
| ***Atp6v0a4*** | 4.06545E-06 | 1 | Predicted to enable ATPase binding activity and P-type proton-exporting transporter activity. Acts upstream of or within proton transmembrane transport. |
| ***A830073O21Rik*** | 4.79775E-06 | 1 | Not available. |
| ***Arsj*** | 4.79775E-06 | 1 | Predicted to enable metal ion binding activity and sulfuric ester hydrolase activity. |
| ***Pcdhb19*** | 4.79775E-06 | 1 | Predicted to be involved in cell adhesion. |
| ***Sord*** | 4.79775E-06 | 1 | Enables L-iditol 2-dehydrogenase activity. Involved in flagellated sperm motility. Acts upstream of or within sorbitol metabolic process. |

| | | | |
|---|---|---|---|
| ***Rora*** | 4.91294E-06 | 1 | The protein encoded by this gene is a member of the NR1 subfamily of nuclear hormone receptors. It can bind as a monomer or as a homodimer to hormone response elements upstream of several genes to enhance the expression of those genes. The encoded protein has been shown to interact with NM23-2, a nucleoside diphosphate kinase involved in organogenesis and differentiation, as well as with NM23-1, the product of a tumor metastasis suppressor candidate gene. Also, it has been shown to aid in the transcriptional regulation of some genes involved in circadian rhythm. |
| ***Car8*** | 5.30967E-06 | 1 | Predicted to enable hydro-lyase activity. Acts upstream of or within phosphatidylinositol-mediated signaling. |
| ***Hs6st3*** | 5.30967E-06 | 1 | Enables heparan sulfate 6-O-sulfotransferase activity. Acts upstream of or within heparan sulfate proteoglycan biosynthetic process, enzymatic modification. |
| ***Mir669b*** | 5.30967E-06 | 1 | Not available. |
| ***Zc3h7b*** | 6.18375E-06 | 1 | Predicted to enable miRNA binding activity. Predicted to be involved in production of miRNAs involved in gene silencing by miRNA. |
| ***Use1*** | 8.27245E-06 | 1 | Predicted to enable SNAP receptor activity. Acts upstream of or within endoplasmic reticulum tubular network organization and regulation of ER to Golgi vesicle-mediated transport. Predicted to be part of SNARE complex. |
| ***Foxp4*** | 8.93842E-06 | 1 | Enables DNA-binding transcription repressor activity, RNA polymerase II-specific; RNA polymerase II cis-regulatory region sequence-specific DNA binding activity; and identical protein binding activity. Acts upstream of or within several processes, including embryonic foregut morphogenesis; lung secretory cell differentiation; and negative regulation of lung goblet cell differentiation. |
| ***Tmem164*** | 8.93842E-06 | 1 | Predicted to be located in membrane. Predicted to be integral component of membrane. |
| ***Zbtb39*** | 8.93842E-06 | 1 | Predicted to enable DNA-binding transcription repressor activity, RNA polymerase II-specific and RNA polymerase II cis-regulatory region sequence-specific DNA binding activity. Predicted to be involved in regulation of transcription by RNA polymerase II. |

| | | | |
|---|---|---|---|
| ***Mir434*** | 9.62578E-06 | 1 | Not available. |
| ***Axl*** | 1.12436E-05 | 1 | Predicted to enable several functions, including phosphatidylinositol 3-kinase binding activity; phosphatidylserine binding activity; and virus receptor activity. Involved in negative regulation of apoptotic process and positive regulation of protein kinase B signaling. Acts upstream of or within several processes, including animal organ development; myeloid cell homeostasis; and negative regulation of tumor necrosis factor production. |
| ***Fbn2*** | 1.12436E-05 | 1 | Predicted to be an extracellular matrix structural constituent. Involved in several processes, including bone trabecula formation; positive regulation of osteoblast differentiation; and sequestering of TGFbeta in extracellular matrix. Acts upstream of or within embryonic limb morphogenesis. |
| ***Gm48822*** | 1.12436E-05 | 1 | Not available |
| ***Bend6*** | 1.40987E-05 | 1 | Enables chromatin binding activity and transcription corepressor activity. Acts upstream of or within negative regulation of Notch signaling pathway and positive regulation of neuron differentiation. |
| ***Mme*** | 1.40987E-05 | 1 | Enables peptidase activity. Involved in amyloid-beta clearance; positive regulation of long-term synaptic potentiation; and sensory perception of pain. Acts upstream of or within amyloid-beta metabolic process. |
| ***Smyd3*** | 1.40987E-05 | 1 | Enables RNA polymerase II complex binding activity; RNA polymerase II transcription regulatory region sequence-specific DNA binding activity; and histone-lysine N-methyltransferase activity. Acts upstream of or within several processes, including cellular response to dexamethasone stimulus; nucleosome assembly; and regulation of protein phosphorylation. |
| ***Zfp553*** | 1.40987E-05 | 1 | Predicted to enable DNA-binding transcription activator activity, RNA polymerase II-specific; RNA polymerase II cis-regulatory region sequence-specific DNA binding activity; and identical protein binding activity. Predicted to be involved in regulation of transcription by RNA polymerase II. |
| ***Zfp783*** | 1.40987E-05 | 1 | Not available. |

| | | | |
|---|---|---|---|
| ***Ccdc85c*** | 1.42797E-05 | 1 | Involved in cerebral cortex development. Located in apical junction complex. |
| ***Id2*** | 1.45652E-05 | 1 | Enables RNA polymerase II-specific DNA-binding transcription factor binding activity and transcription regulator inhibitor activity. Involved in several processes, including animal organ development; entrainment of circadian clock by photoperiod; and regulation of gene expression. Acts upstream of with a negative effect on regulation of neural precursor cell proliferation and regulation of neuron differentiation. Acts upstream of or within several processes, including animal organ development; negative regulation of cell differentiation; and positive regulation of cell differentiation. |
| ***Pop4*** | 1.45652E-05 | 1 | Predicted to enable ribonuclease P RNA binding activity. Predicted to contribute to ribonuclease P activity. Predicted to be involved in rRNA processing and tRNA 5'-leader removal. |
| ***Tcf7l2*** | 1.45652E-05 | 1 | Enables several functions, including DNA-binding transcription activator activity, RNA polymerase II-specific; RNA polymerase II cis-regulatory region sequence-specific DNA binding activity; and beta-catenin binding activity. Involved in several processes, including positive regulation of insulin secretion; regulation of transcription, DNA-templated; and response to glucose. Acts upstream of or within several processes, including animal organ development; negative regulation of cellular response to growth factor stimulus; and regulation of cellular biosynthetic process. |
| ***Fech*** | 1.51263E-05 | 1 | Enables several functions, including ferrochelatase activity; heme binding activity; and iron-responsive element binding activity. Acts upstream of or within several processes, including porphyrin-containing compound metabolic process; regulation of cellular macromolecule biosynthetic process; and very-low-density lipoprotein particle assembly. |
| ***Fam186b*** | 1.53366E-05 | 1 | Predicted to be part of protein-containing complex. |
| ***Gpc5*** | 1.65555E-05 | 1 | Predicted to be involved in cell migration; positive regulation of canonical Wnt signaling pathway; and regulation of protein localization to membrane. |
| ***Smad7*** | 1.65555E-05 | 1 | Enables collagen binding activity. Involved in several processes, including circulatory |

| | | | |
|---|---|---|---|
| | | | system development; negative regulation of adaptive immune response based on somatic recombination of immune receptors built from immunoglobulin superfamily domains; and negative regulation of protein phosphorylation. Acts upstream of or within several processes, including cellular response to leukemia inhibitory factor; negative regulation of BMP signaling pathway; and transforming growth factor beta receptor signaling pathway. |
| ***1700113A16Rik*** | 1.77465E-05 | 1 | Not available. |
| ***Dnpep*** | 1.78878E-05 | 1 | Predicted to enable identical protein binding activity. Predicted to act upstream of or within proteolysis. |
| ***Mirlet7f-2*** | 1.8321E-05 | 1 | Not available. |
| ***Cav2*** | 2.00731E-05 | 1 | This gene belongs to the caveolin family whose members encode the major protein components of caveolae, which are invaginations of plasma membrane. This gene is located adjacent to caveolin-1 and the proteins coexpressed by the two genes localize together in caveolae, where they form hetero-oligomers. The encoded protein may be involved in diverse cellular functions including proliferation, differentiation, endocytosis and trafficking. |
| ***Hnrnpul1*** | 2.00731E-05 | 1 | Predicted to enable RNA binding activity and enzyme binding activity. Predicted to be involved in RNA processing and response to virus. |
| ***Gm32849*** | 2.20851E-05 | 1 | Not available. |
| ***Rad23a*** | 2.20851E-05 | 1 | Predicted to enable several functions, including enzyme binding activity; proteasome binding activity; and ubiquitin binding activity. Acts upstream of or within cellular response to DNA damage stimulus. |
| ***Klf9*** | 2.29055E-05 | 1 | Enables DNA-binding transcription activator activity, RNA polymerase II-specific and RNA polymerase II transcription regulatory region sequence-specific DNA binding activity. Involved in positive regulation of transcription by RNA polymerase II. Acts upstream of or within embryo implantation; progesterone receptor signaling pathway; and regulation of transcription, DNA-templated. |
| ***Slc25a17*** | 2.42963E-05 | 1 | Predicted to enable chaperone binding activity and nucleotide transmembrane transporter activity. Predicted to be |

| | | | |
|---|---|---|---|
| | | | involved in ATP transport; fatty acid beta-oxidation; and fatty acid transport. |
| ***Snhg3*** | 2.43194E-05 | 1 | Not available. |
| ***Ssbp3*** | 2.43194E-05 | 1 | This gene encodes a member of the Ssdp (sequence-specific single-stranded DNA binding protein) family of proteins. The encoded protein binds specifically to single-stranded pyrimidine-rich DNA elements. The encoded protein has been shown to be important for head development and may play a role in the differentiation of spinal interneurons. |
| ***Pcyt2*** | 2.50872E-05 | 1 | Enables ethanolamine-phosphate cytidylyltransferase activity. Involved in phosphatidylethanolamine biosynthetic process. |
| ***Txnip*** | 2.50872E-05 | 1 | Enables enzyme inhibitor activity. Acts upstream of or within several processes, including negative regulation of transcription by RNA polymerase II; platelet-derived growth factor receptor signaling pathway; and protein import into nucleus. |
| ***0610010F05Rik*** | 2.55545E-05 | 1 | Not available. |
| ***Abhd16a*** | 2.55545E-05 | 1 | Enables acylglycerol lipase activity and phospholipase activity. Involved in monoacylglycerol catabolic process; phosphatidylserine catabolic process; and prostaglandin catabolic process. |
| ***Hectd3*** | 2.57115E-05 | 1 | Enables syntaxin binding activity and ubiquitin-protein transferase activity. Predicted to be involved in proteasome-mediated ubiquitin-dependent protein catabolic process. |
| ***Ccdc85b*** | 2.60661E-05 | 1 | Predicted to enable delta-catenin binding activity. Involved in negative regulation of fat cell differentiation and negative regulation of transcription, DNA-templated. |
| ***Fdft1*** | 2.60661E-05 | 1 | Predicted to enable farnesyl-diphosphate farnesyltransferase activity and squalene synthase activity. Predicted to be involved in cholesterol biosynthetic process and farnesyl diphosphate metabolic process. Predicted to act upstream of or within cholesterol metabolic process; isoprenoid biosynthetic process; and sterol biosynthetic process. |
| ***Vmn2r85*** | 2.60661E-05 | 1 | Predicted to enable G protein-coupled receptor activity. |
| ***Jrkl*** | 2.64534E-05 | 1 | Predicted to enable DNA binding activity. |

| | | | |
|---|---|---|---|
| ***Gng3*** | 2.70222E-05 | 1 | Predicted to enable G-protein beta-subunit binding activity and GTPase activity. Predicted to be involved in G protein-coupled receptor signaling pathway and positive regulation of cytosolic calcium ion concentration. Predicted to act upstream of or within signal transduction. |
| ***Slc7a2*** | 2.70222E-05 | 1 | Enables L-amino acid transmembrane transporter activity. Involved in regulation of macrophage activation. Acts upstream of or within several processes, including L-alpha-amino acid transmembrane transport; nitric oxide biosynthetic process; and nitric oxide production involved in inflammatory response. |
| ***Usf2*** | 2.70222E-05 | 1 | Enables DNA binding activity. Involved in lipid homeostasis and positive regulation of transcription from RNA polymerase II promoter by glucose. Acts upstream of or within lactation; positive regulation of transcription, DNA-templated; and regulation of transcription by RNA polymerase II. |
| ***Gm15733*** | 2.75903E-05 | 1 | Not available. |
| ***Iqank1*** | 2.75903E-05 | 1 | Not available. |
| ***Kifc2*** | 2.75903E-05 | 1 | Predicted to enable ATP hydrolysis activity; microtubule binding activity; and microtubule motor activity. Predicted to be involved in microtubule-based movement and mitotic spindle assembly. |
| ***Mta1*** | 2.75903E-05 | 1 | Enables RNA polymerase II cis-regulatory region sequence-specific DNA binding activity and transcription coactivator activity. Involved in several processes, including circadian regulation of gene expression; entrainment of circadian clock by photoperiod; and locomotor rhythm. |
| ***Pak5*** | 2.75903E-05 | 1 | Enables protein serine/threonine kinase activity. Acts upstream of or within learning or memory; locomotory behavior; and negative regulation of extrinsic apoptotic signaling pathway. |
| ***Smarcd1*** | 2.75903E-05 | 1 | Predicted to enable several functions, including chromatin binding activity; molecular adaptor activity; and transcription coregulator activity. Predicted to be involved in epigenetic maintenance of chromatin in transcription-competent conformation; nucleosome disassembly; and regulation of transcription by RNA polymerase II. Predicted to act upstream of or within chromatin organization and nervous |

| | | | |
|---|---|---|---|
| | | | system development. Part of SWI/SNF complex; nBAF complex; and npBAF complex. |
| ***Vac14*** | 2.83222E-05 | 1 | Predicted to enable identical protein binding activity and kinase activator activity. Predicted to be involved in phosphatidylinositol biosynthetic process and positive regulation of kinase activity. Predicted to act upstream of or within response to osmotic stress. |
| ***Slc25a11*** | 2.8869E-05 | 1 | Predicted to enable antiporter activity; dicarboxylic acid transmembrane transporter activity; and sulfur compound transmembrane transporter activity. Predicted to be involved in anion transport. |
| ***Tm7sf2*** | 2.93896E-05 | 1 | Enables delta14-sterol reductase activity. Involved in cholesterol biosynthetic process. |
| ***Bcl2l1*** | 2.9635E-05 | 1 | This gene encodes a member of the Bcl-2 family of apoptosis regulators. The encoded protein is localized to the inner and outer mitochondrial membranes and regulates the programmed cell death pathway during development and tissue homeostasis. This protein binds to voltage-dependent anion channels in the outer mitochondrial membrane to facilitate the uptake of calcium ions. Mice embryos lacking this gene survived for two weeks and exhibited cell death of immature hematopoietic cells and neurons. |
| ***Dbp*** | 3.02127E-05 | 1 | The protein encoded by this gene is a member of the Par bZIP transcription factor family and binds to specific sequences in the promoters of several genes, such as albumin, Cyp2a4, and Cyp2a5. The encoded protein can bind DNA as a homo- or heterodimer and is involved in the regulation of some circadian rhythym genes. |
| ***Ralbp1*** | 3.02127E-05 | 1 | Enables GTPase activator activity and small GTPase binding activity. Involved in positive regulation of GTPase activity. |
| ***Slc30a1*** | 3.02127E-05 | 1 | Enables calcium channel inhibitor activity. Involved in cellular divalent inorganic cation homeostasis; negative regulation of ion transport; and zinc ion transport. Acts upstream of or within in utero embryonic development. |
| ***Umad1*** | 3.02127E-05 | 1 | Not available. |
| ***Gm45360*** | 3.0392E-05 | 1 | Not available. |
| ***Hip1r*** | 3.0392E-05 | 1 | Enables several functions, including SH3 domain binding activity; clathrin binding |

| | | | |
|---|---|---|---|
| | | | activity; and protein dimerization activity. Involved in several processes, including negative regulation of supramolecular fiber organization; positive regulation of clathrin coat assembly; and regulation of gastric acid secretion. Acts upstream of or within receptor-mediated endocytosis. |
| ***Lnx1*** | 3.0392E-05 | 1 | Enables PDZ domain binding activity; identical protein binding activity; and ubiquitin-protein transferase activity. Acts upstream of or within ubiquitin-dependent protein catabolic process. |
| ***Nt5c*** | 3.0392E-05 | 1 | Enables IMP 5'-nucleotidase activity. Involved in allantoin metabolic process and nucleoside monophosphate catabolic process. Acts upstream of or within deoxyribonucleotide catabolic process. |
| ***Timm23*** | 3.12551E-05 | 1 | Predicted to enable enzyme binding activity and protein transmembrane transporter activity. Predicted to be involved in protein import into mitochondrial matrix. Predicted to act upstream of or within protein transport. |
| ***Gm43663*** | 3.15047E-05 | 1 | Not available. |
| ***Gnai2*** | 3.15047E-05 | 1 | Predicted to enable several functions, including G-protein beta/gamma-subunit complex binding activity; GTP binding activity; and GTPase activity. Acts upstream of or within G protein-coupled acetylcholine receptor signaling pathway; adenylate cyclase-inhibiting G protein-coupled receptor signaling pathway; and cell population proliferation. |
| ***Grm2*** | 3.15047E-05 | 1 | Enables calcium channel regulator activity and group II metabotropic glutamate receptor activity. Acts upstream of or within chemical synaptic transmission; gene expression; and glutamate homeostasis. |
| ***H1f0*** | 3.15047E-05 | 1 | This gene is intronless and encodes a replication-independent histone that is a member of the histone H1 family. |
| ***Naxe*** | 3.15047E-05 | 1 | Enables identical protein binding activity. Predicted to be involved in several processes, including membrane raft distribution; negative regulation of angiogenesis; and regulation of cholesterol efflux. Predicted to act upstream of or within lipid transport. |
| ***Rala*** | 3.15047E-05 | 1 | Enables GTPase activity. Involved in membrane raft localization; neural tube closure; and regulation of exocytosis. |
| ***Rasa3*** | 3.15047E-05 | 1 | Predicted to enable calcium-release channel activity. Predicted to be involved |

| | | | |
|---|---|---|---|
| | | | in intracellular signal transduction; negative regulation of Ras protein signal transduction; and regulation of GTPase activity. Predicted to act upstream of or within cellular response to heat. |
| ***Tarsl2*** | 3.15047E-05 | 1 | Enables threonine-tRNA ligase activity. Involved in threonyl-tRNA aminoacylation. |
| ***Tnk2*** | 3.15047E-05 | 1 | Enables WW domain binding activity. Involved in spermatid development. |
| ***Rhou*** | 3.1546E-05 | 1 | Enables GTPase activity. Acts upstream of or within several processes, including G1/S transition of mitotic cell cycle; Rac protein signal transduction; and regulation of cell shape. |
| ***Camk2n1*** | 3.35572E-05 | 1 | Enables protein kinase inhibitor activity. Predicted to be involved in several processes, including negative regulation of ERK1 and ERK2 cascade; negative regulation of cellular protein metabolic process; and positive regulation of insulin secretion involved in cellular response to glucose stimulus. |
| ***Gm12104*** | 3.35572E-05 | 1 | Not available. |
| ***Mrps26*** | 3.37919E-05 | 1 | Not available. |
| ***Cdo1*** | 3.4497E-05 | 1 | Predicted to enable cysteine dioxygenase activity and ferrous iron binding activity. Predicted to be involved in several processes, including L-cysteine catabolic process; lactation; and response to glucagon. Predicted to act upstream of or within L-cysteine catabolic process to taurine. |
| ***Gm48415*** | 3.46252E-05 | 1 | Not available. |
| ***Gm45807*** | 3.47283E-05 | 1 | Not available. |
| ***Uvssa*** | 3.47283E-05 | 1 | Predicted to enable RNA polymerase II complex binding activity. Predicted to be involved in protein ubiquitination; response to UV; and transcription-coupled nucleotide-excision repair. Predicted to act upstream of or within DNA repair. Predicted to be located in nucleoplasm. |
| ***Zfp101*** | 3.47283E-05 | 1 | Predicted to enable DNA-binding transcription repressor activity, RNA polymerase II-specific and RNA polymerase II transcription regulatory region sequence-specific DNA binding activity. Predicted to be involved in negative regulation of transcription by RNA polymerase II. |
| ***Gm24453*** | 3.63496E-05 | 1 | Not available. |
| ***Maf1*** | 3.64585E-05 | 1 | Predicted to enable GABA receptor binding activity; RNA polymerase III cis-regulatory region sequence-specific DNA |

| | | | |
|---|---|---|---|
| | | | binding activity; and RNA polymerase III core binding activity. Involved in negative regulation of transcription by RNA polymerase III. |
| ***Tmem165*** | 3.65103E-05 | 1 | Predicted to enable calcium ion transmembrane transporter activity and manganese ion transmembrane transporter activity. Predicted to be involved in cellular cation homeostasis; metal ion transport; and protein N-linked glycosylation. |
| ***Trp53i11*** | 3.65451E-05 | 1 | Not available. |
| ***Tubg2*** | 3.69291E-05 | 1 | Predicted to enable GTP binding activity. Predicted to be a structural constituent of cytoskeleton. Predicted to be involved in microtubule cytoskeleton organization and mitotic sister chromatid segregation. |
| ***Scaper*** | 3.74084E-05 | 1 | Not available. |
| ***Sparcl1*** | 3.82358E-05 | 1 | Predicted to enable calcium ion binding activity; collagen binding activity; and extracellular matrix binding activity. Involved in synaptic membrane adhesion. |

**Supplementary Table 4. The Top100 genes that acquired rhythmicity.** The identified genes' names and biological function in mice (*Mus musculus*) were determined based on the NCBI National Center for Biotechnology Information (https://www.ncbi.nlm.nih.gov/gene) tool.

| **Gene identified** | **PBS meta2d_BH. Q** | **KA meta2d_BH. Q** | **Biological function** |
|---|---|---|---|
| ***Gm35552*** | 1 | 0.0000019 | Not available. |
| ***D030051J21Rik*** | 1 | 0.0000021 | Not available. |
| ***Gm40578*** | 1 | 0.0000029 | Not available. |
| ***Mas1*** | 1 | 0.000004 | Enables angiotensin receptor activity and peptide binding activity. Acts upstream of or within G protein-coupled receptor signaling pathway; activation of NF-kappaB-inducing kinase activity; and regulation of inflammatory response. |
| ***F2r*** | 1 | 0.0000375 | Enables G protein-coupled receptor activity; G-protein alpha-subunit binding activity; and G-protein beta-subunit binding activity. Involved in several processes, including negative regulation of renin secretion into blood stream; positive regulation of intracellular signal transduction; and trans-synaptic signaling by endocannabinoid, modulating synaptic transmission. Acts upstream of or within several processes, including regulation of interleukin-1 beta production; response to lipopolysaccharide; and thrombin-activated receptor signaling pathway. |

| | | | |
|---|---|---|---|
| ***Gm7967*** | 1 | 0.0000389 | Not available. |
| ***Cyp27a1*** | 1 | 0.0000389 | Predicted to enable Hsp70 protein binding activity; heme binding activity; and monooxygenase activity. Predicted to be involved in bile acid biosynthetic process; calcitriol biosynthetic process from calciol; and cholesterol catabolic process. Predicted to act upstream of or within steroid biosynthetic process. |
| ***Gldc*** | 1 | 0.000042 | Predicted to enable anion binding activity; glycine dehydrogenase (decarboxylating) activity; and protein homodimerization activity. Acts upstream of or within cellular response to leukemia inhibitory factor. |
| ***Gm10561*** | 1 | 0.0000448 | Not available. |
| ***Gm13322*** | 1 | 0.0000462 | Not available. |
| ***Cecr2*** | 1 | 0.0000462 | Predicted to enable ATP-dependent chromatin remodeler activity. Acts upstream of or within inner ear development; primary neural tube formation; and single fertilization. |
| ***4833421K07Rik*** | 1 | 0.000048 | Not available. |
| ***Olfr56*** | 1 | 0.000048 | Olfactory receptors interact with odorant molecules in the nose, to initiate a neuronal response that triggers the perception of a smell. |
| ***Gstk1*** | 1 | 0.0000598 | Enables glutathione peroxidase activity and glutathione transferase activity. Acts upstream of or within glutathione metabolic process. |
| ***1110002E22Rik*** | 1 | 0.000062 | Not available. |
| ***Gm43759*** | 1 | 0.0000664 | Not available. |
| ***Dsg1c*** | 1 | 0.0000823 | This gene encodes a member of the cadherin family of proteins that forms an integral transmembrane component of desmosomes, the multiprotein complexes involved in cell adhesion, organization of the cytoskeleton, cell sorting and cell signaling. |
| ***Lrriq3*** | 1 | 0.0001063 | Not available. |
| ***Nr4a3*** | 1 | 0.0001089 | This gene encodes a member of the NR4A subfamily of nuclear hormone receptors that bind to DNA and modulate gene expression. The encoded protein has been implicated in T and B lymphocyte apoptosis, and immune cell proliferation. Mice lacking the encoded protein exhibit partial bidirectional circling behavior and inner ear dysfunction. Disruption of this gene in mice also results |

| | | | |
|---|---|---|---|
| | | | in defective hippocampal axonal growth and postnatal neuronal cell death. |
| ***Gm27004*** | 1 | 0.0001115 | Not available. |
| ***Gm49310*** | 1 | 0.0001166 | Not available. |
| ***Gm48678*** | 1 | 0.0001337 | Not available. |
| ***Gm47097*** | 1 | 0.0001415 | Not available. |
| ***A830008E24Rik*** | 1 | 0.0001692 | Not available. |
| ***Gm11266*** | 1 | 0.0001692 | Not available. |
| ***Krt23*** | 1 | 0.0001692 | Predicted to enable structural molecule activity. |
| ***Sdc1*** | 1 | 0.0001692 | Predicted to enable identical protein binding activity and protein C-terminus binding activity. Involved in myoblast development. Acts upstream of or within canonical Wnt signaling pathway. |
| ***Dmc1*** | 1 | 0.0001692 | This gene encodes a member of the superfamily of recombinases (also called DNA strand-exchange proteins). Recombinases are important for repairing double-strand DNA breaks during mitosis and meiosis. This protein, which is evolutionarily conserved, is reported to be essential for meiotic homologous recombination and may thus play an important role in generating diversity of genetic information. In mouse, deficiency of this gene causes infertility. |
| ***Gm6145*** | 1 | 0.0001841 | Not available. |
| ***Slc44a3*** | 1 | 0.000185 | Predicted to enable transmembrane transporter activity. Predicted to be involved in transmembrane transport. |
| ***Atf3*** | 1 | 0.0002277 | Enables DNA-binding transcription repressor activity, RNA polymerase II-specific and RNA polymerase II cis-regulatory region sequence-specific DNA binding activity. Involved in several processes, including cellular response to amino acid starvation; negative regulation of ERK1 and ERK2 cascade; and regulation of gene expression. Acts upstream of or within gluconeogenesis; negative regulation of transcription, DNA-templated; and skeletal muscle cell differentiation. |
| ***Ccng2*** | 1 | 0.0002492 | Predicted to enable cyclin-dependent protein serine/threonine kinase regulator activity. Acts upstream of or within regulation of cell cycle. |
| ***1700019A02Rik*** | 1 | 0.0002674 | Not available. |
| ***Gm38059*** | 1 | 0.0002918 | Not available. |

| | | | |
|---|---|---|---|
| ***Pxylp1*** | 1 | 0.0002994 | Predicted to enable phosphatase activity. Predicted to be involved in several processes, including chondroitin sulfate proteoglycan biosynthetic process; glycosaminoglycan biosynthetic process; and positive regulation of heparan sulfate proteoglycan biosynthetic process. |
| ***Traip*** | 1 | 0.0003004 | Enables ubiquitin protein ligase activity. Involved in negative regulation of interferon-beta production and protein ubiquitination. Acts upstream of or within negative regulation of NF-kappaB transcription factor activity and signal transduction. |
| ***Gm5089*** | 1 | 0.0003246 | Not available. |
| ***Gm29508*** | 1 | 0.0003437 | Not available. |
| ***Lct*** | 1 | 0.0004021 | Predicted to enable lactase activity and transferase activity. Predicted to be involved in several processes, including response to iron(II) ion; response to lead ion; and response to nickel cation. |
| ***Tprkb*** | 1 | 0.0004174 | Predicted to enable protein kinase binding activity. Predicted to be involved in tRNA threonylcarbamoyladenosine modification and telomere maintenance via recombination. Predicted to act upstream of or within tRNA processing. |
| ***Prss23*** | 1 | 0.0004201 | Predicted to enable serine-type endopeptidase activity. Predicted to be involved in proteolysis. |
| ***Gm18284*** | 1 | 0.0004201 | Not available. |
| ***Kif18a*** | 1 | 0.0004371 | Predicted to enable several functions, including ATP hydrolysis activity; cytoskeletal protein binding activity; and plus-end-directed microtubule motor activity. Acts upstream of or within nuclear division; regulation of microtubule cytoskeleton organization; and seminiferous tubule development. |
| ***Tll1*** | 1 | 0.0004464 | Enables collagen binding activity. Predicted to be involved in proteolysis. Predicted to act upstream of or within cell differentiation. |
| ***Gli2*** | 1 | 0.0004717 | Enables DNA-binding transcription activator activity, RNA polymerase II-specific; RNA polymerase II cis-regulatory region sequence-specific DNA binding activity; and promoter-specific chromatin binding activity. Acts upstream of or within several processes, including animal organ development; chordate embryonic development; and regulation of smoothened signaling pathway. |

| | | | |
|---|---|---|---|
| ***Raver2*** | 1 | 0.0005153 | Predicted to enable mRNA binding activity. Predicted to be involved in regulation of alternative mRNA splicing, via spliceosome. |
| ***Gm49526*** | 1 | 0.000558 | Not available. |
| ***Trpc6*** | 1 | 0.0006377 | Enables store-operated calcium channel activity. Acts upstream of or within positive regulation of cytosolic calcium ion concentration. |
| ***Nrn1*** | 1 | 0.0006377 | Enables identical protein binding activity. Involved in regulation of synaptic vesicle exocytosis. Acts upstream of or within neuron projection extension. |
| ***Hsp25-ps1*** | 1 | 0.000641 | Not available. |
| ***Batf3*** | 1 | 0.0007461 | Predicted to enable DNA-binding transcription repressor activity, RNA polymerase II-specific and RNA polymerase II cis-regulatory region sequence-specific DNA binding activity. Involved in myeloid dendritic cell differentiation and response to virus. |
| ***Gm47138*** | 1 | 0.0007733 | Not available. |
| ***Upk1b*** | 1 | 0.0008428 | Acts upstream of or within response to bacterium. |
| ***Echdc2*** | 1 | 0.0009435 | Predicted to enable enoyl-CoA hydratase activity. Predicted to be involved in fatty acid beta-oxidation. Predicted to act upstream of or within fatty acid metabolic process. |
| ***Gm48940*** | 1 | 0.0009435 | Not available. |
| ***Lrrc10b*** | 1 | 0.0009713 | Not available. |
| ***Gm18955*** | 0.941076904 | 0.0001921 | Not available. |
| ***Gm17473*** | 0.908772533 | 0.0000021 | Not available. |
| ***Gm19967*** | 0.892268929 | 0.0000562 | Not available. |
| ***Gm14565*** | 0.886425733 | 0.0009109 | Not available. |
| ***Kcng3*** | 0.793476803 | 0.0009435 | Predicted to enable delayed rectifier potassium channel activity. Predicted to be involved in potassium ion transmembrane transport. Predicted to act upstream of or within potassium ion transport. |
| ***Rhbdf1*** | 0.780148355 | 0.0004716 | Predicted to be involved in several processes, including negative regulation of protein secretion; regulation of epidermal growth factor receptor signaling pathway; and regulation of proteasomal protein catabolic process. |
| ***Adam7*** | 0.747079423 | 0.0003088 | This gene encodes a member of a disintegrin and metalloprotease (ADAM) family of endoproteases that play important roles in various biological |

| | | | |
|---|---|---|---|
| | | | processes including cell signaling, adhesion and migration. |
| ***Pcsk6*** | 0.666247503 | 0.0009435 | Predicted to enable heparin binding activity; nerve growth factor binding activity; and serine-type endopeptidase activity. Acts upstream of or within determination of left/right symmetry; protein processing; and zygotic determination of anterior/posterior axis, embryo. |
| ***Tgtp2*** | 0.635487589 | 0.0004716 | Predicted to enable GTPase activity. Predicted to be involved in cellular response to interferon-beta and defense response. |
| ***Acsf2*** | 0.603891898 | 0.0008572 | Predicted to enable medium-chain fatty acid-CoA ligase activity. Predicted to be involved in fatty acid metabolic process. |
| ***Susd5*** | 0.600241597 | 0.0008237 | Acts upstream of or within Notch signaling pathway. |
| ***Crlf1*** | 0.564918873 | 0.0001692 | This gene encodes a member of the cytokine type I receptor family. The encoded protein functions as a cytokine receptor subunit and may be involved in immune system regulation and fetal development. |
| ***Mfap4*** | 0.557095771 | 0.0007747 | Predicted to enable antigen binding activity; carbohydrate derivative binding activity; and signaling receptor binding activity. Predicted to be involved in several processes, including complement activation, lectin pathway; elastic fiber assembly; and response to UV. Predicted to act upstream of or within cell adhesion. |
| ***Acot11*** | 0.539842822 | 0.0000956 | Predicted to enable long-chain acyl-CoA hydrolase activity; long-chain fatty acyl-CoA binding activity; and palmitoyl-CoA hydrolase activity. Involved in negative regulation of cold-induced thermogenesis. Acts upstream of or within response to cold. |
| ***Nat8f4*** | 0.52518713 | 0.0001769 | Predicted to enable cysteine-S-conjugate N-acetyltransferase activity and lysine N-acetyltransferase activity, acting on acetyl phosphate as donor. Predicted to be involved in several processes, including negative regulation of apoptotic process; peptide metabolic process; and peptidyl-lysine N6-acetylation. |
| ***Gm8188*** | 0.496916323 | 0.0000598 | Not available. |
| ***Gm43715*** | 0.46315953 | 0.0000534 | Not available. |
| ***A230009B12Rik*** | 0.382426743 | 0.0009435 | Not available. |

| | | | |
|---|---|---|---|
| ***Htra1*** | 0.380006899 | 0.0000304 | Enables serine-type peptidase activity. Involved in proteolysis. Acts upstream of or within chorionic trophoblast cell differentiation; negative regulation of transmembrane receptor protein serine/threonine kinase signaling pathway; and placenta development. |
| ***Fmnl1*** | 0.304523777 | 0.0003507 | Enables several functions, including GTPase activating protein binding activity; profilin binding activity; and small GTPase binding activity. Involved in actin filament severing. Acts upstream of or within substrate-dependent cell migration. |
| ***Skil*** | 0.276069183 | 0.0004201 | This gene encodes a member of a small family of proteins that play a key role in the response of cells to extracellular growth signals. The encoded protein regulates members of the transforming growth factor beta signaling pathway. It is highly expressed in certain cancer cells, where it may have both tumor-suppressing and tumor-promoting roles. |
| ***Anln*** | 0.241497222 | 0.000042 | Predicted to enable actin binding activity. Acts upstream of or within hematopoietic progenitor cell differentiation. |
| ***Ppp1r16b*** | 0.232038719 | 0.0000448 | Predicted to enable myosin phosphatase regulator activity and protein phosphatase 1 binding activity. Acts upstream of or within establishment of endothelial barrier. |
| ***Trerf1*** | 0.225313021 | 0.0003507 | Predicted to enable several functions, including nuclear receptor coactivator activity; progesterone receptor binding activity; and transcription coactivator binding activity. Predicted to be involved in cellular response to steroid hormone stimulus; histone deacetylation; and regulation of transcription, DNA-templated. |
| ***Cldn10*** | 0.214639584 | 0.0000304 | This intronless gene encodes a member of the claudin family. Claudins are integral membrane proteins and components of tight unction strands. |
| ***C2cd4c*** | 0.214714647 | 0.0006829 | Not available. |
| ***Gm44741*** | 0.209695294 | 0.0003931 | Not available. |
| ***Gjb6*** | 0.174080187 | 0.0000598 | Predicted to enable cytoskeletal protein binding activity and gap junction channel activity involved in cell communication by electrical coupling. Acts upstream of or within ear morphogenesis and sensory perception of sound. |
| ***Gm47033*** | 0.154100585 | 0.0003507 | Not available. |
| ***Paqr8*** | 0.149377933 | 0.0000223 | Predicted to enable steroid binding activity and steroid hormone receptor activity. Predicted to be involved in response to |

| | | | |
|---|---|---|---|
| | | | steroid hormone. Predicted to act upstream of or within oogenesis. |
| ***Pde8b*** | 0.11647962 | 0.0003931 | Enables 3',5'-cyclic-AMP phosphodiesterase activity. Acts upstream of or within several processes, including behavioral fear response; learning; and negative regulation of steroid hormone biosynthetic process. |
| ***Rftn2*** | 0.115912839 | 0.000004 | Acts upstream of or within dsRNA transport and response to exogenous dsRNA. |
| ***Gbp2*** | 0.111672236 | 0.0002447 | Predicted to enable several functions, including Hsp90 protein binding activity; cytoskeletal protein binding activity; and guanyl ribonucleotide binding activity. Acts upstream of or within several processes, including cellular response to interferon-beta; cellular response to lipopolysaccharide; and defense response to other organism. |
| ***Aldh1a1*** | 0.106130405 | 0.0000562 | Enables 3-chloroallyl aldehyde dehydrogenase activity and aminobutyraldehyde dehydrogenase activity. Involved in gamma-aminobutyric acid biosynthetic process and negative regulation of cold-induced thermogenesis. Acts upstream of or within several processes, including optic cup morphogenesis involved in camera-type eye development; positive regulation of apoptotic process; and retinoid metabolic process. |
| ***Nde1*** | 0.095846445 | 0.0001924 | Enables identical protein binding activity and microtubule binding activity. Involved in centrosome duplication and neuron migration. Acts upstream of or within microtubule nucleation; nervous system development; and vesicle transport along microtubule. |
| ***Phka1*** | 0.091444518 | 0.0005201 | Predicted to enable calmodulin binding activity. Predicted to be involved in protein autophosphorylation. Predicted to act upstream of or within glycogen metabolic process. |
| ***B230362B09Rik*** | 0.088004417 | 0.0007056 | Not available. |
| ***Cmbl*** | 0.087562499 | 0.0004201 | Predicted to enable hydrolase activity. |
| ***Slc4a4*** | 0.084247474 | 0.0000849 | Predicted to enable identical protein binding activity and sodium:bicarbonate symporter activity. Involved in regulation of intracellular pH. Acts upstream of or within bicarbonate transport and sodium ion transport. |

| | | | |
|---|---|---|---|
| ***Dio2*** | 0.064070513 | 0.0000159 | The protein encoded by this gene belongs to the iodothyronine deiodinase family. It catalyzes the conversion of prohormone thyroxine (3,5,3',5'-tetraiodothyronine, T4) to the bioactive thyroid hormone (3,5,3'-triiodothyronine, T3) by outer ring 5'-deiodination. |
| ***Cryab*** | 0.060906165 | 0.0009435 | This gene encodes a member of the small heat-shock protein (HSP20) family. The encoded protein is a molecular chaperone that protects proteins against thermal denaturation and other stresses. |
| ***H3c4*** | 0.057216332 | 0.000824 | This gene is intronless and encodes a replication-dependent histone that is a member of the histone H3 family. |
| ***Mir124a-1*** | 0.050413199 | 0.0002925 | Not available. |
| ***Cdh19*** | 0.046912397 | 0.0004662 | Predicted to enable cadherin binding activity and calcium ion binding activity. Predicted to be involved in calcium-dependent cell-cell adhesion via plasma membrane cell adhesion molecules; cell morphogenesis; and cell-cell junction organization. Predicted to be part of catenin complex. |

**Supplementary Table 5. The 21 genes that kept their diurnal rhythms in mice hippocampus during epileptogenesis.** A total of 21 genes identified in the RNAseq kept rhythmicity during epileptogenesis, wherein 19 presented an inverted rhythm in epileptogenesis (significant negative Pearson correlation), while just 2 genes maintained the same oscillating pattern in both groups (significant positive Pearson correlation). The identified genes' names and biological function in mice (*Mus musculus*) were determined based on the NCBI National Center for Biotechnology Information (https://www.ncbi.nlm.nih.gov/gene) tool.

| Gene identified | Pearson correlation | Gene name | Biological function |
|---|---|---|---|
| ***Slc25a18*** | −0.56 | solute carrier family 25 (mitochondrial carrier), member 18 | Predicted to enable L-aspartate transmembrane transporter activity and L-glutamate transmembrane transporter activity. Predicted to be involved in L-glutamate transmembrane transport; aspartate transmembrane transport; and malate-aspartate shuttle. |
| ***Myorg*** | −0.51 | myogenesis regulating glycosidase | Predicted to enable hydrolase activity, hydrolyzing O-glycosyl compounds. Acts upstream of or within positive regulation of insulin-like growth factor receptor signaling pathway; positive regulation of protein kinase B signaling; and skeletal muscle fiber development. |
| ***Ptprz1*** | −0.48 | protein tyrosine phosphatase receptor type Z, polypeptide 1 | Enables protein tyrosine phosphatase activity. Involved in several processes, including oligodendrocyte differentiation; peptidyl-tyrosine dephosphorylation; and regulation of myelination. Acts upstream of |

| | | | |
|---|---|---|---|
| | | | or within axonogenesis and hematopoietic progenitor cell differentiation. |
| ***Prrg1*** | −0.44 | proline rich Gla (G-carboxyglutamic acid) 1 | Not available. |
| ***Gja1*** | −0.43 | gap junction protein, alpha 1 | Enables several functions, including beta-tubulin binding activity; glutathione transmembrane transporter activity; and scaffold protein binding activity. Involved in several processes, including animal organ development; cellular response to amyloid-beta; and positive regulation of cold-induced thermogenesis. Acts upstream of or within several processes, including circulatory system development; regulation of gene expression; and regulation of membrane depolarization. |
| ***Nkain4*** | −0.43 | Na+/K+ transporting ATPase interacting 4 | Predicted to be involved in regulation of sodium ion transport. |
| ***Phgdh*** | −0.41 | 3-phosphoglycerate dehydrogenase | Predicted to enable phosphoglycerate dehydrogenase activity. Acts upstream of or within several processes, including G1 to G0 transition; cellular amino acid metabolic process; and nervous system development. |
| ***Ucp3*** | −0.41 | uncoupling protein 3 (mitochondrial, proton carrier) | Enables oxidative phosphorylation uncoupler activity. Acts upstream of or within fatty acid metabolic process and response to superoxide. |
| ***S1pr1*** | −0.36 | sphingosine 1-phosphate receptor 1 | This gene encodes a G-protein-coupled receptor bound by the lysophospholipid, sphingosine 1-phosphate. The gene product functions in endothelial cells and is involved in vascular and heart development. |
| ***Carmil1*** | −0.33 | capping protein regulator and myosin 1 linker 1 | Involved in several processes, including positive regulation of lamellipodium organization; positive regulation of substrate adhesion-dependent cell spreading; and positive regulation of supramolecular fiber organization. |
| ***Hmgcll1*** | −0.32 | 3-hydroxymethyl-3-methylglutaryl-Coenzyme A lyase-like 1 | Predicted to enable hydroxymethylglutaryl-CoA lyase activity. Predicted to be involved in ketone body biosynthetic process; leucine catabolic process; and lipid metabolic process. |
| ***Acsl3*** | −0.31 | acyl-CoA synthetase long-chain family member 3 | Predicted to enable arachidonate-CoA ligase activity; protein domain specific binding activity; and protein kinase binding activity. Predicted to be involved in several processes, including fatty acid metabolic process; positive regulation of |

| | | | |
|---|---|---|---|
| | | | phosphatidylcholine biosynthetic process; and positive regulation of transport. Predicted to act upstream of or within fatty acid metabolic process. |
| ***Cdh12*** | −0.31 | cadherin 12 | This gene encodes a member of the cadherin family of calcium-dependent glycoproteins that mediate cell adhesion and regulate many morphogenetic events during development. |
| ***Cdh20*** | −0.28 | cadherin 20 | Predicted to enable cadherin binding activity and calcium ion binding activity. Predicted to be involved in calcium-dependent cell-cell adhesion via plasma membrane cell adhesion molecules; cell morphogenesis; and cell-cell junction organization. |
| ***Utp14b*** | −0.28 | UTP14B small subunit processome component | Acts upstream of or within spermatogenesis. |
| ***Map7*** | −0.24 | microtubule-associated protein 7 | Enables signaling receptor binding activity. Involved in protein localization to plasma membrane and response to osmotic stress. Acts upstream of or within several processes, including gamete generation; male gonad development; and response to retinoic acid. |
| ***Selenop*** | −0.2 | selenoprotein P | Encodes a selenoprotein that contains multiple selenocysteine (Sec) residues per polypeptide and accounts for most of the selenium in plasma. It has been implicated as an extracellular antioxidant, and in the transport of selenium to extrahepatic tissues via apolipoprotein E receptor-2 (apoER2). Mice lacking this gene exhibit neurological dysfunction, suggesting its importance in normal brain function. |
| ***Gpt2*** | −0.11 | glutamic pyruvate transaminase (alanine aminotransfe-rase) 2 | Predicted to enable L-alanine:2-oxoglutarate aminotransferase activity. Predicted to be involved in 2-oxoglutarate metabolic process and L-alanine metabolic process. |
| ***Cyp2j9*** | −0.06 | cytochrome P450, family 2, subfamily j, polypeptide 9 | Predicted to enable heme binding activity; isomerase activity; and monooxygenase activity. Predicted to be involved in epoxygenase P450 pathway; linoleic acid metabolic process; and xenobiotic metabolic process. |
| ***Pnpla7*** | 0.09 | patatin-like phospholipase domain containing 7 | Enables lysophospholipase activity. Involved in phosphatidylcholine catabolic process. |

| | | | |
|---|---|---|---|
| ***Klf15*** | 0.2 | Kruppel-like transcription factor 15 | Enables DNA-binding transcription factor activity and transcription cis-regulatory region binding activity. Involved in cardiac muscle hypertrophy in response to stress; glomerular visceral epithelial cell differentiation; and negative regulation of peptidyl-lysine acetylation. Acts upstream of or within several processes, including cellular glucose homeostasis; positive regulation of glucose import; and response to insulin. |

**Supplementary Table 6. The main components identified in the neuroinflammation canonical pathway IPA analysis.** The identified genes' names and biological function in mice (*Mus musculus*) were determined based on the NCBI National Center for Biotechnology Information (https://www.ncbi.nlm.nih.gov/gene) tool.

| Code | Name | Biological function |
|---|---|---|
| ***Il34*** | interleukin 34 | It is a cytokine that promotes the differentiation and viability of monocytes and macrophages through colony-stimulating factor receptor-1 binding. |
| ***Csf1r*** | colony stimulating factor 1 receptor | Already described in Supplementary Table 1. |
| ***Ide*** | insulin degrading enzyme | Predicted to enable several functions, including ATP binding activity; ATP hydrolysis activity; and peptide binding activity. Involved in insulin catabolic process. Acts upstream of or within amyloid-beta clearance and response to oxidative stress. |
| ***Cx3cl1*** | C-X3-C motif chemokine ligand 1 | Enables CX3C chemokine receptor binding activity and chemokine activity. Acts upstream of or within several processes, including angiogenesis involved in wound healing; leukocyte chemotaxis; and negative regulation of extrinsic apoptotic signaling pathway in absence of ligand. |
| ***Cx3cr1*** | C-X3-C motif chemokine receptor 1 | Enables C-X3-C chemokine binding activity and C-X3-C chemokine receptor activity. Involved in several processes, including leukocyte migration; nervous system development; and synapse organization. Acts upstream of or within several processes, including macrophage chemotaxis; microglial cell activation involved in immune response; and negative regulation of extrinsic apoptotic signalling pathway in absence of ligand. |
| ***Cxcl10*** | C-X-C motif chemokine ligand 10 | Predicted to enable chemoattractant activity; chemokine receptor binding activity; and heparin binding activity. Acts upstream of or within several processes, including defense response to virus; negative regulation of |

| | | |
|---|---|---|
| | | myoblast differentiation; and negative regulation of myoblast fusion. |
| ***Cxcl8*** | C-X-C motif chemokine ligand 8 | The protein encoded by this gene is a member of the CXC chemokine family and is a major mediator of the inflammatory response. The encoded protein is commonly referred to as interleukin-8 (IL-8). IL-8 is secreted by mononuclear macrophages, neutrophils, eosinophils, T lymphocytes, epithelial cells, and fibroblasts. It functions as a chemotactic factor by guiding the neutrophils to the site of infection. IL-8 also participates with other cytokines in the proinflammatory signalling cascade and plays a role in systemic inflammatory response syndrome (SIRS). |
| ***Cxcl12*** | C-X-C motif chemokine ligand 12 | This gene encodes a member of the alpha chemokine protein family. The encoded protein is secreted and functions as the ligand for the G-protein coupled receptor, chemokine (C-X-C motif) receptor 4. The encoded protein plays a role in many diverse cellular functions, including embryogenesis, immune surveillance, inflammation response, tissue homeostasis, and tumor growth and metastasis. |
| ***Mfge8*** | milk fat globule EGF-factor 8 | Already described in Supplementary Table 2. |
| ***Jnk*** | mitogen-activated protein kinase 8 | Enables JUN kinase activity and protein serine/threonine/tyrosine kinase activity. Involved in several processes, including JNK cascade; generation of neurons; and positive regulation of cellular protein metabolic process. Acts upstream of or within several processes, including cellular response to nitric oxide; positive regulation of determination of dorsal identity; and protein phosphorylation. |
| ***Nfe2l2*** | nuclear factor erythroid 2-related factor 2 | Already described in Supplementary Table 4. |
| ***Hmox1*** | heme oxygenase 1 | Already described in Supplementary Table 4. |
| ***Pi3k*** | phosphoinositide-3-kinase | Enables insulin receptor substrate binding activity; phosphatidylinositol 3-kinase regulatory subunit binding activity; and protein heterodimerization activity. Involved in several processes, including negative regulation of stress fiber assembly; positive regulation of cellular component organization; and positive regulation of protein import into nucleus. Acts upstream of or within several processes, including apoptotic signalling pathway; negative regulation of osteoclast differentiation; and positive regulation of tumor necrosis factor production. |

| | | |
|---|---|---|
| ***Trem2*** | triggering receptor expressed on myeloid cells 2 | The protein encoded by this gene is part of the immunoglobulin and lectin-like superfamily and functions as part of the innate immune system. This protein associates with the adaptor protein Dap-12 and recruits several factors, such as kinases and phospholipase C-gamma, to form a receptor signalling complex that activates myeloid cells, including dendritic cells and microglia. |
| ***Syk*** | spleen associated tyrosine kinase | Enables several functions, including ATP binding activity; SH2 domain binding activity; and enzyme binding activity. Involved in several processes, including cell surface receptor signaling pathway; myeloid cell activation involved in immune response; and regulation of vesicle-mediated transport. Acts upstream of or within several processes, including positive regulation of leukocyte activation; positive regulation of macromolecule metabolic process; and protein phosphorylation. |
| ***Ip3*** | inositol 1,4,5-trisphosphate receptor type 1 | Enables inositol 1,4,5-trisphosphate receptor activity involved in regulation of postsynaptic cytosolic calcium levels; phosphatidylinositol binding activity; and protein domain specific binding activity. Involved in several processes, including epithelial fluid transport; intrinsic apoptotic signalling pathway in response to endoplasmic reticulum stress; and release of sequestered calcium ion into cytosol. Acts upstream of or within several processes, including calcium ion transport; endoplasmic reticulum calcium ion homeostasis; and voluntary musculoskeletal movement. |
| ***Nfat*** | nuclear factor of activated T-cells | Enables several functions, including DNA-binding transcription factor activity, RNA polymerase II-specific; RNA polymerase II cis-regulatory region sequence-specific DNA binding activity; and mitogen-activated protein kinase p38 binding activity. Involved in heart development and positive regulation of transcription by RNA polymerase II. Acts upstream of or within several processes, including animal organ development; branching involved in lymph vessel morphogenesis; and regulation of gene expression. |
| ***Akt*** | alpha serine threonine-protein kinase | This gene encodes the founding member of the Akt serine-threonine protein kinase gene family that also includes Akt2 and Akt3. This kinase is a major downstream effector of the phosphatidylinositol 3-kinase (PI3K) pathway that mediates the effects of various growth factors such as platelet-derived growth factor |

| | | |
|---|---|---|
| | | (PDGF), epidermal growth factor (EGF), insulin and insulin-like growth factor I (IGF-I). It plays a role in mediating a variety of cellular processes, such as glucose metabolism, glycogen biosynthesis, protein synthesis and turn over, inflammatory response, cell survival (anti-apoptosis) and development. |
| ***Cd200*** | CD200 | Predicted to enable protein binding activity involved in heterotypic cell-cell adhesion. Involved in negative regulation of cell population proliferation; negative regulation of macrophage activation; and negative regulation of neuroinflammatory response. |
| ***Stat1*** | signal transducer and activator of transcription 1 | Enables DNA-binding transcription factor activity. Involved in activation of cysteine-type endopeptidase activity involved in apoptotic process and defence response to other organisms. Acts upstream of or within several processes, including cell surface receptor signalling pathway; negative regulation of macrophage fusion; and response to exogenous dsRNA. |
| ***Ifng*** | interferon gamma | This gene encodes a soluble cytokine that is a member of the type II interferon class. The encoded protein is secreted by cells of both the innate and adaptive immune systems. The active protein is a homodimer that binds to the interferon gamma receptor which triggers a cellular response to viral and microbial infections. |
| ***Nos2*** | nitric oxide synthase 2 | Nitric oxide is a reactive free radical which acts as a biologic mediator in several processes, including neurotransmission and antimicrobial and antitumoral activities. This gene encodes a nitric oxide synthase which is expressed in liver and is inducible by a combination of lipopolysaccharide and certain cytokines. |
| ***Ifngr*** | interferon gamma receptor 1 | Predicted to enable cytokine receptor activity. Involved in several processes, including astrocyte activation; negative regulation of amyloid-beta clearance; and positive regulation of macromolecule metabolic process. Acts upstream of or within defence response to virus. Predicted to be integral component of plasma membrane. |
| ***Gsk3b*** | glycogen synthase kinase 3 β | Enables several functions, including beta-catenin binding activity; dynein complex binding activity; and protein kinase activity. Involved in several processes, including apoptotic signalling pathway; negative regulation of signal transduction; and regulation of gene expression. Acts upstream |

| | | |
|---|---|---|
| | | of or within several processes, including cellular response to hepatocyte growth factor stimulus; positive regulation of nitrogen compound metabolic process; and regulation of neuron projection development. |
| ***Il-6*** | interleukin 6 | This gene encodes a member of the interleukin family of cytokines that have important functions in immune response, hematopoiesis, inflammation and the acute phase response. The ectopic overexpression of the encoded protein in mice results in excessive plasma cells in circulation, leading to death. |
| ***Jak*** | janus kinase | Enables interleukin-12 receptor binding activity and protein tyrosine kinase activity. Involved in several processes, including hemopoiesis; positive regulation of macromolecule metabolic process; and regulation of signal transduction. Acts upstream of or within several processes, including cell surface receptor signaling pathway; cellular response to lipid; and regulation of apoptotic process. |
| ***Mapk*** | mitogen-activated protein kinase 1 | Enables several functions, including identical protein binding activity; phosphotyrosine residue binding activity; and protein kinase activity. Involved in peptidyl-serine phosphorylation; peptidyl-threonine phosphorylation; and response to nicotine. Acts upstream of or within several processes, including animal organ development; cell surface receptor signalling pathway; and cellular response to cytokine stimulus. |
| ***Nfkb*** | nuclear factor Kappa B | Enables RNA polymerase II cis-regulatory region sequence-specific DNA binding activity; chromatin binding activity; and identical protein binding activity. Involved in JNK cascade; cellular response to virus; and negative regulation of cytokine production. Acts upstream of or within several processes, including cellular response to dsRNA; cellular response to tumour necrosis factor; and regulation of gene expression. |
| ***Ap1*** | activator protein-1 heterodimeric transcription factor | Enables chromatin binding activity and transcription cis-regulatory region binding activity. Involved in cellular response to anisomycin and positive regulation of transcription from RNA polymerase II promoter involved in cellular response to chemical stimulus. Acts upstream of or within several processes, including animal organ development; negative regulation of protein autophosphorylation; and positive regulation of cell population proliferation. |

| | | |
|---|---|---|
| ***Creb*** | CREB binding protein | Enables DNA-binding transcription activator activity, RNA polymerase II-specific and cAMP response element binding activity. Involved in several processes, including cellular response to retinoic acid; positive regulation of biosynthetic process; and protein stabilization. Acts upstream of or within several processes, including animal organ development; cellular response to hepatocyte growth factor stimulus; and cellular response to leukemia inhibitory factor. |
| ***Ptgf2*** | Not available. | Not available. |
| ***Pge2*** | Not available. | Not available. |
| ***Cpla2*** | cytosolic phospholipase A2 | The protein encoded by this gene is a member of the phospholipase A2 group IV family. This enzyme hydrolyzes membrane phospholipids, thereby releasing the polyunsaturated fatty acid, arachidonic acid. Arachidonic acid is further metabolized into eicosanoids such as leukotrienes, thromboxanes and prostaglandins, that play important roles in regulating diverse biological processes such as inflammatory responses, membrane and actin dynamics, and tumorigenesis. |
| ***Mmp9*** | matrix metallopeptidase 9 | This gene encodes a member of the matrix metalloproteinase family of extracellular matrix-degrading enzymes that are involved in tissue remodeling, wound repair, progression of atherosclerosis and tumor invasion. The encoded preproprotein undergoes proteolytic processing to generate a mature, zinc-dependent endopeptidase enzyme that degrades collagens of type IV, V and XI, and elastin. |
| ***Ifng/ Ifngr*** | interferon gamma | Predicted to enable cytokine receptor activity. Involved in several processes, including astrocyte activation; negative regulation of amyloid-beta clearance; and positive regulation of macromolecule metabolic process. Acts upstream of or within defence response to virus. |
| ***Ager*** | advanced glycosylation end-product specific receptor | Enables S100 protein binding activity; advanced glycation end-product binding activity; and heparin binding activity. Involved in several processes, including cellular response to amyloid-beta; negative regulation of long-term synaptic potentiation; and positive regulation of cytokine production. Acts upstream of or within several processes, including induction of positive chemotaxis; negative regulation of advanced glycation end-product receptor activity; and positive |

| | | |
|---|---|---|
| | | regulation of macromolecule metabolic process. |
| ***Tgfβ*** | transforming growth factor beta | This gene encodes a secreted ligand of the TGF-beta (transforming growth factor-beta) superfamily of proteins. Ligands of this family bind various TGF-beta receptors leading to recruitment and activation of SMAD family transcription factors that regulate gene expression. This encoded protein regulates cell proliferation, differentiation and growth, and can modulate expression and activation of other growth factors including interferon gamma and Tnf- alpha. |
| ***Mmp3*** | matrix metallopeptidase 3 | This gene encodes a member of the matrix metalloproteinase family of extracellular matrix-degrading enzymes that are involved in tissue remodelling, wound repair, progression of atherosclerosis and tumour invasion. The encoded protein is activated by the removal of an N-terminal activation peptide to generate a zinc-dependent endopeptidase with a broad range of substrates such as proteoglycans, laminin, fibronectin, elastin, and collagens. |
| ***Hmgb1*** | high mobility group box 1 | This gene encodes a protein that belongs to the High Mobility Group-box superfamily. The encoded non-histone, nuclear DNA-binding protein regulates transcription, and is involved in organization of DNA. This protein plays a role in several cellular processes, including inflammation, cell differentiation and tumor cell migration. |
| ***Tlr2*** | toll-like receptor 2 | Enables lipoteichoic acid binding activity and peptide binding activity. Involved in several processes, including negative regulation of cellular component organization; positive regulation of cytokine production; and positive regulation of macromolecule biosynthetic process. Acts upstream of or within several processes, including pattern recognition receptor signaling pathway; regulation of cytokine production; and response to bacterium. Located in external side of plasma membrane. Part of Toll-like receptor 2-Toll-like receptor 6 protein complex. |
| ***Tlr3*** | toll-like receptor 3 | Predicted to enable double-stranded RNA binding activity; identical protein binding activity; and signaling receptor activity. Involved in several processes, including JNK cascade; positive regulation of angiogenesis; and positive regulation of cytokine production. Acts upstream of or within several processes, including positive regulation of NF-kappaB transcription factor activity; positive regulation |

| | | |
|---|---|---|
| | | of intracellular signal transduction; and regulation of cytokine production. |
| ***Tlr4*** | toll-like receptor 4 | This gene belongs to the Toll-like receptor family, that are involved in innate immunity. The receptor encoded by this gene mediates the innate immune response to bacterial lipopolysaccharide, a major component of the outer membrane of Gram-negative bacteria, through synthesis of pro-inflammatory cytokines and chemokines. In addition, this protein can recognize other pathogens from Gram-negative and Gram-positive bacteria as well as viral components. |
| ***Tlr9*** | toll-like receptor 9 | Predicted to enable interleukin-1 receptor binding activity; pattern recognition receptor activity; and siRNA binding activity. Involved in several processes, including positive regulation of cytokine production; regulation of defence response; and regulation of macromolecule biosynthetic process. Acts upstream of or within several processes, including cellular response to chloroquine; negative regulation of ERK1 and ERK2 cascade; and regulation of dendritic cell cytokine production. |
| ***Il1β*** | interleukin 1 beta | The protein encoded by this gene is a member of the interleukin 1 cytokine family. This cytokine is produced by activated macrophages as a proprotein, which is proteolytically processed to its active form by caspase 1. The encoded protein plays a role in thymocyte proliferation and is involved in the inflammatory response. |
| ***Casp8*** | caspase 8 | This gene is part of a family of caspases, aspartate-specific cysteine proteases well studied for their involvement in immune and apoptosis signalling. This protein, an initiator of apoptotic cell death, is activated by death-inducing tumor necrosis family receptors and targets downstream effectors. |
| ***Casp3*** | caspase 3 | This gene encodes a protein that belongs to a family of cysteinyl aspartate-specific proteases that function as essential regulators of programmed cell death through apoptosis. Members of this family contain an N-terminal pro-domain and require cleavage at specific aspartate residues to become mature. The protein encoded by this gene belongs to a subgroup of cysteinyl aspartate-specific proteases that are activated by initiator caspases and that perform the proteolytic cleavage of apoptotic target proteins. |

| | | |
|---|---|---|
| ***Tirap*** | TIR domain containing adaptor protein | Enables Toll-like receptor 2 binding activity; identical protein binding activity; and phosphatidylinositol-4,5-bisphosphate binding activity. Involved in several processes, including positive regulation of B cell proliferation; positive regulation of NF-kappaB transcription factor activity; and regulation of cytokine production. Acts upstream of or within several processes, including I-kappaB kinase/NF-kappaB signaling; myeloid cell differentiation; and response to bacterium. |
| ***Myd88*** | myeloid differentiation primary response protein 88 | Enables Toll-like receptor binding activity and interleukin-1 receptor binding activity. Involved in several processes, including cellular response to oxidised low-density lipoprotein particle stimulus; positive regulation of cytokine production; and response to bacterium. Acts upstream of or within several processes, including positive regulation of intracellular signal transduction; regulation of cytokine production; and response to bacterium. |
| ***Ticam2*** | TIR domain containing adaptor molecule 2 | Enables phospholipid binding activity. Involved in several processes, including positive regulation of cytokine production; positive regulation of interleukin-18-mediated signaling pathway; and response to interleukin-12. Acts upstream of or within defense response to virus and regulation of cytokine production. |
| ***Ticam1*** | TIR domain containing adaptor molecule 1 | Predicted to enable protein kinase binding activity. Involved in several processes, including TRIF-dependent toll-like receptor signalling pathway; cellular response to oxidised low-density lipoprotein particle stimulus; and positive regulation of cytokine production. Acts upstream of or within several processes, including positive regulation of cytokine production; positive regulation of lymphocyte activation; and positive regulation of nitrogen compound metabolic process. |
| ***Traf3*** | TNF receptor associated factor 3 | Enables protein kinase binding activity. Involved in several processes, including regulation of defence response to virus; regulation of gene expression; and tumor necrosis factor-mediated signalling pathway. Part of CD40 receptor complex. |
| ***Tbk1*** | TANK binding kinase 1 | Enables several functions, including identical protein binding activity; protein phosphatase binding activity; and protein serine/threonine kinase activity. Involved in negative regulation of gene expression; peptidyl-serine phosphorylation; and positive regulation of defense response. Acts upstream of or within several processes, including defence response |

| | | |
|---|---|---|
| | | to Gram-positive bacterium; dendritic cell proliferation; and positive regulation of interferon-beta production. |
| ***Irf3*** | interferon regulatory factor 3 | Enables several functions, including DNA-binding transcription factor activity, RNA polymerase II-specific; identical protein binding activity; and promoter-specific chromatin binding activity. Involved in several processes, including cellular response to virus; positive regulation of type I interferon production; and type I interferon signalling pathway. Acts upstream of or within several processes, including lipopolysaccharide-mediated signalling pathway; negative regulation of macromolecule metabolic process; and regulation of defence response. |
| ***Ifnb1*** | interferon beta | Enables cytokine activity and type I interferon receptor binding activity. Involved in B cell proliferation; cellular response to virus; and type I interferon signalling pathway. Acts upstream of or within several processes, including cellular response to organic cyclic compound; defence response to other organism; and negative regulation of osteoclast differentiation. |
| ***Cflar*** | CASP8 and FADD like apoptosis regulator | Enables peptidase activator activity. Involved in several processes, including negative regulation of myoblast fusion; positive regulation of NF-kappaB transcription factor activity; and skeletal muscle atrophy. Acts upstream of or within negative regulation of programmed cell death and response to bacterium. |
| ***Prkcg*** | protein kinase C gamma | Predicted to enable calcium-dependent protein kinase C activity and protein serine/threonine/tyrosine kinase activity. Involved in several processes, including modulation of chemical synaptic transmission; negative regulation of cellular protein metabolic process; and response to morphine. Acts upstream of or within chemosensory behaviour and regulation of phagocytosis. |
| ***Irak*** | interleukin 1 receptor associated kinase | Enables interleukin-1 receptor binding activity and protein kinase activity. Involved in negative regulation of cholesterol efflux; negative regulation of transcription, DNA-templated; and positive regulation of JUN kinase activity. Acts upstream of or within several processes, including interleukin-1-mediated signalling pathway; response to molecule of bacterial origin; and toll-like receptor signalling pathway. |

| | | |
|---|---|---|
| ***Traf6*** | TNF receptor associated factor 6 | This gene encodes a member of the TNF receptor associated factor (TRAF) family of adaptor proteins that mediate signalling events from members of the TNF receptor and Toll/IL-1 receptor families to activate transcription factors such as NF-kappa-B and AP-1. The product of this gene is essential for perinatal and postnatal survival. |
| ***Irf7*** | interferon regulatory factor 7 | Enables DNA-binding transcription factor activity, RNA polymerase II-specific and cis-regulatory region sequence-specific DNA binding activity. Involved in defence response to other organism; positive regulation of transcription, DNA-templated; and regulation of toll-like receptor signalling pathway. Acts upstream of or within immunoglobulin mediated immune response; positive regulation of type I interferon production; and positive regulation of type I interferon-mediated signalling pathway. |
| ***Bdnf*** | brain-derived neurotrophic factor | The protein encoded by this gene is a member of the nerve growth factor family. It is involved in the growth, differentiation and survival of specific types of developing neurons both in the central nervous system (CNS) and the peripheral nervous system. It is also involved in regulating synaptic plasticity in the CNS. |
| ***Ngf*** | nerve growth factor | Enables transmembrane receptor protein tyrosine kinase activator activity. Involved in positive regulation of DNA binding activity; positive regulation of Ras protein signal transduction; and positive regulation of protein phosphorylation. Acts upstream of or within several processes, including cell surface receptor signalling pathway; positive regulation of cell projection organization; and positive regulation of macromolecule metabolic process. |
| ***Il1r1*** | interleukin 1 receptor type 1 | Enables interleukin-1 binding activity; interleukin-1 receptor activity; and protease binding activity. Involved in interleukin-1-mediated signalling pathway. Acts upstream of or within several processes, including cytokine-mediated signaling pathway; positive regulation of interleukin-1-mediated signaling pathway; and positive regulation of neutrophil extravasation. |
| ***Ccl5*** | C-C motif chemokine ligand 5 | Enables CCR1 chemokine receptor binding activity. Involved in negative regulation of macrophage apoptotic process and positive regulation of monocyte chemotaxis. Acts upstream of or within inflammatory response; |

| | | |
|---|---|---|
| | | positive regulation of epithelial cell proliferation; and response to tumor necrosis factor. |
| ***Ccll2*** | C-C motif chemokine ligand 2 | This chemokine is a member of the CC subfamily which is characterized by two adjacent cysteine residues. This cytokine displays chemotactic activity for monocytes and memory T cells but not for neutrophils. |
| ***Gad*** | glutamate decarboxylase | Enables thiol-dependent deubiquitinase and ubiquitin binding activity. Involved in cellular response to xenobiotic stimulus. Acts upstream of or within several processes, including adult walking behaviour; axon target recognition; and response to ischemia. |
| ***Tnf*** | tumour necrosis factor | This gene encodes a multifunctional proinflammatory cytokine that belongs to the tumor necrosis factor (TNF) superfamily. It plays an important role in the innate immune response as well as regulating homeostasis but is also implicated in diseases of chronic inflammation. |
| ***Il18*** | interleukin 18 | Enables cytokine activity. Involved in lipid homeostasis; positive regulation of cell differentiation; and positive regulation of cold-induced thermogenesis. Acts upstream of or within several processes, including interleukin-18-mediated signalling pathway; positive regulation of NIK/NF-kappaB signalling; and positive regulation of interferon-gamma production. |
| ***Il12*** | interleukin 12 | Contributes to cytokine activity. Acts upstream of or within several processes, including T-helper 1 cell activation; defense response to protozoan; and positive regulation of T cell activation. |
| ***Il10*** | interleukin 10 | This gene encodes an anti-inflammatory cytokine that is a member of the class-2 cytokine family. The encoded protein is secreted by cells of both the innate and adaptive immune systems and is crucial for limiting the immune response to a broad range of pathogens. It also has been shown to suppress autoimmune responses. This protein mediates its immunosuppressive signal through a specific interleukin 10 receptor complex. |
| ***Crp*** | C-reactive protein | Predicted to enable several functions, including cholesterol binding activity; complement component C1q complex binding activity; and low-density lipoprotein particle binding activity. Predicted to be involved in several processes, including complement activation, classical pathway; regulation of gene expression; and |

| | | |
|---|---|---|
| | | regulation of superoxide anion generation. Predicted to act upstream of or within vasoconstriction. |
| ***Il4*** | interleukin 4 | Enables cytokine activity. Involved in several processes, including innate immune response in mucosa; negative regulation of white fat cell proliferation; and regulation of gene expression. Acts upstream of or within several processes, including T-helper cell differentiation; positive regulation of macromolecule metabolic process; and regulation of leukocyte activation. |
| ***Mapt*** | microtubule associated protein tau | Enables DNA binding activity; microtubule binding activity; and protein kinase binding activity. Involved in negative regulation of tubulin deacetylation; regulation of cellular response to heat; and regulation of response to DNA damage stimulus. Acts upstream of or within several processes, including adult walking behavior; generation of neurons; and transport along microtubule. |
| ***Calb*** | calbindin | Enables calcium ion binding activity involved in regulation of postsynaptic cytosolic calcium ion concentration and calcium ion binding activity involved in regulation of presynaptic cytosolic calcium ion concentration. Involved in regulation of long-term synaptic potentiation. Acts upstream of or within several processes, including locomotory behavior; metanephros development; and retina layer formation. |
| ***Sod2*** | superoxide dismutase 2 | Enables superoxide dismutase activity. Acts upstream of or within several processes, including animal organ development; apoptotic signaling pathway; and regulation of cellular biosynthetic process. |
| ***Bcl2*** | apoptosis regulator BCL-2 | This gene encodes a member of the B cell lymphoma 2 protein family. Members of this family regulate cell death in multiple cell types and can have either proapoptotic or antiapoptotic activities. The protein encoded by this gene inhibits mitochondrial-mediated apoptosis. This protein is an integral outer mitochondrial membrane protein that functions as part of signalling pathway that controls mitochondrial permeability in response to apoptotic stimuli. This protein may also play a role in neuron cell survival and autophagy. |
| ***Gdnf*** | glial cell derived neurotrophic factor | Ligands of this family bind various TGF-beta receptors leading to recruitment and activation of SMAD family transcription factors that regulate gene expression. The encoded preproprotein is proteolytically processed to |

| | | |
|---|---|---|
| | | generate each subunit of the disulfide-linked homodimer. The recombinant form of this protein, a highly conserved neurotrophic factor, was shown to promote the survival and differentiation of dopaminergic neurons in culture and was able to prevent apoptosis of motor neurons induced by axotomy. This protein is a ligand for the product of the RET (rearranged during transfection) protooncogene. |
| ***Iap*** | Not available | Not available. |
| ***Cd80*** | CD80 | Predicted to enable coreceptor activity. Acts upstream of or within T cell costimulation; cellular response to lipopolysaccharide; and positive regulation of alpha-beta T cell proliferation. |
| ***Cd86*** | CD86 | Predicted to enable signaling receptor binding activity. Involved in several processes, including CD40 signaling pathway; activation of phospholipase C activity; and positive regulation of macromolecule metabolic process. Acts upstream of or within several processes, including cellular response to lipopolysaccharide; defense response to virus; and positive regulation of T cell proliferation. |
| ***Cd40*** | CD40 | Predicted to enable antigen binding activity; protein domain specific binding activity; and ubiquitin protein ligase binding activity. Involved in B cell mediated immunity; CD40 signalling pathway; and cellular calcium ion homeostasis. Acts upstream of or within several processes, including defence response to other organism; positive regulation of B cell activation; and positive regulation of interleukin-12 production. |
| ***Ntf3*** | neurotrophin 3 | This gene encodes a member of the neurotrophins that have a wide variety of functions in both neural and non-neural tissues. The encoded preproprotein undergoes proteolytic processing to generate a noncovalently linked homodimeric mature protein that can bind to the transmembrane receptor tyrosine kinases to initiate a series of signalling events. |
| ***Pycard*** | PYD and CARD domain containing | Enables identical protein binding activity and protein dimerization activity. Involved in several processes, including activation of cysteine-type endopeptidase activity; positive regulation of cytokine production; and regulation of defence response. Acts upstream of or within several processes, including defence response to Gram-positive bacterium; positive regulation of |

| | | |
|---|---|---|
| | | macromolecule metabolic process; and regulation of autophagy. |
| ***Casp1*** | caspase 1 | Enables cysteine-type endopeptidase activity. Involved in several processes, including positive regulation of I-kappaB kinase/NF-kappaB signalling; positive regulation of interleukin-1 beta production; and protein auto processing. Acts upstream of or within several processes, including membrane hyperpolarization; mitochondrial depolarization; and positive regulation of interleukin-1 alpha production. |
| ***Nlrp3*** | NACHT, LRR and PYD domains-containing protein 3 | Enables DNA-binding transcription factor binding activity and sequence-specific DNA binding activity. Involved in several processes, including positive regulation of T-helper cell differentiation; positive regulation of cytokine production; and response to bacterium. Acts upstream of or within several processes, including NLRP3 inflammasome complex assembly; activation of cysteine-type endopeptidase activity involved in apoptotic process; and defence response to virus. |
| ***S100b*** | S100 calcium binding protein B | Predicted to enable several functions, including RAGE receptor binding activity; S100 protein binding activity; and metal ion binding activity. Involved in learning or memory. Acts upstream of or within memory and regulation of neuronal synaptic plasticity. |
| ***Wnt1*** | wingless-type MMTV integration site family, member 1 | Enables cytokine activity and protein domain specific binding activity. Involved in several processes, including astrocyte-dopaminergic neuron signalling; central nervous system development; and regulation of transcription, DNA-templated. Acts upstream of or within several processes, including animal organ development; regulation of cellular protein metabolic process; and ubiquitin-dependent SMAD protein catabolic process. |
| ***Tnfrsf1a*** | TNF receptor superfamily member 1A | This gene encodes a member of the TNF receptor superfamily of proteins. The encoded receptor is found in membrane-bound and soluble forms that interact with membrane-bound and soluble forms, respectively, of its ligand, tumor necrosis factor alpha. Binding of membrane-bound tumor necrosis factor alpha to the membrane-bound receptor induces receptor trimerization and activation, which plays a role in cell survival, apoptosis, and inflammation. Proteolytic processing of the encoded receptor results in release of the soluble form of the receptor, which can interact |

| | | |
|---|---|---|
| | | with free tumor necrosis factor alpha to inhibit inflammation. |
| ***Fasf*** | Not available | Not available. |
| ***Aslg*** | Not available | Not available. |
| ***Fzd1*** | frizzled class receptor 1 | Enables Wnt-protein binding activity. Involved in astrocyte-dopaminergic neuron signalling; negative regulation of oxidative stress-induced neuron death; and regulation of presynapse assembly. Acts upstream of or within several processes, including Wnt signalling pathway; heart morphogenesis; and negative regulation of signal transduction. |
| ***P2rx7*** | purinergic receptor P2X 7 | Enables several functions, including ATP binding activity; extracellularly ATP-gated cation channel activity; and lipopolysaccharide binding activity. Involved in negative regulation of cell volume and sensory perception of pain. Acts upstream of or within several processes, including organophosphate ester transport; positive regulation of cytokine production; and positive regulation of secretion. |
| ***Snca*** | synuclein alpha | Enables arachidonic acid binding activity; histone binding activity; and identical protein binding activity. Involved in several processes, including positive regulation of peptidyl-serine phosphorylation; protein-containing complex assembly; and synaptic vesicle endocytosis. Acts upstream of or within several processes, including chemical synaptic transmission; modulation of chemical synaptic transmission; and regulation of amine transport. |
| ***Gsk3b*** | glycogen synthase kinase 3 β | Enables several functions, including beta-catenin binding activity; dynein complex binding activity; and protein kinase activity. Involved in several processes, including apoptotic signalling pathway; negative regulation of signal transduction; and regulation of gene expression. Acts upstream of or within several processes, including cellular response to hepatocyte growth factor stimulus; positive regulation of nitrogen compound metabolic process; and regulation of neuron projection development. |
| ***Gls*** | glutaminase | Enables glutaminase activity and identical protein binding activity. Involved in glutamate biosynthetic process; glutamine catabolic process; and protein homotetramerization. Acts upstream of or within chemical synaptic transmission; regulation of respiratory gaseous exchange by nervous system process; and suckling behaviour. |

| | | |
|---|---|---|
| ***Ctnnb1*** | catenin beta 1 | This gene encodes not only an important cytoplasmic component of the classical cadherin adhesion complex that forms the adherens junction in epithelia and mediates cell-cell adhesion in many other tissues but also a key signalling molecule in the canonical Wnt signalling pathway that controls cell growth and differentiation during both normal development and tumorigenesis. Beta-catenin is therefore necessary for the adhesive function of classical cadherins. |
| ***Bace1*** | beta-secretase 1 | This gene encodes a member of the peptidase A1 family of aspartic proteases. This transmembrane protease catalyzes the first step in the formation of amyloid beta peptide from amyloid precursor protein. |
| ***Icam1*** | intercellular adhesion molecule 1 | This gene encodes an integral membrane protein that binds leukocyte adhesion protein LFA-1. It participates in the innate immune response. |
| ***Vcam1*** | vascular cell adhesion molecule 1 | Predicted to enable integrin binding activity and primary amine oxidase activity. Acts upstream of or within several processes, including cellular response to glucose stimulus; chorio-allantoic fusion; and heterophilic cell-cell adhesion via plasma membrane cell adhesion molecules. |
| ***Cntf*** | ciliary neurotrophic factor | The protein encoded by this gene is a polypeptide hormone whose actions appear to be restricted to the nervous system where it promotes neurotransmitter synthesis and neurite outgrowth in certain neuronal populations. The protein is a potent survival factor for neurons and oligodendrocytes, and it may be involved in reducing tissue destruction during inflammatory attacks. |
| ***Glul*** | glutamate-ammonia ligase | Already described in Supplementary Table 2. |
| ***Slc1a2*** | solute carrier family 1 member 2 | Enables cysteine transmembrane transporter activity and high-affinity glutamate transmembrane transporter activity. Involved in several processes, including L-glutamate import across plasma membrane; cellular response to cocaine; and glutathione biosynthetic process. Acts upstream of or within several processes, including L-glutamate transmembrane transport; positive regulation of glucose import; and visual behaviour. |
| ***Slc1a3*** | solute carrier family 1 member 3 | Enables glutamate binding activity and high-affinity glutamate transmembrane transporter activity. Involved in D-aspartate import across plasma membrane and cellular response to cocaine. Acts upstream of or within several |

| | | |
|---|---|---|
| | | processes, including L-glutamate import across plasma membrane; gamma-aminobutyric acid biosynthetic process; and nervous system development. |

**Supplementary Table 7. *Bmal1* as master regulator exhibits different gene targets regulated based on *Bmal1* inhibition levels across different ZTs.** The identified genes' names and biological function in mice (*Mus musculus*) were determined based on the NCBI National Center for Biotechnology Information (https://www.ncbi.nlm.nih.gov/gene) tool.

| Code | Name | Biological function |
|---|---|---|
| ***Acox1*** | acyl-CoA oxidase 1 | This gene encodes a member of the acyl-coenzyme A oxidase family. The encoded protein is localized to peroxisomes and is the first enzyme of the fatty acid beta-oxidation pathway, which catalyzes the desaturation of acyl-coenzyme A to 2-trans-enoyl-coenzyme A. |
| ***Angpt1*** | angiopoietin 1 | This gene encodes a secreted glycoprotein that belongs to the angiopoietin family of vascular growth factors. The encoded protein is a ligand in the vascular tyrosine kinase signaling pathway and regulates the formation and stabilization of blood vessels. It also plays an essential role in vascular response to tissue injury. |
| ***Ccl2*** | C-C motif chemokine ligand 2 | Chemokines are a superfamily of secreted proteins involved in immunoregulatory and inflammatory processes. This chemokine is a member of the CC subfamily which is characterized by two adjacent cysteine residues. This cytokine displays chemotactic activity for monocytes and memory T cells but not for neutrophils. |
| ***Ccng2*** | cyclin G2 | Predicted to enable cyclin-dependent protein serine/threonine kinase regulator activity. Acts upstream of or within regulation of cell cycle. Predicted to be located in cytosol. Predicted to be part of cyclin-dependent protein kinase holoenzyme complex. |
| ***Cebpa*** | CCAAT enhancer binding protein alpha | Activity of this protein can modulate the expression of genes involved in cell cycle regulation as well as in body weight homeostasis. |
| ***Cry1*** | cryptochrome circadian regulator 1 | This gene encodes a flavin adenine dinucleotide-binding protein, a key component of the circadian core oscillator complex that regulates the circadian clock. This gene is upregulated by CLOCK/ARNTL heterodimers but then represses this upregulation in a feedback loop using PER/CRY heterodimers to interact with CLOCK/ARNTL. |

| | | |
|---|---|---|
| ***Dbp*** | D-Box binding PAR BZIP transcription factor | The protein encoded by this gene is a member of the PAR bZIP transcription factor family and binds to specific sequences in the promoters of several genes, such as albumin, CYP2A4, and CYP2A5. The encoded protein can bind in the DNA as a homo- or heterodimer and is involved in the regulation of some circadian rhythm genes. |
| ***Egln3*** | egl-9 family hypoxia inducible factor 3 | Enables peptidyl-proline dioxygenase activity. Involved in regulation of cell population proliferation and regulation of neuron apoptotic process. Acts upstream of or within cellular response to leukaemia inhibitory factor. |
| ***Fasn*** | fatty acid synthase | Enables fatty acid synthase activity. Involved in ether lipid biosynthetic process; mammary gland development; and myeloid leukocyte differentiation. Acts upstream of or within cellular response to interleukin-4; epithelial cell development; and fatty acid biosynthetic process. |
| ***Gpam*** | glycerol-3-phosphate acyltransferase mitochondrial | Enables glycerol-3-phosphate O-acyltransferase activity. Acts upstream of or within several processes, including glycerolipid metabolic process; lipid homeostasis; and negative regulation of activation-induced cell death of T cells. |
| ***Gsr*** | glutathione-disulfide reductase | Enables phosphatidylinositol 3-kinase catalytic subunit binding activity. Involved in several processes, including actin filament capping; actin polymerization or depolymerization; and positive regulation of protein processing in phagocytic vesicle. Acts upstream of or within cellular response to interferon-gamma and vesicle-mediated transport. |
| ***Hif1a*** | hypoxia inducible factor 1 subunit alpha | This gene encodes the alpha subunit which, along with the beta subunit, forms a heterodimeric transcription factor that regulates the cellular and developmental response to reduced oxygen tension. The transcription factor has been shown to regulate genes involved in several biological processes, including erythropoiesis and angiogenesis which aid in increased delivery of oxygen to hypoxic regions. It also plays a role in the induction of genes involved in cell proliferation and survival, energy metabolism, apoptosis, and glucose and iron metabolism. |
| ***Hk2*** | hexokinase 2 | Enables hexokinase activity. Involved in negative regulation of mitochondrial membrane permeability and negative regulation of reactive oxygen species metabolic process. Acts upstream of or within several processes, |

| | | |
|---|---|---|
| | | including carbohydrate phosphorylation; cellular response to leukemia inhibitory factor; and regulation of glucose import. |
| ***Hlf*** | HLF transcription factor, PAR BZIP family member | Predicted to enable DNA-binding transcription activator activity, RNA polymerase II-specific and RNA polymerase II cis-regulatory region sequence-specific DNA binding activity. Acts upstream of or within skeletal muscle cell differentiation. |
| ***Hmgcs2*** | 3-hydroxy-3-methylglutaryl-CoA synthase 2 | Enables hydroxymethylglutaryl-CoA synthase activity. Predicted to be involved in acetyl-CoA metabolic process; farnesyl diphosphate biosynthetic process, mevalonate pathway; and ketone body biosynthetic process. Predicted to act upstream of or within cholesterol biosynthetic process. |
| ***Hmox1*** | heme oxygenase 1 | Predicted to enable several functions, including heme binding activity; heme oxygenase (decyclizing) activity; and protein homodimerization activity. Acts upstream of or within several processes, including cellular response to cisplatin; cellular response to metal ion; and regulation of macroautophagy. |
| ***Hspa5*** | heat shock protein family A (Hsp70) member 5 | Enables misfolded protein binding activity and ribosome binding activity. Involved in several processes, including IRE1-mediated unfolded protein response; cerebellum development; and positive regulation of protein ubiquitination. Acts upstream of or within several processes, including cellular response to interleukin-4; negative regulation of transforming growth factor beta receptor signaling pathway; and toxin transport. |
| ***Il1β*** | interleukin 1β | The protein encoded by this gene is a member of the interleukin 1 cytokine family. This cytokine is produced by activated macrophages as a proprotein, which is proteolytically processed to its active form by caspase 1. The encoded protein plays a role in thymocyte proliferation and is involved in the inflammatory response. |
| ***Itgb8*** | integrin subunit β8 | Predicted to enable extracellular matrix protein binding activity and integrin binding activity. Involved in regulation of transforming growth factor beta activation. Acts upstream of or within several processes, including Langerhans cell differentiation; ganglioside metabolic process; and hard palate development. Part of integrin alphav-beta8 complex. |
| ***Lamp2*** | lysosomal-associated membrane protein 2 | Enables protein domain specific binding activity. Involved in several processes, including autophagosome maturation; protein |

| | | |
|---|---|---|
| | | stabilization; and protein targeting to lysosome involved in chaperone-mediated autophagy. Acts upstream of or within muscle cell cellular homeostasis. Located in late endosome membrane; lysosomal membrane; and phagocytic vesicle membrane. Is integral component of autophagosome membrane. |
| ***Ldlr*** | low-density lipoprotein receptor | Enables several functions, including amyloid-beta binding activity; low-density lipoprotein particle binding activity; and low-density lipoprotein particle receptor activity. Involved in several processes, including lipid transport; regulation of inflammatory response; and regulation of lipid metabolic process. Acts upstream of or within several processes, including cholesterol homeostasis; lipoprotein catabolic process; and low-density lipoprotein particle clearance. |
| ***Mcm6*** | minichromosome maintenance complex component 6 | Enables single-stranded DNA binding activity. Contributes to DNA helicase activity. Acts upstream of or within DNA unwinding involved in DNA replication. |
| ***Nfe2l2*** | NFE2 Like BZIP transcription factor 2 | This gene encodes a transcription factor which is a member of a small family of basic leucine zipper (bZIP) proteins. The encoded transcription factor regulates genes which contain antioxidant response elements (ARE) in their promoters; many of these genes encode proteins involved in response to injury and inflammation which includes the production of free radicals. |
| ***Nfil3*** | nuclear factor, interleukin 3 regulated | The protein encoded by this gene is a transcriptional regulator that binds as a homodimer to activate transcription factor sites in many cellular and viral promoters. The encoded protein represses PER1 and PER2 expression and therefore plays a role in the regulation of circadian rhythm. |
| ***Npy*** | neuropeptide Y | This gene encodes a neuropeptide that plays a pivotal role in many physiological functions such as food intake, energy homeostasis, circadian rhythm, and cognition. The encoded protein precursor undergoes proteolytic processing to generate the biologically active peptide. |
| ***Nqo1*** | NAD(P)H quinone dehydrogenase 1 | Enables NAD(P)H dehydrogenase (quinone) activity. Acts upstream of or within negative regulation of catalytic activity and response to oxidative stress. Predicted to be located in cytoplasm; dendrite; and neuronal cell body. |
| ***Nr1d1*** | nuclear receptor subfamily 1 group D member 1 | This gene encodes a transcription factor that is a member of the nuclear receptor subfamily 1. |

| | | |
|---|---|---|
| | | The encoded protein is a ligand-sensitive transcription factor that negatively regulates the expression of core clock proteins. In particular, this protein represses the circadian clock transcription factor aryl hydrocarbon receptor nuclear translocator-like protein 1 (ARNTL). This protein may also be involved in regulating genes that function in metabolic, inflammatory, and cardiovascular processes. |
| ***Parvb*** | parvin β | Predicted to enable actin binding activity. Predicted to be involved in several processes, including actin cytoskeleton reorganization; lamellipodium assembly; and substrate adhesion-dependent cell spreading. |
| ***Pdp1*** | pyruvate dehydrogenase phosphatase catalytic subunit 1 | Predicted to enable metal ion binding activity and phosphoprotein phosphatase activity. Predicted to be involved in peptidyl-threonine dephosphorylation and positive regulation of pyruvate dehydrogenase activity. |
| ***Per1*** | period circadian regulator 1 | This gene is a member of the Period family of genes and is expressed in a circadian pattern in the suprachiasmatic nucleus, the primary circadian pacemaker in the mammalian brain. Genes in this family encode components of the circadian rhythms of locomotor activity, metabolism, and behavior. This gene is upregulated by CLOCK/ARNTL heterodimers but then represses this upregulation in a feedback loop using PER/CRY heterodimers to interact with CLOCK/ARNTL. Polymorphisms in this gene may increase the risk of getting certain cancers. |
| ***Per2*** | period circadian regulator 2 | This gene is a member of the Period family of genes and is expressed in a circadian pattern in the suprachiasmatic nucleus, the primary circadian pacemaker in the mammalian brain. Genes in this family encode components of the circadian rhythms of locomotor activity, metabolism, and behavior. This gene is upregulated by CLOCK/ARNTL heterodimers but then represses this upregulation in a feedback loop using PER/CRY heterodimers to interact with CLOCK/ARNTL. |
| ***Per3*** | period circadian regulator 3 | This gene is a member of the Period family of genes and is expressed in a circadian pattern in the suprachiasmatic nucleus, the primary circadian pacemaker in the mammalian brain. Genes in this family encode components of the circadian rhythms of locomotor activity, metabolism, and behavior. This gene is upregulated by CLOCK/ARNTL heterodimers but then represses this upregulation in a |

| | | |
|---|---|---|
| | | feedback loop using PER/CRY heterodimers to interact with CLOCK/ARNTL. |
| ***Rara*** | retinoic acid receptor alpha | Enables RNA polymerase II transcription regulatory region sequence-specific DNA binding activity and nuclear receptor activity. Involved in several processes, including Sertoli cell fate commitment; cellular response to lipopolysaccharide; and germ cell development. Acts upstream of or within several processes, including animal organ development; negative regulation of cartilage development; and regulation of cellular macromolecule biosynthetic process. |
| ***Ryr2*** | ryanodine receptor 2 | Enables several functions, including calmodulin binding activity; identical protein binding activity; and ryanodine-sensitive calcium-release channel activity. Involved in several processes, including cellular response to organonitrogen compound; regulation of heart contraction; and release of sequestered calcium ion into cytosol by sarcoplasmic reticulum. Acts upstream of or within several processes, including calcium ion transport; left ventricular cardiac muscle tissue morphogenesis; and response to caffeine. |
| ***Scd*** | Stearoyl-CoA desaturase | Enables metal ion binding activity; palmitoyl-CoA 9-desaturase activity; and stearoyl-CoA 9-desaturase activity. Involved in several processes, including cholesterol esterification; positive regulation of cold-induced thermogenesis; and tarsal gland development. Acts upstream of or within several processes, including brown fat cell differentiation; monounsaturated fatty acid biosynthetic process; and white fat cell differentiation. |
| ***Srebf1*** | sterol regulatory element binding transcription factor 1 | This gene encodes a transcription factor that binds to the sterol regulatory element-1 (SRE1), which is a decamer flanking the low density lipoprotein receptor gene and some genes involved in sterol biosynthesis. Following cleavage, the mature protein translocates to the nucleus and activates transcription by binding to the SRE1. Sterols inhibit the cleavage of the precursor, and the mature nuclear form is rapidly catabolized, thereby reducing transcription. The protein is a member of the basic helix-loop-helix-leucine zipper (bHLH-Zip) transcription factor family. |
| ***Stard4*** | StAR related lipid transfer domain containing 4 | Predicted to enable cholesterol binding activity and cholesterol transfer activity. Involved in cholesterol esterification. |
| ***Thra*** | thyroid hormone receptor alpha | The protein encoded by this gene is one of several nuclear hormone receptors that bind |

| | | |
|---|---|---|
| | | thyroid hormones such as triiodothyronine and thyroxine with high affinity. The encoded protein is a transcription factor that can activate or repress transcription. |
| ***Tp53*** | tumour protein P53 | This gene encodes tumor protein p53, which responds to diverse cellular stresses to regulate target genes that induce cell cycle arrest, apoptosis, senescence, DNA repair, or changes in metabolism. p53 protein is expressed at low level in normal cells and at a high level in a variety of transformed cell lines, where it's believed to contribute to transformation and malignancy. p53 is a DNA-binding protein containing transcription activation, DNA-binding, and oligomerization domains. |
| ***Ttk*** | TTK protein kinase | Enables identical protein binding activity; kinetochore binding activity; and protein serine/threonine/tyrosine kinase activity. Involved in mitotic spindle assembly checkpoint signaling. Acts upstream of or within several processes, including meiotic spindle assembly checkpoint signaling; protein localization to organelle; and protein phosphorylation. |
| ***Wee1*** | WEE1 G2 checkpoint kinase | Enables protein kinase activity. Acts upstream of or within several processes, including establishment of cell polarity; neuron projection morphogenesis; and peptidyl-tyrosine phosphorylation. |

**Supplementary Table 8.** Demographic and clinical data collected from MTLE+HS and control participants. CTRL, control; F, female; HS, hippocampal sclerosis; M, male; MTLE, mesial temporal lobe epilepsy; NA, not Applicable; ND, not determined; NC, not classified; ys, years of disease.

| Condition | Age | Sex | Age onset | Disease (ys) | HS lateralitty | HS classification |
|---|---|---|---|---|---|---|
| Case | 38 | F | 13 | 25 | Left | 1 |
| Case | 23 | F | 6 | 17 | Left | 1 |
| Case | 46 | M | 29 | 17 | Right | 1 |
| Case | 32 | F | 1,5 | 30,5 | Left | 1 |
| Control | 33 | M | NA | NA | ND | NA |
| Case | 38 | F | 6 | 32 | Right | 1 |
| Case | 38 | M | 13 | 25 | Left | 1 |
| Control | 54 | M | NA | NA | ND | NA |
| Case | 61 | F | 15 | 46 | Right | 1 |
| Control | 66 | F | NA | NA | ND | NA |
| Control | 66 | F | NA | NA | ND | NA |
| Case | 47 | F | 27 | 20 | Left | 1 |
| Control | 55 | F | NA | NA | ND | NA |
| Case | 41 | F | 18 | 23 | Right | 2 |

| | | | | | | |
|---|---|---|---|---|---|---|
| Case | 39 | M | 0 | 39 | Left | 1 |
| Case | 60 | M | 14 | 46 | Left | 1 |
| Case | 54 | F | 2 | 52 | Right | 1 |
| Case | 44 | F | 1 | 43 | Right | 2 |
| Case | 43 | M | 3 | 40 | Left | NC |
| Case | 49 | M | 9 | 40 | Left | 1 |
| Case | 38 | F | 14 | 24 | Right | 1 |
| Case | 40 | F | 36 | 4 | Left | 1 |
| Control | 56 | F | NA | NA | ND | NA |

**Supplementary Table 9.** List of the genes and primers sequences used in the qPCR analysis in this study.

| Gene | Primer sequence (5' to 3') |
|---|---|
| ***Gfap*** | Forward: CCCTGAGGCAGAAGCTCCAA<br>Reverse: GAGCCAGGGTGGCTTCATCT |
| ***C4b*** | Forward: CAACAAGGGAGACCCCCAGT<br>Reverse: TAAGGCCTCACACCTGGCAC |
| ***Serpina3n*** | Forward: CTCCACCGACTACAGCCTGG<br>Reverse: CAGCCTTGTGGACCACCTGA |
| ***Glul*** | Forward: CGGAAACCTGCAGAGACCAAC<br>Reverse: CAACCAAATGGGTGGCCGTC |
| ***Bmal1*** | Forward: TGCAATGTCCAGGAAGTTAGA<br>Reverse: GTTTGCTTCTGTGTATGGGTT |
